# Low hybridization temperatures improve target capture success of invertebrate loci

**DOI:** 10.1101/2022.03.02.482542

**Authors:** Michael Forthman, Eric R. L. Gordon, Rebecca T. Kimball

**Affiliations:** California State Collection of Arthropods, Plant Pest Diagnostics Branch, California Department of Food and Agriculture, 3294 Meadowview Road, Sacramento, CA 95832, USA; Entomology & Nematology Department, University of Florida, 1881 Natural Area Drive, Gainesville, FL 32611, USA; Department of Ecology & Evolutionary Biology, University of Connecticut, 75 N. Eagleville Road, Unit 3043, Storrs, CT 06269, USA; Department of Biology, University of Florida, 876 Newell Drive, Gainesville, FL 32611, USA

**Keywords:** Annealing temperature, exons, Pentatomomorpha, touchdown hybridization, ultraconserved elements

## Abstract

Target capture approaches are widely used in phylogenomic studies, yet only four experimental comparisons of a critical parameter, hybridization temperature, have been published. These studies provide conflicting conclusions regarding the benefits of lower temperatures during target capture, and none include invertebrates where bait-target divergences may be higher than seen in vertebrate capture studies. Most capture studies use a fixed hybridization temperature of 65°C to maximize the proportion of on-target data, but many invertebrate capture studies report low locus recovery. Lower hybridization temperatures, which might improve locus recovery, are not commonly employed in invertebrate capture studies. We used leaf-footed bugs and relatives (Hemiptera: Coreoidea) to investigate the effect of hybridization temperature on capture success of ultraconserved elements (UCE) targeted by previously published baits derived from divergent hemipteran genomes and other loci targeted by newly designed baits derived from less divergent coreoid transcriptomes. We found touchdown capture approaches with lower hybridization temperatures generally resulted in lower proportions of on-target reads and lower read depth but were associated with more contigs and improved recovery of UCE loci. Low temperatures were also associated with increased numbers of putative paralogs of UCE loci. Hybridization temperatures did not generally affect recovery of newly targeted loci, which we attributed to their lower bait-target divergences (compared to higher divergences between UCE baits and targets) and greater bait tiling density. Thus, optimizing *in vitro* target capture conditions to accommodate low hybridization temperatures can provide a cost-effective, widely applicable solution to improve recovery of protein-coding loci in invertebrates.

## Introduction

Many biological disciplines have witnessed a dramatic increase in the amount of genomic data sampled due to recent advances and declining cost of next-generation sequencing technologies. While low-coverage, whole genome sequencing may be cost-effective for some research questions, genome reduction approaches that enrich for genomic regions of interest prior to high-throughput sequencing are often more feasible (reviewed in Lemmon & Lemmon, 2013). One of the most frequently employed genome reduction techniques in phylogenomic studies are target capture approaches, which include exon capture, anchored hybrid enrichment (AHE), and capture of ultraconserved elements (UCEs) (e.g., Bi et al., 2012; Faircloth et al., 2012; Lemmon et al., 2012; Li et al., 2013). In general, target capture leverages existing genomic resources to synthesize short (60–120 bp) nucleotide bait sequences complementary to genomic regions of interest. Baits are then hybridized to DNA libraries, and unbound DNA (i.e., non- or off-targets) is removed via a series of washing steps prior to sequencing.

Target capture approaches have been widely applied to a diversity of taxa at various evolutionary timescales, e.g., birds (Faircloth et al., 2012), lizards (Bragg et al., 2016), fish (Dornburg et al., 2017), plants (Fragoso-Martínez et al., 2017), spiders (Hamilton et al., 2016), and insects (Faircloth et al., 2015). While vertebrate capture studies often recover a relatively high proportion (>50%) of targeted loci (e.g., Faircloth et al., 2012; Crawford et al., 2012; Li et al., 2013), many invertebrate studies suffer from a much lower yield, particularly when highly conserved loci like UCEs are targeted (e.g., Faircloth et al., 2015; Hamilton et al., 2016; Baca et al., 2017; Van Dam et al., 2017; Dietrich et al., 2017; Kieran et al., 2019). While these and other studies may vary in the conditions of the capture experiment, the often-consistent disparity in the proportion of loci recovered between vertebrate and invertebrate target capture experiments suggests that capture dynamics may differ due to greater bait-target divergences among invertebrate taxa compared to many vertebrates (Makunin et al., 2013; Gustafson et al., 2019; Van Dam et al., 2021).

Optimizing existing invertebrate target capture bait sets to be more tailored to focal taxa has been shown to improve recovery (e.g., Branstetter et al., 2017; Gustafson et al., 2020), suggesting that baits may often be too divergent from some taxa to allow effective recovery. However, genomic resources that permit such optimization for many other groups are still lacking. As such, optimizing one or more *in vitro* target capture conditions may provide a more cost-effective solution to improve locus recovery. Recent studies in various non-invertebrate taxa have investigated several parameters that may affect the success of *in vitro* target capture approaches, including, e.g., GC content and tiling of baits, starting amount of DNA or baits, bait-target divergence, and washing stringency (e.g., Ávila-Arcos et al., 2011; Li et al., 2013; Cruz-Dávalos et al., 2017). Another condition that can affect capture success is hybridization temperature during bait-target annealing.

Invertebrate target capture studies typically employ a fixed hybridization temperature at 65°C as suggested by standard protocols (we note that temperatures are often not reported in AHE studies). However, lower hybridization temperatures, whether fixed or achieved through touchdown, may improve on-target and locus recovery due to relaxed specificity and increased sensitivity between baits and targets. This may be particularly advantageous if some baits are more divergent from their targets (as is commonly the case in invertebrates) or have lower optimal annealing temperatures than other baits. However, relaxing specificity to allow for partial matching between baits and targets should also increase the risk of baits to potentially hybridize with paralogous sequences exhibiting some degree of divergence from the corresponding target sequence and/or may increase the number of off-target sequences (e.g., Cruz-Dávalos et al., 2017) and thus reduce read numbers for targeted regions. While a few invertebrate capture studies have used lower temperatures during bait-target hybridization (Zhang et al., 2019; Braby et al., 2020; Emberts et al., 2020; Forthman et al., 2020, 2022; Miller et al.; 2022), no studies have experimentally investigated the effect of altering hybridization temperatures on capture success of targeted loci in invertebrates.

Here, we evaluated the impact of four different protocols that varied hybridization temperatures on invertebrate target capture success in leaf-footed bugs and allies (Hemiptera: Coreoidea) and closely related taxa using samples from various sources (fresh versus older, dried specimens) and library qualities to reflect conditions typical of empirical studies. One protocol used a fixed, standard hybridization temperature (65°C) while the remaining three protocols employed touchdown approaches with different final temperatures. Specifically, we addressed the following questions: 1) do touchdown target capture approaches with lower hybridization temperatures result in a greater total of on-target reads, total assembled contigs, total targeted loci, and longer targeted assemblies compared to the commonly used standard hybridization temperature?; 2) does the touchdown target capture protocol with the lowest final temperature produce the most data for the variables listed in question 1?; and 3) does a lower hybridization temperature from touchdown capture protocols generate a greater proportion of off-target reads and/or paralogous sequences?

Our study utilized a subsampled version of an existing Hemiptera-derived UCE bait set (Faircloth, 2017; see Forthman et al., 2019), but we also introduced newly designed baits (with slightly greater tiling) derived from coreoid transcriptomes and evaluated the ability of these new baits to enrich samples *in vitro*. Given the different bait designs, we also examined whether the effects of hybridization temperature exhibited different patterns across bait design strategies; specifically, we investigated if 1) the proportion of on-target reads and coverage exhibit similar trends across hybridization temperature conditions regardless of bait design strategy, 2) the capture of loci with greater divergences from baits shows the greatest improvement at lower hybridization temperatures than loci with less divergence from their baits, 3) the number of putative paralogous loci recovered increases as hybridization temperature decreases regardless of bait design strategy, and 4) an increase in bait tiling improves read depth of captured loci.

### Novelty of the present study

Based on recent reviews (e.g., Andermann et al., 2020) and our own search of prior literature, only four studies—all in vertebrates—have explicitly tested the impacts of lower hybridization temperatures by comparing two temperature conditions (see Table 1). These studies have provided *mixed* results and recommendations on the *benefits* of implementing lower temperatures in capture experiments based on various metrics (Table 1). Thus, studies comparing different hybridization temperatures are needed in systems where bait-target divergences are much higher and prior capture studies have shown comparably lower target locus recovery.

**Table 1.** Experimental design of four studies that investigated the effects of target capture hybridization temperatures on various metrics used to assess capture performance.

| Characteristics of study | Li et al. (2013) | Paijmans et al. (2016) | Cruz-Dávalos et al. (2017) | Mohandesan et al. (2017) |
| --- | --- | --- | --- | --- |
| Taxon | Gnathostomata | Felidae | Equidae | *Camelus* (Camelidae) |
| Target data type | CDS | Mitogenome | SNP | Mitogenome |
| No. of targets | 1449 | 1 (entire mitogenome) | ~5000 (SNPs) | 1 (entire mitogenome) |
| Bait type | Biotinylated RNA | “Oligonucleotides” | Biotinylated RNA | Biotinylated RNA |
| Bait length (bp) | 120 | 60 | 60 | 80 |
| Tiling density | 2x | 30x | 3x | 4x |
| Sample quality | Fresh | Fresh & degraded | Degraded | Degraded |
| Target capture method | In-solution | Solid-state | In-solution | In-solution |
| Capture Protocol (CP) 1 | 65°C hybridization<br>(duration not reported) | 65°C hybridization;<br>captured targets used for<br>additional capture under<br>same conditions<br>(duration not reported) | 65°C hybridization<br>(24 hrs) | 65°C hybridization<br>(36 hrs) |
| Capture Protocol (CP) 2 | 65°C to 50°C hybridization<br>(5°C decrease every 11 hrs) | Same as CP1 but 50°C<br>hybridization | 55°C hybridization<br>(40 hrs) | 65°C to 50°C hybridization<br>(5°C decrease every 12 hrs) |
| Capture Protocol (CP) 3 | CP3 used captured targets &<br>same capture conditions from<br>CP 2 | 65°C to 50°C hybridization<br>(5°C decrease every 16.25<br>hrs); captured targets used<br>for additional capture under<br>same conditions |  |  |
| Metrics evaluated | Proportion of target loci<br>recovered<br>GC content<br>Bait-target divergence<br>Target length recovered<br>Chromosomal position | Proportion of on-target<br>reads<br>Target coverage<br>Bait-target divergence<br>(fresh tissues only) | Proportion of on-target<br>reads<br>Target coverage<br>Fold enrichment<br>Read length | Proportion of on-target<br>reads<br>Percentage of endogenous<br>DNA<br>Percentage of PCR<br>duplicate reads |

| Recommended CP based on metrics | CP 3 | CP 3 (fresh)<br>CP 1 (archival & ancient) | CP 2 (samples with low to medium endogenous DNA)<br>CP 1 & CP 2 not significantly different (samples with high endogenous DNA) | CP 1 & CP 2 performance not significantly different |
| --- | --- | --- | --- | --- |
| Relevant conclusions | Lower hybridization temperatures improve proportion of target loci recovered, particularly when bait-target divergence is high | Sample type & quality likely determines which hybridization temperature results in best capture based on proportion of on-target reads; higher temperatures may be better for degraded samples with greater amounts of contamination | Lower hybridization temperatures result in greater enrichment when samples have greater amounts of contamination but no impact when samples have higher amounts of endogenous DNA | Lower hybridization temperatures perform similarly with standard capture conditions for recovery of uniquely mapped reads or capture efficiency |

Our study addresses this gap by studying a group of invertebrates that are ∼160–230 my old (Li et al., 2017; Liu et al., 2019; Wang et al., 2019) and for which target capture studies have yielded low numbers of recovered loci (e.g., Forthman et al., 2019), which is likely due to high bait-target divergences. Our study also explores the effects of hybridization temperatures on other aspects of capture success not previously investigated in the studies listed in Table 1, such as the number of putative paralogs recovered between different hybridization temperature conditions.

Lastly, we explore intermediate temperatures to evaluate whether there is no further improvement to locus recovery or if costs outweigh the benefits at a particular temperature, which has also not been investigated by the studies in Table 1.

## Material and methods

### Sample material

Our target capture experiment was performed on 39 taxa (36 species of Coreoidea, 3 outgroup taxa), of which 30 were ethanol (EtOH), frozen, or silica bead (“fresh”) preserved samples (collected 2008–2017) and nine were degraded samples from pinned museum material of varying ages (1935–2017) (Table S1). These 39 taxa were selected for this experiment based on the availability of extra DNA libraries for additional target captures and to include a diversity of preservation methods and specimen ages (recent/fresh vs. historical/dried), as well as library qualities (best, moderate, and marginal quality based on initial sequencing outcomes relative to other samples). All taxa had previously been subjected to target capture protocols and sequenced prior to the start of this study (i.e., freshly preserved samples or dried preserved samples were subjected to the standard or TD-60 protocols shown in Fig. 1, respectively); capture data for 27 taxa have already been published following protocols described in Forthman et al. (2019, 2020) and Emberts et al. (2020) (see Table S1 and references therein).

**Figure 1.**
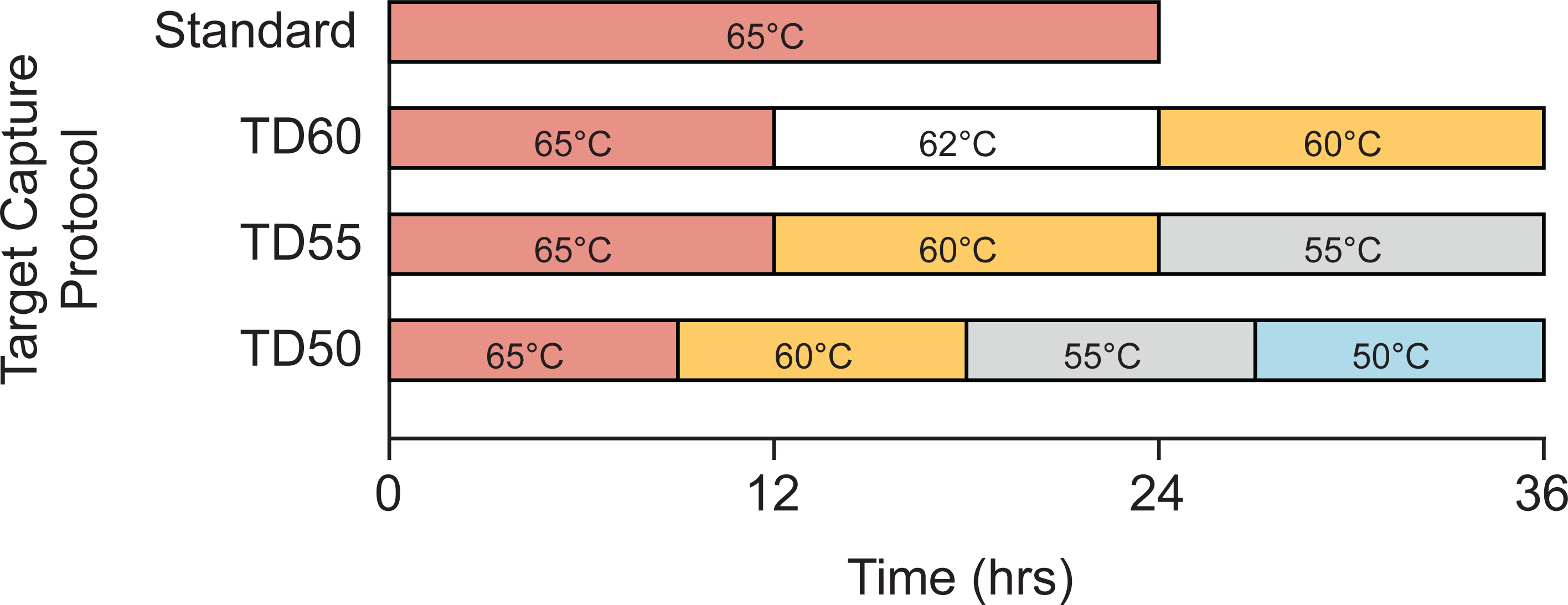
Experimental target capture set-up. Hybridization temperatures across time (in hours) are eported. Abbreviations: TD-60, touchdown hybridization approach starting at 65°C for 12 hrs followed by 62°C for 12 hrs and ending at 60°C for 12 hrs; TD-55, touchdown hybridization approach starting at 65°C for 12 hrs followed by 60°C for 12 hrs and ending at 55°C for 12 hrs; TD-50, touchdown hybridization approach starting at 65°C for 9 hrs followed by 60°C for 9 hrs, 55°C for 9 hrs, and ending at 50°C for 9 hrs.

### Target capture baits

For a list of terms and their definitions used in this study, see Table 2. A summary of bait properties from our different bait design strategies are given in Fig. 2.

**Figure 2.**
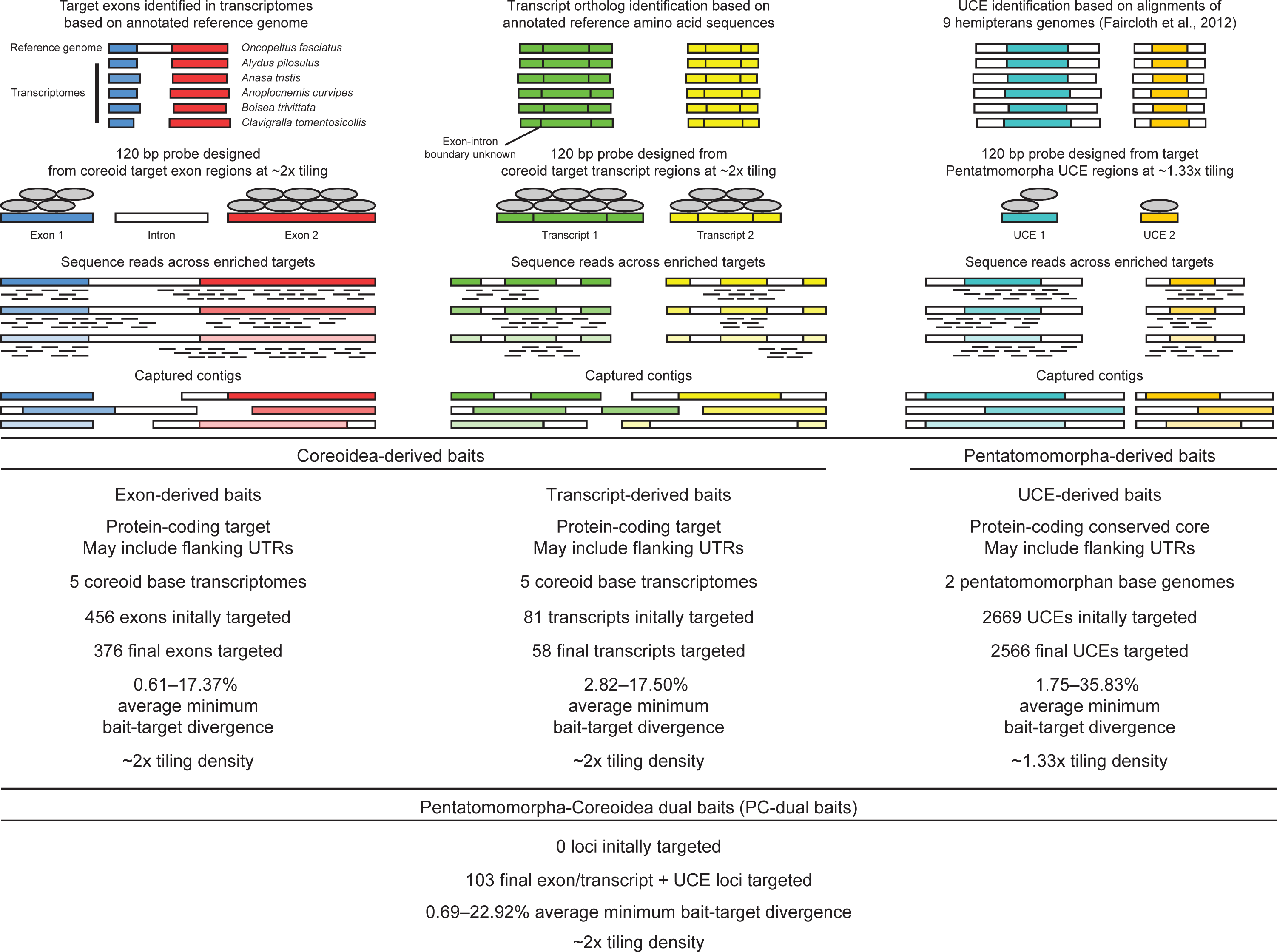
Design and properties of bait used in this study. Abbreviations: bp, base pair; UCE, ultraconserved element; UTR, untranslated region.

**Table 2.** Terminology related to bait design and targeted loci.

| Term | Definition |
| --- | --- |
| Ultraconserved element (UCE) baits | 120 bp baits designed to target highly conserved genomic regions at ~1.33x tiling. A Hemiptera-wide UCE bait set was designed by Faircloth (2017), which targets the conserved core of protein-coding loci (Kieran et al., 2019); however, contigs captured by UCE baits may also include introns and/or untranslated regions (UTRs). |
| Pentatomomorpha-derived baits | The Faircloth (2017) Hemiptera-wide UCE bait set was subsampled to include only those baits designed from the genomes of two species within the infraorder Pentatomomorpha, which are the closest relatives to the ingroup of this study. |
| Exon-derived baits | 120 bp baits designed at ~2x tiling from individual exon sequences (i.e., baits do not span across introns) exhibiting $\geq 60\%$ conservation across transcriptomes from five species of Coreoidea. While the baits do not target UTRs, given that they are derived from transcriptome sequences, contigs captured by these baits may include introns and/or UTRs. |
| Transcript-derived baits | 120 bp baits designed at ~2x tiling from sequences that may include one or more exons and that exhibit $\geq 60\%$ conservation across transcriptomes from five species of Coreoidea. While the baits do not target UTRs, given that they are derived from transcriptome sequences, baits may span across one or more introns and, thus, contigs captured by these baits may include introns and/or UTRs. |
| Coreoidea-derived baits | Baits designed primarily from transcriptomes of five species of Coreoidea (i.e., exon- and transcript-derived baits combined). |
| Pentatomomorpha-Coreoidea (PC) dual baits | Loci targeted by Pentatomomorpha-derived UCE baits as well as by exon-derived or transcript-derived baits (i.e., Coreoidea-derived baits). |

We used our previously published custom myBaits kit (Forthman et al., 2019), which subsampled a Hemiptera-wide derived UCE bait set (at ∼1.33x tiling) designed by Faircloth (2017) to only include two pentatomomorphan taxa that are more closely related to but not included in our ingroup taxa (herein, referred to as “Pentatomomorpha-derived baits”; Table 2; Fig. 2). This kit also included an independently designed set of baits that were derived from coreoid transcriptomes, but these have not yet been introduced in the literature prior to this study. Thus, we introduce our coreoid bait design procedures, which included two design strategies: baits were designed from individual exons while others were designed across entire transcripts (collectively referred to as “Coreoidea-derived baits”). We provide specific details on our bait design procedures for the Coreoidea-derived baits in the Supplementary Materials, but we provide a brief overview of the different sets of new baits below.

First, we designed baits from individual coreoid exon sequences (herein, “exon-derived baits”, which is a subset of the Coreoidea-derived baits; Table 2; Fig. 2), using localized reciprocal blastn searches with an annotated draft genome of *Oncopeltus fasciatus* (Dallas, 1852) (Lygaeidae) from the Baylor College of Medicine – Human Genome Sequencing Center (https://www.hgsc.bcm.edu/arthropods/milkweed-bug-genome-project) and five unannotated coreoid transcriptomes: *Alydus pilosulus* Herrich-SchDffer, 1847 (Johnson et al., 2018; NCBI BioProject PRJNA272214), *Anasa tristis* (De Geer, 1773) (Johnson et al., 2018; NCBI BioProject PRJNA272215), *Anoplocnemis curvipes* (Fabricius, 1781) (Agunbiade et al., 2013; NCBI BioProject PRJNA192258), *Boisea trivittata* (Say, 1825) (Johnson et al., 2018; NCBI BioProject PRJNA272221), and *Clavigralla tomentosicollis* Stål, 1855 (Agunbiade et al., 2013; NCBI BioProject PRJNA192261). Sequences for 456 individual exons were selected for part of the Coreoidea-derived bait set, which were ≥200 bp in length, had a GC-content between 30%–70%, and did not include low complexity sequences and simple repeats.

Using the same genomic resources above, we then designed a second set of baits across transcript sequences following the approaches of Portik et al. (2016) (i.e., RNAseq-derived sequences that may include more than one exon, with baits potentially spanning across one or more introns; herein, “transcript-derived baits”; Table 2; Fig. 2). Based on annotations, 141 transcripts (out of 172 initially selected) did not correspond to any of the 456 exons selected for bait design. Of these, we selected 81 transcripts for part of the Coreoidea-derived baits that were found across most of our coreoid transcriptomes and that ranged between 256 bp and 1000 bp to reduce the number of baits potentially targeting multiple exons interspersed by long introns.

The final Coreoidea-derived bait sequences (i.e., exon- and transcript-derived baits) were submitted to Arbor Biosciences (Ann Arbor, MI) to produce 120 bp baits with ∼2x tiling density. We also preliminarily compared our Coreoidea-derived baits against the Pentatomomorpha-derived baits using blastn (e-value threshold set to 1e-20) to determine whether any were potentially associated with a locus already targeted by the latter set of baits. Those Coreoidea-derived bait sequences matching to Pentatomomorpha-derived baits were not removed from the final selection of baits, as their inclusion could allow for some targeted loci to be captured by more baits from more closely related species (herein, “Pentatomomorpha-Coreoidea [PC] dual baits”; Table 2; Fig. 2).

### DNA extraction, library preparation, and target capture

See references in Table S1 for details on library preparation and target capture of previously published samples. For new samples, we extracted genomic DNA from any part of the body or the entire body from EtOH-preserved, silica-bead preserved, frozen, or dried specimens to sample similar amounts of tissue across taxa, where possible (Table S1), and constructed libraries following procedures we have used elsewhere (e.g. Forthman et al., 2019, 2020, 2022) and outlined in the Supplementary Methods.

To evaluate the effects of different hybridization conditions on target capture, we compared four experimental protocols using 1/2 or 1/4 volume baits for fresh and dried samples, respectively, which differed only in hybridization temperatures used over a 24- (common in standard target capture protocols) or 36-hour period (see Fig. 1). Due to the limited availability of extra DNA libraries, previously sequenced samples could only be assigned to one of three different target capture protocols (see Table S2). In assigning samples to protocols, we attempted to distribute sample preservation methods, sample age, and library quality (based on our initial sequencing efforts).

All post-capture protocols followed Forthman et al. (2019), with the exception that captures were washed at temperatures corresponding to the final hybridization temperature used in each capture protocol (i.e., 65°C for standard, 60°C for TD-60, etc.). All enriched library pools were combined in equimolar amounts and sequenced on an Illumina HiSeq3000 lane (2x100) at the University of Florida’s Interdisciplinary Center for Biotechnology Research.

### Sequence data processing and analysis

Unless otherwise stated, all data processing steps and analyses mentioned below used default settings. Sequence reads were demultiplexed by the sequencing facility. Adapters were trimmed with illumiprocessor (Faircloth, 2013; Bolger et al., 2014). Duplicate reads were filtered with PRINSEQ-lite. Reads were then error corrected using QuorUM and subsequently assembled *de novo* using SPAdes v3.13.0 with the single-cell and auto coverage cutoff options invoked (Bankevich et al., 2012; Nurk et al., 2013; Prjibelski et al., 2020). PHYLUCE v1.5.0 (Faircloth, 2016) was then used to extract targeted loci from assembled contigs.

Because our preliminary comparison of loci targeted by Coreoidea-derived baits against the Pentatomomorpha-derived baits prior to bait design indicated that some loci were targeted by both sets of baits, a more thorough confirmation was performed after *in vitro* target capture. Using captured loci targeted by our Coreoidea-derived baits, we performed a tblastx (e-value threshold = 1e-10) search against those loci captured by the Pentatomomorpha-derived baits and extracted matches with ALiBaSeq (Knyshov et al., 2021). Of the loci captured by the Pentatomomorpha-derived baits, we found that 103 of these were targeted by both Pentatomomorpha- and Coreoidea-derived baits. It is worth noting that during this process, we also determined that some targeted transcript loci were also targeted by multiple, adjacent UCE loci by Pentatomomorpha-derived baits; in such cases, we treated these loci as a single locus. Thus, we had 376 loci targeted by exon-derived baits, 58 targeted by transcript-derived baits, 2566 targeted by Pentatomomorpha-derived baits, and 103 targeted by both Pentatomomorpha- and Coreoidea-derived baits (i.e., PC dual baits), resulting in a total of 3103 targeted loci.

We calculated the number and lengths of assembled contigs and captured loci using PHYLUCE. Because our Pentatomomorpha- and Coreoidea-derived baits were designed from different sets of genomes and transcriptomes of varying divergences, we also calculated the average minimum distances between our baits and captured loci for exon-derived, transcript-derived, Pentatomomorpha-derived, and PC dual baits. We then calculated read depth using our filtered reads and the total number of on-target filtered reads using BBMap v38.44 (Bushnell, 2014). We determined the number of captured loci with putative paralogs by invoking the keep-duplicates option in the PHYLUCE phyluce_assembly_match_contigs_to_probes.py script.

Off-targets reads may contain sequences from loci traditionally used in phylogenetic studies (herein referred to as “legacy loci”) (e.g., Amaral et al., 2015; Wang et al., 2017; Simon et al., 2019; Miller et al., 2022). As such, we also extracted 15 mitochondrial and two nuclear legacy loci from off-target contigs in our target capture data for comparison between capture protocols (see Appendix 1 for further details on background, methods, and results pertaining to legacy loci).

As one part of the experiment, we also wanted to quantify the effect of different tiling strategies (∼1.33x vs. ∼2x tiling density) on locus recovery. As most loci with baits tiled differently also exhibited major differences in bait-target divergences (see Results), we were not able to directly measure the effect of tiling strategy for most loci. However, some loci (i.e., 40 out of 2566) captured with Pentatomomorpha-derived baits had low average minimum bait-target divergences similar to loci captured with Coreoidea-derived bait (see Results). Thus, we had a limited opportunity to explore the effect of tiling strategy on read depth while controlling for bait-target divergences. For this, we selected loci captured with Coreoidea- or Pentatomomorpha-derived baits that had divergences ranging from 0.05–0.10 and calculated read depth.

Sequencing depth between different sequence lanes/runs may affect how many loci are recovered or the proportion of on-target reads. Furthermore, the effective sample size across different sequencing efforts can vary as some samples may fail or have poor sequencing outcomes, which can have an impact on sequencing depth across samples. Our samples were subjected to three different sequencing efforts, with each producing different effective sample sizes: 1) 99 samples combined for previously sequenced samples subjected to the standard capture conditions, 2) 96 samples combined for previously sequenced samples subjected to the TD-60 capture conditions and 3) 88 samples combined for sequenced samples subjected to the different target capture conditions conducted in this study. To investigate the influence of different sequencing depths on our metrics of capture success, we equalized sequencing depth by subsampling 2,000,000 raw reads generated under the different capture protocols for 24 taxa that were preserved fresh using Seqtk v1.3 (https://https://github.com/lh3/seqtk) (random seed [-s option] = 100). Eleven taxa were not included in this analysis because at least one of the target capture protocols for each of these taxa were associated with fewer than 3,000,000 raw reads total; this drastically lowered the sample size of available samples that were preserved dried (four taxa remaining), and thus, we excluded dried samples. The subsampled reads were then processed and evaluated as described above to determine if patterns observed in the subsampled dataset differed from what was observed in the original data.

### Statistical analyses

Statistical analyses using linear mixed models (LMMs), generalized linear models (GLMs), and generalized linear mixed models (GLMMs) were performed using *lme4* v1.1.30 (Bates et al., 2015) in R v4.1.2 (R Core Team, 2022). We excluded dried samples from statistical analyses due to very low sample size for some target capture protocols, with the exception that they were included with fresh samples when analyzing bait-target divergences since this was independent of target capture protocol. We refrained from treating library quality as a factor in analyses since this was used as an *a posteriori* criterion for selecting samples to be included in a second capture protocol. We also did not include the different sequencing efforts across samples as a random effect since it has been suggested that random effect terms should have at least five levels for inclusion (Harrison, 2015; but see Gomes, 2022). However, to account for potential effects of different sequencing efforts, we subsampled our data as described in the previous section to normalize sequencing depth across sequencing efforts. For LMM and GLMM analyses, we treated target capture protocol as a fixed effect. Because the same sample was subjected to two capture protocols, we included replication as a random effect. The GLM analyses treated bait design strategy as a fixed effect. For specific details regarding the response variables and specific structure of all models analyzed, see Supplementary Methods. For all analyses, we performed simultaneous tests for general linear hypotheses using Tukey contrasts for multiple comparisons of means using the *multcomp* v1.4.20 R package (Hothorn et al., 2008).

## Results

### Raw read and assembled contig yield across target capture conditions

Overall, lower hybridization temperatures generated significantly more raw sequence reads for samples preserved fresh (Table S3; Fig. 3A), with dried samples exhibiting similar trends (Fig. S1A). Of the 39 samples in our dataset, 26 had more raw reads sequenced when these samples were subjected to lower hybridization temperatures in pairwise comparisons (median increase = 272%) (Table S3). The TD-50 capture protocol produced the most raw reads for freshly preserved samples (Fig. 3A), while the TD-50 and TD-55 produced the most for dried samples (Fig. S1A). Three degraded (i.e., preserved dried) samples failed or nearly failed to produce any raw reads under the standard protocol, but they had >33,000 reads sequenced at a modestly lower hybridization temperature (TD-60) (Table S3).

**Figure 3.**
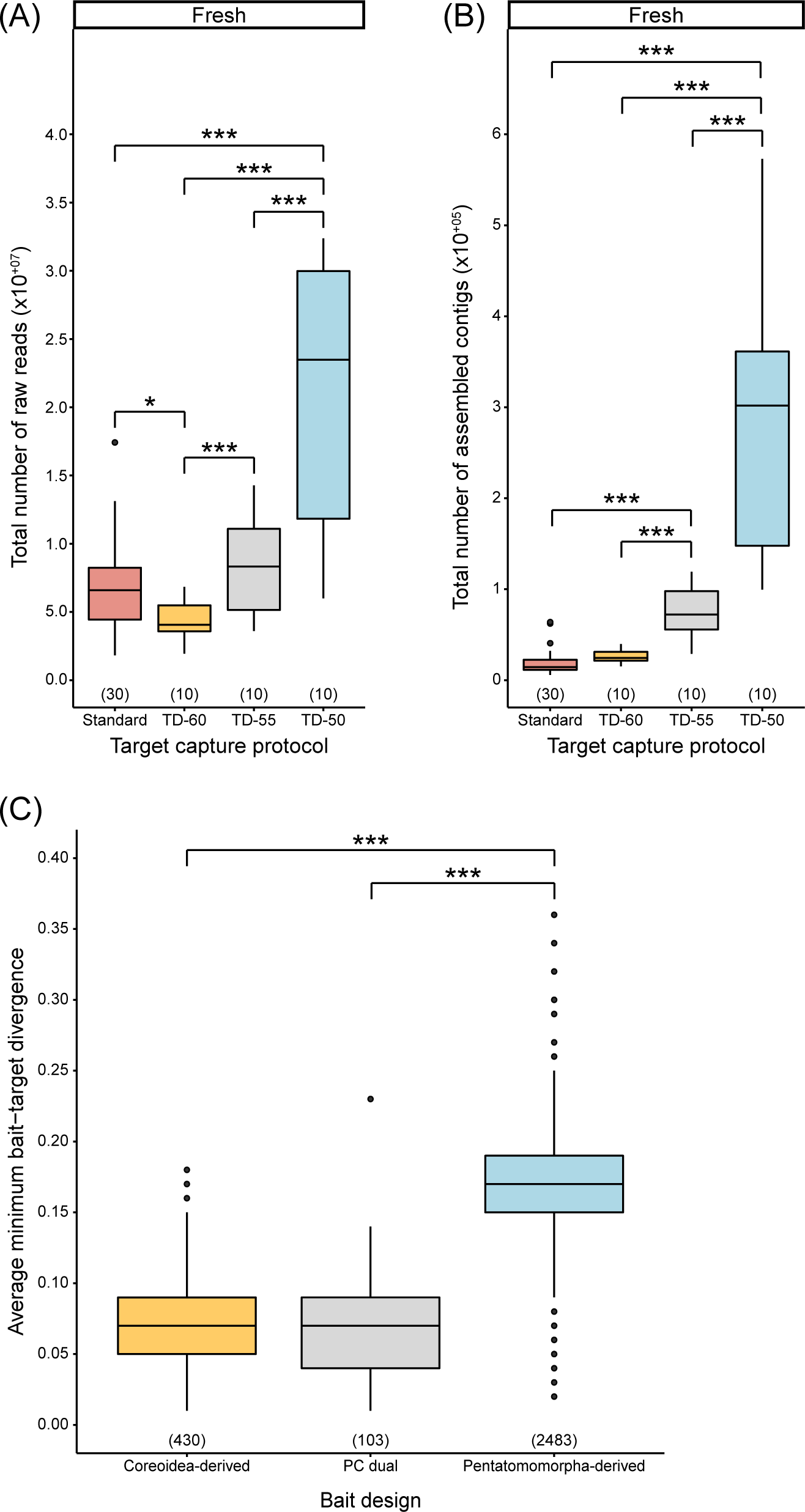
(A) Effects of target capture protocols on the total number of raw reads and (B) assembled contigs for samples preserved fresh. (C) Average minimum bait-target divergences by bait design strategy. Numbers in parentheses above x-axis denote sample size. Single and triple asterisks denote statistically significant pairwise comparisons, with p < 0.050 and p < 0.001, respectively. Abbreviations: PC dual baits, loci targeted by both Pentatomomorpha- and Coreoidea-derived baits.

In general, touchdown protocols also produced a greater total of assembled contigs and nucleotides (Table S3; Figs. 3B,S1B). Contigs had similar ranges of median lengths between capture protocols, but, for freshly preserved samples, all the touchdown protocols had significantly longer median contig lengths than the standard protocol while there were no statistical differences between touchdown protocols (Fig. S1C). For dried samples, median contig length exhibited increasing trends as hybridization temperature was lowered (Fig. S1D). Data generated under protocols with lower hybridization temperatures yielded some of the longest contigs recovered (Table S3). Thus, while lower hybridization temperatures varied in their success when looking at a given taxon, the lowest hybridization temperature resulted in the most raw reads and contigs across samples, on average.

### Bait-target distances, reads on-target, and read depth

We found that loci targeted by either Coreoidea exon- and transcript-derived bait sequences followed similar trends in our experiment, as well as shared similar average minimum bait-target divergences (Fig. S1D). Due to this, as well as shared genomic resources during bait design and the relatively few number of loci, we combined results from the exon- and transcript-targeted loci together, which we refer to as Coreoidea-derived baits (for separate exon- and transcript-targeted locus results, see Tables S4–S6 and Figs. S2–S4). Similar average minimum bait-target divergences were observed for loci captured by Coreoidea-derived baits and PC dual baits, and these were significantly lower and exhibited less variation compared to those captured by Pentatomomorpha- derived baits (Fig. 3C).

There was a general increase in the number of filtered on-target reads in pairwise comparisons as hybridization temperature was lowered (Table S4). Despite this, the TD-50 capture protocol had a significantly lower proportion of on-target reads across all targeted loci for freshly preserved samples (Fig. 4A). When partitioning by bait design strategy, this pattern remained for freshly preserved samples, but with the TD-55 protocol also resulting in a lower proportion of on-target reads compared to the TD-60 and standard protocols (Fig. 4B–D). Dried samples did not show any noticeable trends as hybridization temperature was lowered, with the exception that all touchdown capture protocols had a higher proportion of on-target reads compared to the standard protocol (Fig. S5).

**Figure 4.**
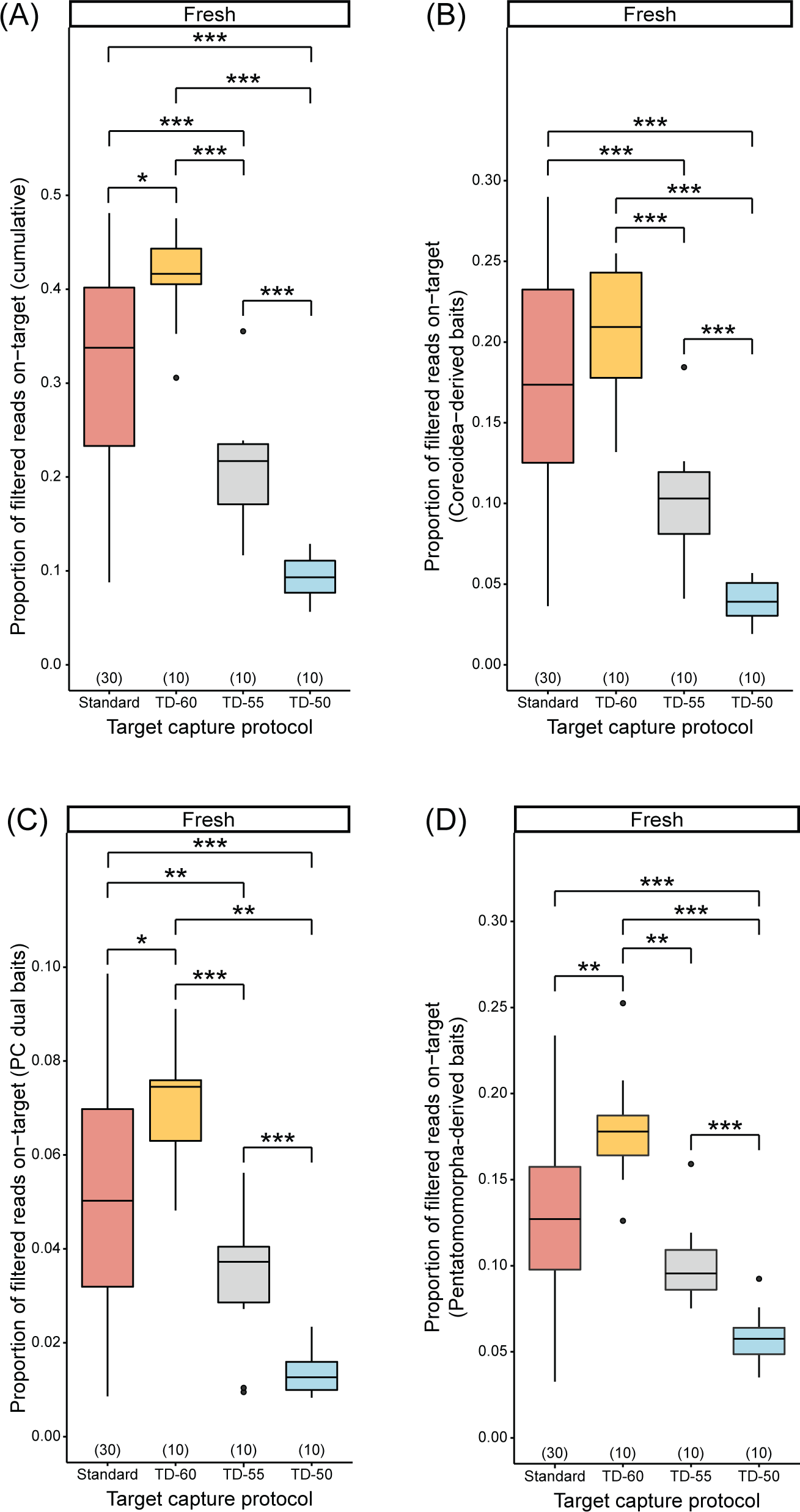
Effects of target capture protocols on the proportion of filtered reads on-target (A) across all argeted loci, (B) loci captured by Coreoidea-derived baits, (C) loci captured by PC dual baits, and (D) oci captured by Pentatomomorpha-derived baits for samples preserved fresh. Numbers in parentheses above x-axis denote sample size. Single, double, and triple asterisks denote statistically significant pairwise comparisons, with p < 0.050, p < 0.010, and p < 0.001, respectively. Abbreviations: see Figs. 1 and 4.

In pairwise comparisons, there was frequently a decrease in read depth at lower temperatures regardless of bait design strategy, with overall read depth significantly different between the TD-50 and TD-55 protocols and the standard protocol for freshly preserved samples only (Tables S4; Fig. 5A). However, this pattern was not consistently observed when partitioning by bait design strategy (Fig. 5B–D): 1) the TD-55 capture protocol did not have a significantly lower read depth for loci targeted by Coreoidea-derived baits compared to the standard protocol; 2) read depth of loci targeted by PC dual baits was not significantly different across capture protocols; and 3) Pentatomomorpha-derived baits from the TD-60 and TD-55 protocols were significantly lower than the standard, but were not significantly different from each other nor the TD-50 protocol. For dried samples, read depth did not appear to exhibit any trends across capture protocols, with the exception that lower temperatures exhibited greater read depth for loci targeted by Coreoidea-derived and PC dual baits (Table S4; Fig. S6).

**Figure 5.**
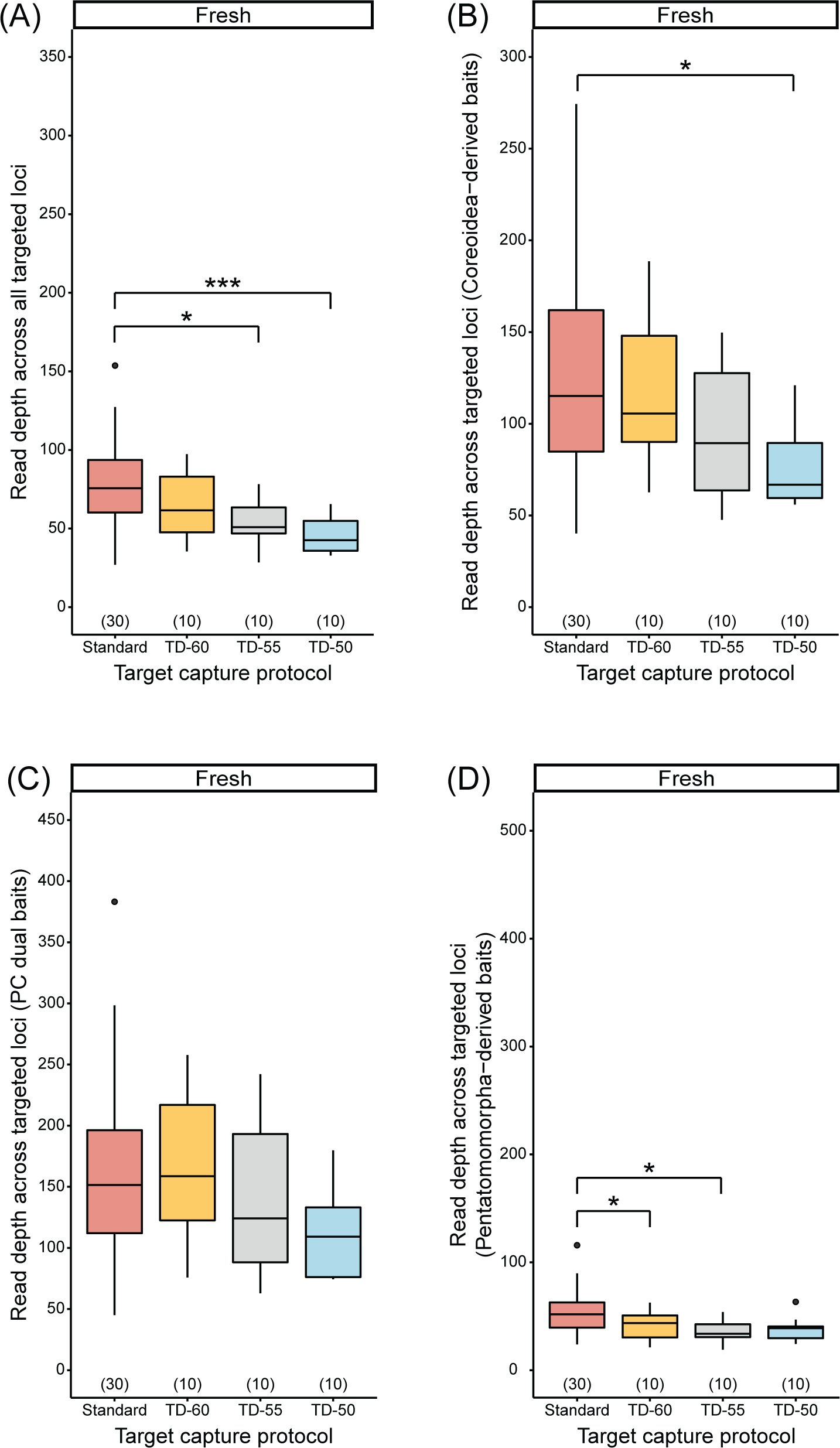
Effects of target capture protocols on read depth (A) across all targeted loci, (B) loci captured by Coreoidea-derived baits, (C) loci captured by PC dual baits, and (D) loci captured by Pentatomomorpha-derived baits for samples preserved fresh. Numbers in parentheses above x-axis denote sample size. Single and triple asterisks denote statistically significant pairwise comparisons, with p < 0.050 and p < 0.001, espectively. Abbreviations: see Figs. 1 and 4.

### Targeted locus recovery, locus lengths, and putative paralogs

Lower hybridization temperatures produced significantly more loci targeted by Coreoidea- and Pentatomomorpha-derived baits compared to the standard temperature for freshly preserved samples, with the TD-50 protocol resulting in the highest proportion of loci recovered (Tables S5– S8; Fig. 6A–C). The median lengths of targeted loci were significantly longer when the TD-50 protocol was used compared to higher hybridization temperatures for freshly preserved samples, regardless of bait design strategy (Tables S5–S8; Fig. 6D–F). Dried specimens did not exhibit any trends with respect to target capture protocol and the proportion of targeted loci recovered or the median lengths of targeted loci (Tables S5–S8; Figs. S7,S8).

**Figure 6.**
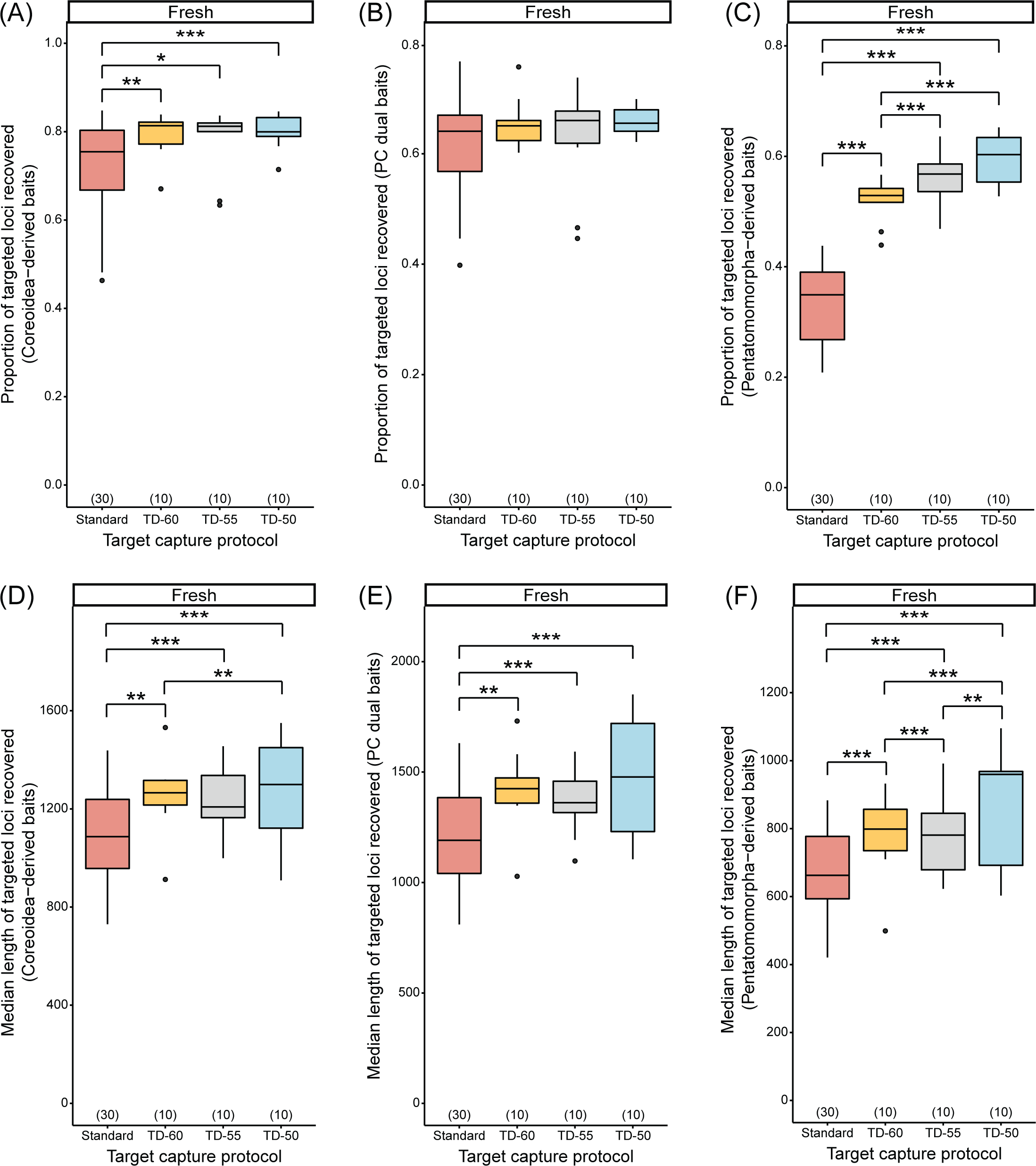
Effects of target capture protocols on the proportion of recovered loci targeted by (A) Coreoidea-derived baits, (C) PC dual baits, and (D) Pentatomomorpha-derived baits for samples preserved fresh. Effects of target capture protocols on the median length of recovered loci targeted by (D) Coreoidea-derived baits, (E) PC dual baits, and (F) Pentatomomorpha-derived baits for samples preserved fresh. Numbers in parentheses above x-axis denote sample size. Single, double, and triple asterisks denote statistically significant pairwise comparisons, with p < 0.050, p < 0.010, and p < 0.001, respectively. Abbreviations: see Figs. 1 and 4.

We did not observe any patterns with respect to hybridization temperatures and the number of putative paralogs detected for Coreoidea-derived and PC dual baited strategies for samples preserved fresh (Tables S5–S7; Figs. 7A,B). In contrast, we observed increases in putative paralogs associated with loci targeted by the Pentatomomorpha-derived baits as hybridization temperatures decreased (Table S8; Fig. 7C). We did not observe any noticeable trends with respect to hybridization temperatures and putative paralogs detected by any bait strategy for dried samples (Tables S5–S8; Fig. S9).

**Figure 7.**
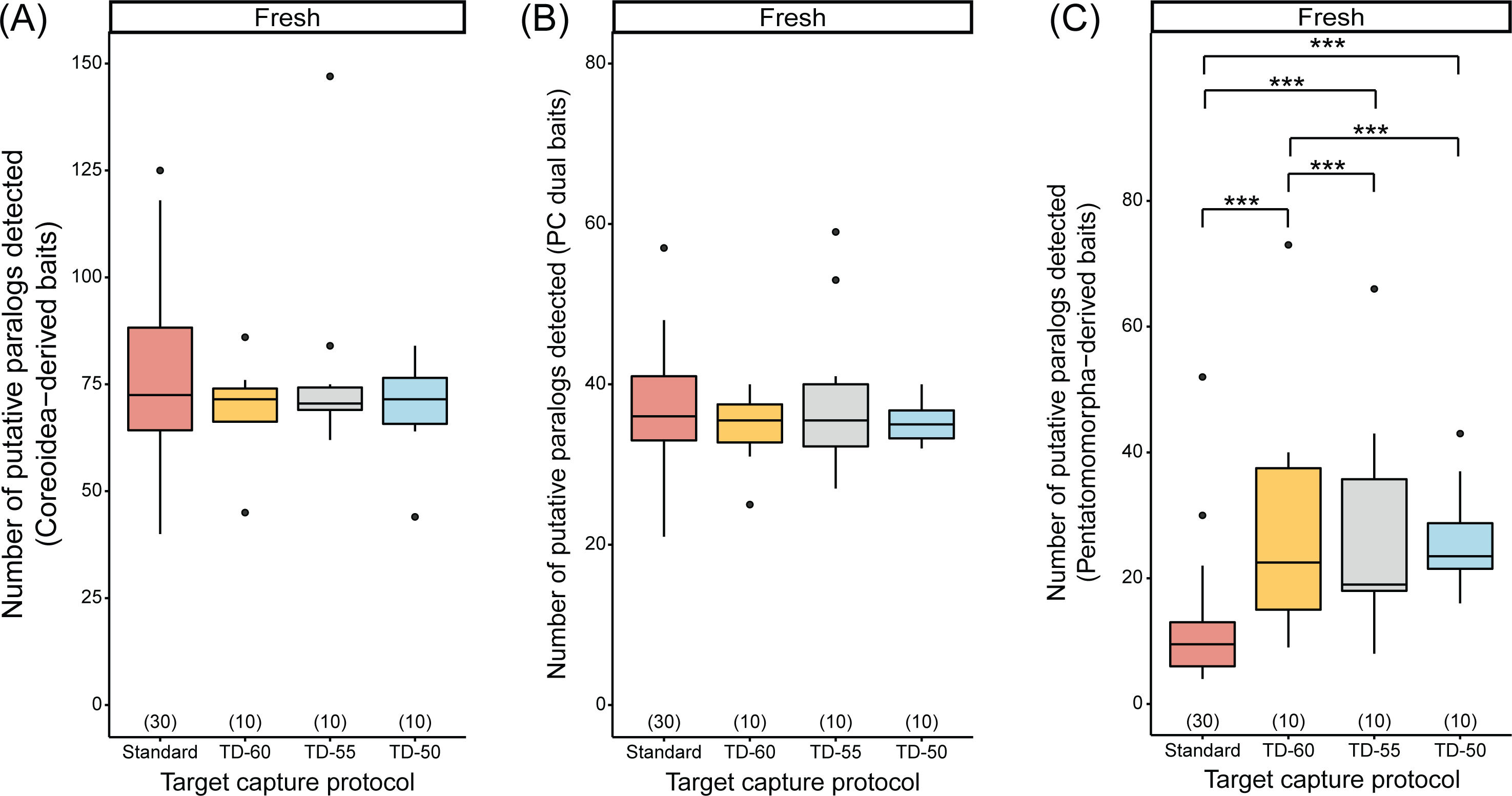
Effects of target capture protocols on the number of putative paralogs of loci captured by (A) Coreoidea-derived baits, (C) PC dual baits, and (D) Pentatomomorpha-derived baits for samples preserved resh. Numbers in parentheses above x-axis denote sample size. Triple asterisks denote statistically significant pairwise comparisons, with p < 0.001. Abbreviations: see Figs. 1 and 4.

### Additional measures and parameters investigated

#### Tiling strategies

We quantified the effect of different tiling strategies on one aspect of locus recovery, read depth. Most loci with baits tiled differently exhibited major differences in bait-target divergences (Fig. 3C), but 40 loci captured with Pentatomomorpha-derived baits had low average minimum bait-target divergences similar to loci captured with Coreoidea-derived bait. We selected loci captured with Coreoidea-or Pentatomomorpha-derived baits that had divergences ranging from 0.05–0.10 to control for bait-target divergences and explore the effect of tiling strategy on read depth. We found that loci captured by Coreoidea-derived baits (∼2x tiling) had significantly greater average read depth than those captured by Pentatomomorpha-derived bait (1.33x tiling; Fig. S10).

#### Effects of sequencing depth on freshly preserved samples

To investigate the influence of sequencing depth on our metrics of capture success, we evaluated whether the capture success metrics for 24 freshly preserved taxa were still congruent when reads were subsampled evenly (Tables S9–S14; Figs. S11–S16). In general, patterns across hybridization temperatures remained congruent for most metrics compared to when reads were not subsampled, including: total number of assembled contigs, proportion of filtered reads on-target, proportion of targeted loci recovered, and number of putative paralogs found (Tables S9–S14; Figs. S11A, S11C, S12, S14, S16).

In contrast the median length of assembled contigs was not significantly different across hybridization temperatures but exhibited similar ranges of variation and general trends compared to those when reads were not subsampled (Table S9; Fig. S11B). We also found read depth across targeted loci to decrease significantly as hybridization temperatures were reduced, regardless of bait design strategy (compared to little or no significant reductions when reads were not subsampled) (Tables S9, S10; Figs. S11D, S13). Lastly, the median lengths of targeted loci recovered were significantly shorter at lower hybridization temperatures (Tables S11–S14; Fig. S15), in contrast to the significantly longer loci at these same temperatures when reads were not subsampled.

## Discussion

Invertebrate target capture studies employing fixed, high hybridization temperatures often recover relatively low proportions of targeted loci. For many taxonomic groups, like the leaf-footed bugs studied here, genomic resources are lacking or too limited to adequately optimize target capture baits to improve locus recovery. Furthermore, optimizing or re-designing target capture baits to improve locus recovery may result in more bait sets that are too narrowly targeted from a taxonomic perspective (e.g., a family) when there can be advantages of making bait sets more broadly applicable (e.g., a superfamily, infraorder, or higher ranks). Thus, modifying target capture conditions may be the best approach to improve the success of capture experiments across a broader taxon sampling.

Our evaluation of four different in-solution capture protocols found hybridization temperatures lower than the standard 65°C led to more assembled contigs and improved recovery of targeted loci, which held true even after normalizing sequencing depth. While this improvement was observed for loci targeted by the less divergent Coreoidea-derived baits, this was even more apparent for loci targeted by the more divergent Pentatomomorpha-derived baits. Such an improvement occurred despite touchdown capture ending at 50°C generally having a negative impact on the proportion of on-target reads and read depth, as well as increased numbers of putative paralogous loci when using the Pentatomomorpha-derived baits. We also found that lower hybridization temperatures also led to a greater recovery of other common loci of historical use from off-target reads (particularly mitochondrial regions), which is an alternative approach to designing custom baits for such loci or relying on low coverage genome sequencing (see Appendix S1 for more information on this topic and specific results). Given our results, we find that optimizing *in vitro* target capture conditions to accommodate low hybridization temperatures can provide a cost-effective solution to improve the successful recovery of targeted loci in invertebrates (albeit with some costs), while also providing opportunities to gain additional data.

Our touchdown capture protocols were performed with longer durations than the standard protocol (36 hrs vs. 24 hrs, respectively), and one of these touchdown protocols had four incremental temperature decreases every 9 hrs instead of three incremental decreases every 12 hrs (Fig. 1). Thus, the total capture duration and duration of the incremental temperature decreases might be considered a confounding factor in our experiment. However, two of our target capture protocols (TD-60 and TD-55) used the same timing (36-hour total duration, with temperature decreases every 12 hrs), and, as with all other capture protocols we used, the lower hybridization temperature (i.e., TD-55) improved target capture success for some metrics (such as number of raw reads, contigs, and number of recovered loci targeted by Pentatomomorpha-derived baits) while exhibiting negative effects on other metrics (e.g., less reads on-target or more putative paralogs of recovered loci targeted by Pentatomomorpha-derived baits). Thus, the duration of target capture or the duration of bait-target hybridization at specific temperatures does not appear to be a major confounding factor in our study.

### Considerations for using transcript-derived baits

We employed a bait design procedure in which coreoid transcriptomes were used to identify individual exons that baits would target, as well as entire transcripts that could contain multiple exons with baits potentially dissected by introns (i.e., a single bait that is derived from two exons). Based on our metrics—irrespective of target capture conditions (particularly the number of targeted loci recovered)—, we found that our exon-derived baits were highly successful (78% recovery, on average) compared to the transcript-derived baits (33% recovery, on average). This finding indicates that most of the 58 loci targeted by transcript-derived baits could not be captured. The poor *in vitro* performance of our transcript-derived baits may be due to transcript sequences comprised of many short exons leading to many baits dissected by introns. This could result in multiple assembled contigs if the entire intron(s) was not sequenced, which would result in the associated transcript baits matching multiple “different” contigs in PHYLUCE and exclusion of these sequences due to putative paralogy. Thus, our transcript-derived baits do not appear very useful in Coreoidea target capture studies, and our transcript bait design strategy may not be suitable for other groups in general unless exon-intron boundaries are known or different bioinformatic strategies are used.

### Considerations for using low hybridization temperatures

Low hybridization temperatures were associated with improved recovery of loci targeted by the Coreoidea- and Pentatomomorpha-derived UCE baits, with the latter exhibiting markedly improved success. These results, as well as the lack of statistical differences in target locus recovery across capture conditions for PC dual baited loci, suggest that other factors, such as bait-target divergences and bait tiling strategy, also impacted our capture experiments.

Previous studies suggest that the efficacy of target capture decreases when bait-target divergences exceed 5–10% (Vallender, 2011; Bi et al., 2012; Paijmans et al., 2016), although successful capture has been reported when divergences are much higher (e.g., 40% in Li et al. [2013]). The Pentatomomorpha-derived baits used here were derived from two taxa in the same infraorder as the leaf-footed bugs (Faircloth, 2017), but these taxa diverged from coreoids ∼160– 230 mya (Li et al., 2017; Liu et al., 2019; Wang et al., 2019). The Pentatomomorpha-derived baits generally exhibited relatively high levels of divergence from the targeted loci in coreoids, sometimes as high as that seen in Li et al. (2013). By reducing target capture stringency via lower hybridization temperatures, a greater mismatch tolerance between divergent baits and targets can lead to improved locus recovery, which our study supports. Given that our Coreoidea-derived baits or PC dual baits were designed from coreoid transcriptomes, we were able to target loci with much lower divergences from their respective baits (<10%).

Bait tiling strategy also appears to have played a role, although we could not thoroughly explore its effects on many of our metrics. Despite some loci targeted by the Pentatomomorpha-derived baits exhibiting similar levels of divergences as those targeted by Coreoidea-derived baits, we found that a modest increase in bait tiling density (i.e., the latter set of baits) was associated with greater read depth of the targeted loci. Lower bait-target divergences and/or greater tiling of baits across targeted loci likely explain the relatively high proportion of loci recovered by these Coreoidea-derived baits compared to those recovered with Pentatomomorpha-derived baits.

If locus recovery is the main goal of invertebrate capture studies (especially if bait-target divergences exhibit a broad range of variation as in our study), using less stringent hybridization temperatures can be particularly beneficial when baits and targets are expected to exhibit high divergences and/or when baits are tiled at a low density. The lowest hybridization temperature tested (TD-50) resulted in the best target locus recovery, especially for the more divergent target loci. This result is likely due in part to the higher amount of data (i.e., reads and contigs) obtained for this treatment. However, when we subsampled raw reads across all capture conditions to normalize sequencing depth, we found that the TD-60 protocol provided the best balance with respect to the proportion of on-target reads and lower hybridization stringency allowing for maximal target locus recovery. Thus, to maximize target data for many samples with relatively lower sequencing depth, the TD-60 protocol may be preferable, but if sequencing depth is not a critically limiting factor, the TD-50 protocol may be a more desirable option.

It is expected that lowering hybridization temperature to reduce target capture stringency should result in a greater proportion of off-target than on-target sequences in capture data due to greater mismatch tolerance. Previous studies have observed this pattern (Paijmans et al., 2016 [ancient and archival specimens]; Cruz-Dávalos et al., 2017), which we also support here. More off-target data may be considered undesirable for many studies, and for phylogenomic studies, this could include potential sequences deemed paralogous to target data. When putative paralogs are detected in some sequence processing pipelines like PHYLUCE, both the target and putative paralog are excluded from further analysis, so even targeted data may be excluded from final analyses. We found more paralogs recovered for loci targeted by Pentatomomorpha-derived baits, but not other targeted loci, which may be related to the degree of bait-target divergences or tiling strategy. Thus, for capture studies using baits expected to be quite divergent from ingroup taxa, lower hybridization temperatures can result in more targets recovered, but possibly at the cost of recovering more paralogous loci and a loss of some corresponding true targets. However, further evaluation of putative paralogs and potential targets can often prevent loss of data, especially since bioinformatic pipelines like PHYLUCE can report information on multiple contigs matching to a bait. This then allows analyzing sequences more carefully with bioinformatic tools like BLAST, inspecting alignments containing reference sequences, or refining ortholog binning procedures in bioinformatic pipelines to also use untargeted paralogs in analyses. Accurate paralog detection steps are very important regardless of hybridization temperatures (and how many paralogs are expected to be recovered) as any undetected paralogs have the potential to mislead phylogenetic inference.

Tissue quality appears to have some influence on the trends we observed with respect to hybridization temperatures and many of our metrics of capture success. Our three touchdown hybridization temperatures improved locus recovery for degraded, dried samples when compared to the fixed standard 65°C, but recovery did not appear to differ when comparing across the touchdown protocols. For samples with highly fragmented DNA, it may be that the hybridization temperature of baits to bind to short target DNA (shorter than DNA usually targeted from fresh samples) only requires a modest reduction to obtain the maximum amount of targeted data possible. Furthermore, slightly lowering hybridization temperature may have the benefit of recovering enough loci to ensure a taxon can be represented in a final analysis where missing data is minimized. Thus, based on our assessment, employing capture approaches with hybridization temperatures less than 65°C generally have positive outcomes, especially with samples of low quality.

## Conclusion

Our study primarily focused on the effects of a single target capture parameter, i.e., hybridization temperature, on several metrics of capture success in an invertebrate protein-coding dataset. However, given that our baits were derived from different genomic resources, we were also able to explore the role of bait-target divergence and bait tiling strategy in our experiment, as well as tissue quality. While hybridization temperature, bait-target divergence, and bait tiling strategy had effects on some of our metrics, we recognize other parameters likely affect capture efficacy and remain to be thoroughly investigated in target capture experiments similar to ours (i.e., in-solution capture, sample quality, etc.). Such parameters may also include base genomes used in probe design (Gustafson et al., 2019), GC content of baits (Tewhey et al., 2009; Ávila-Arcos et al., 2011), amount of baits used (Cruz-Dávalos et al., 2017), and post-hybridization washing stringency (Li et al., 2013), among many others. Regardless, our study suggests that lowering the hybridization temperature during capture can be beneficial to similar studies (especially those with high bait-target divergences) seeking to improve target recovery in fresh and degraded invertebrate material without major negative impacts overall, while also retrieving other potentially useful data for comparative analyses.

## Acknowledgements

We thank Joe Eger, Christiane Weirauch, and the Field Museum of Natural History for providing new samples for our study. We also thank Kevin Johnson for kindly providing transcriptome data prior to their publication. Caroline Miller and Min Zhao assisted with molecular benchwork. Gavin Naylor and Shannon Corrigan kindly provided use of their Covaris ultrasonication. Edward L. Braun helped with some perl and python scripts used in this study, and Ke Bi modified the Portik et al. (2016) scripts in order to accommodate our data. We also thank Daniela Wilner and Ellen Humbel for their advice and resources for the statistical analyses. Christine W. Miller and members of the Miller lab provided feedback on early versions of the manuscript. This study was funded by the National Science Foundation Grant IOSD1553100 (awarded to Christine W. Miller).

## Data availability

Newly generated sequence read files are available on NCBI’s Sequence Read Archive (SRA) under BioProject PRJNA763985. For previously published sequence read data, see NCBI SRA BioProjects PRJNA531965 (Forthman et al., 2019), PRJNA546248 (Forthman et al., 2020), and PRJNA609116 (Emberts et al., 2020).

## Supplementary material

**Appendix 1.** Effect of hybridization temperature on legacy locus recovery from off-target target capture data.

Table S1. Information regarding sample age, preservation method, and DNA extraction and library preparation protocols used. Abbreviations: DNeasy, Qiagen, DNeasy Blood and Tissue Kit; DNQIA, DNeasy with Qiagen QIAquick PCR Purification Kit.

Table S2. Target capture experimental design. Freshly preserved samples or samples preserved dried were subjected to the standard and TD-60 protocols, respectively, prior to the start of this study. Abbreviations: TD-60, touchdown hybridization approach starting at 65°C for 12 hrs followed by 62°C for 12 hrs and ending at 60°C for 12 hrs; TD-55, touchdown hybridization approach starting at 65°C for 12 hrs followed by 60°C for 12 hrs and ending at 55°C for 12 hrs; TD-50, touchdown hybridization approach starting at 65°C for 9 hrs followed by 60°C for 9 hrs, 55°C for 9 hrs, and ending at 50°C for 9 hrs.

Table S3. Summary data for raw and filtered sequence reads, reads on-target and read depth across all argeted loci, and contigs. Abbreviations: FRO, filtered reads on-target; see Table S2 for additional abbreviations.

Table S4. Summary data for on-target reads and read depth of targeted loci, partitioned based on type of baits used. Abbreviations: C-baits, loci targeted by exon- or transcript-derived baits (i.e., Coreoidea-derived baits); exon, loci targeted by exon-derived baits; P-baits, loci targeted only by Pentatomomorpha-derived UCE baits; PC dual, loci targeted by both Pentatomomorpha ultraconserved element (UCE) baits and Coreoidea-derived baits (i.e., Pentatomomorpha-Coreoidea dual baits); RD, read depth; transcript, loci argeted by transcript-derived baits; see Tables S2 and S3 and Table 1 of main text for explanation of erms used.

Table S5. Summary data for captured loci targeted by exon-derived baits (376 loci targeted). Abbreviations: see Tables S2–S4.

Table S6. Summary data for captured loci targeted by transcript-derived baits (58 loci targeted). Abbreviations: see Tables S2–S4.

Table S7. Summary data for captured loci targeted by Pentatomomorpha-Coreoidea dual baits (103 loci argeted). Abbreviations: see Tables S2–S4.

Table S8. Summary data for captured loci targeted by Pentatomomorpha-derived baits (2566 loci argeted). Abbreviations: see Tables S2–S4.

Table S9. Summary data for raw and filtered sequence reads, reads on-target and read depth across all argeted loci, and contigs from 24 taxa that had 2,000,000 million raw reads subsampled to equalize sequencing depth across capture conditions. Abbreviations: see Tables S2 and S3.

Table S10. Summary data for on-target reads and read depth of targeted loci, partitioned based on type of baits used for 24 taxa that had 2,000,000 million raw reads subsampled to equalize sequencing depth across capture conditions. Abbreviations: see Tables S2–S4 and Table 1 of main text.

Table S11. Summary data for captured loci targeted by exon-derived baits (376 loci targeted) from 24 taxa hat had 2,000,000 million raw reads subsampled to equalize sequencing depth across capture conditions. Abbreviations: see Tables S2–S4.

Table S12. Summary data for captured loci targeted by transcript-derived baits (58 loci targeted) from 24 axa that had 2,000,000 million raw reads subsampled to equalize sequencing depth across capture conditions. Abbreviations: see Tables S2–S4.

Table S13. Summary data for captured loci targeted by Pentatomomorpha-Coreoidea dual baits (103 loci argeted) from 24 taxa that had 2,000,000 million raw reads subsampled to equalize sequencing depth across capture conditions. Abbreviations: see Tables S2–S4.

Table S14. Summary data for captured loci targeted by Pentatomomorpha-derived baits (2566 loci argeted) from 24 taxa that had 2,000,000 million raw reads subsampled to equalize sequencing depth across capture conditions. Abbreviations: see Tables S2–S4.

Figure S1. Effects of target capture protocols on the (A) total number of raw reads, (B) assembled contigs, and (C) median contig length, separated by preservation method. (D) Average minimum bait-target divergences by bait design strategy. Numbers in parentheses above x-axis denote sample size. Single, double, and triple asterisks denote statistically significant pairwise comparisons, with p < 0.050, p < 0.010, and p < 0.001, respectively (statistical analyses not performed on dried samples due to low sample sizes). See Tables S2–S4 for abbreviations.

Figure S2. Effects of target capture protocols on (A) the proportion of filtered reads on-target for loci argeted by exon-derived baits, (B) the proportion of filtered reads on-target for loci targeted by transcript-derived baits, (C) read depth across loci targeted by exon-derived baits, and (D) read depth across loci argeted by transcript-derived baits, separated by sample preservation method. Numbers in parentheses above x-axis denote sample size. Double and triple asterisks denote statistically significant pairwise comparisons, with p < 0.010 and p < 0.001, respectively (statistical analyses not performed on dried samples due to low sample sizes). See Tables S2–S4 for abbreviations.

Figure S3. Effects of target capture protocols on (A) the proportion of targeted loci captured by exon-derived baits, (B) the proportion of targeted loci captured by transcript-derived baits, (C) the median engths of loci targeted by exon-derived baits, and (D) the median lengths of loci targeted by transcript-derived baits, separated by sample preservation method. Numbers in parentheses above x-axis denote sample size. Single, double, and triple asterisks denote statistically significant pairwise comparisons, with p < 0.050, p < 0.010, and p < 0.001, respectively (statistical analyses not performed on dried samples due o low sample sizes). See Tables S2–S4 for abbreviations.

Figure S4. Effects of target capture protocols on the number of putative paralogs of loci captured (A) exon-derived and (B) transcript-derived baits, separated by sample preservation method. Numbers in parentheses above x-axis denote sample size. All pairwise comparisons are not significant (statistical analyses not performed on dried samples due to low sample sizes). See Tables S2–S4 for abbreviations.

Figure S5. Effects of target capture protocols on the proportion of filtered reads on-target (A) across all argeted loci, (B) loci captured by Coreoidea-derived baits, (C) loci captured by PC dual baits, and (D) oci captured by Pentatomomorpha-derived baits, separated by preservation method. Numbers in parentheses above x-axis denote sample size. Single, double, and triple asterisks denote statistically significant pairwise comparisons, with p < 0.050, p < 0.010, and p < 0.001, respectively (statistical analyses not performed on dried samples due to low sample sizes). See Tables S2–S4 for abbreviations.

Figure S6. Effects of target capture protocols on read depth (A) across all targeted loci, (B) loci captured by Coreoidea-derived baits, (C) loci captured by PC dual baits, and (D) loci captured by Pentatomomorpha-derived baits, separated by preservation method. Numbers in parentheses above x-axis denote sample size. Single and triple asterisks denote statistically significant pairwise comparisons, with p < 0.050 and p < 0.001, respectively (statistical analyses not performed on dried samples due to low sample sizes). See Tables S2–S4 for abbreviations.

Figure S7. Effects of target capture protocols on the proportion of recovered loci targeted by (A) Coreoidea-derived baits, (C) PC dual baits, and (D) Pentatomomorpha-derived baits, separated by preservation method. Numbers in parentheses above x-axis denote sample size. Single, double, and triple asterisks denote statistically significant pairwise comparisons, with p < 0.050, p < 0.010, and p < 0.001, espectively (statistical analyses not performed on dried samples due to low sample sizes). See Tables S2– S4 for abbreviations.

Figure S8. Effects of target capture protocols on the median length of recovered loci targeted by (A) Coreoidea-derived baits, (C) PC dual baits, and (D) Pentatomomorpha-derived baits, separated by preservation method. Numbers in parentheses above x-axis denote sample size. Double and triple asterisks denote statistically significant pairwise comparisons, with p < 0.010 and p < 0.001, respectively (statistical analyses not performed on dried samples due to low sample sizes). See Tables S2–S4 for abbreviations.

Figure S9. Effects of target capture protocols on the number of putative paralogs of loci captured by (A) Coreoidea-derived baits, (C) PC dual baits, and (D) Pentatomomorpha-derived baits, separated by preservation method. Numbers in parentheses above x-axis denote sample size. Triple asterisks denote statistically significant pairwise comparisons, with p < 0.001 (statistical analyses not performed on dried samples due to low sample sizes). See Tables S2–S4 for abbreviations.

Figure S10. Effects of tiling strategy (Coreoidea-derived ∼2x tiling density; Pentatomomorpha-derived ∼1.33x tiling density) on average read depth per locus (captured loci exhibit 0.05–0.10 average minimum bait-target divergences for each bait design strategy). Numbers in parentheses above x-axis denote sample size. Triple asterisks denote statistically significant pairwise comparisons, with p < 0.001. See Tables S2– S4 for abbreviations.

Figure S11. When controlling for sequencing depth, effects of target capture protocols on (A) the number of assembled contigs, (B) median contig length, (C) proportion of filtered reads on-target across all argeted loci, and (D) read depth across all targeted loci for samples preserved fresh. Numbers in parentheses above x-axis denote sample size. Single, double, and triple asterisks denote statistically significant pairwise comparisons, with p < 0.050, p < 0.010, and p < 0.001, respectively. See Tables S2– S4 for abbreviations.

Figure S12. When controlling for sequencing depth, effects of target capture protocols on the proportion of filtered reads on-target for loci targeted by (A) exon-derived baits, (B) transcript-derived baits, (C) PC dual baits, and (D) Pentatomomorpha-derived baits for samples preserved fresh. Numbers in parentheses above x-axis denote sample size. Double and triple asterisks denote statistically significant pairwise comparisons, with p < 0.010 and p < 0.001, respectively. See Tables S2–S4 for abbreviations.

Figure S13. When controlling for sequencing depth, effects of target capture protocols on the read depth or loci targeted by (A) exon-derived baits, (B) transcript-derived baits, (C) PC dual baits, and (D) Pentatomomorpha-derived baits for samples preserved fresh. Numbers in parentheses above x-axis denote sample size. Single, double, and triple asterisks denote statistically significant pairwise comparisons, with p < 0.050, p < 0.010, and p < 0.001, respectively. See Tables S2–S4 for abbreviations.

Figure S14. When controlling for sequencing depth, effects of target capture protocols on the proportion of loci captured by (A) exon-derived baits, (B) transcript-derived baits, (C) PC dual baits, and (D) Pentatomomorpha-derived baits for samples preserved fresh. Numbers in parentheses above x-axis denote sample size. Single, double, and triple asterisks denote statistically significant pairwise comparisons, with p < 0.050, p < 0.010, and p < 0.001, respectively. See Tables S2–S4 for abbreviations.

Figure S15. When controlling for sequencing depth, effects of target capture protocols on the median engths of loci captured by (A) exon-derived baits, (B) transcript-derived baits, (C) PC dual baits, and (D) Pentatomomorpha-derived baits for samples preserved fresh. Numbers in parentheses above x-axis denote sample size. Single, double, and triple asterisks denote statistically significant pairwise comparisons, with p < 0.050, p < 0.010, and p < 0.001, respectively. See Tables S2–S4 for abbreviations.

Figure S16. When controlling for sequencing depth, effects of target capture protocols on the number of putative paralogs of loci captured by (A) exon-derived baits, (B) transcript-derived baits, (C) PC dual baits, and (D) Pentatomomorpha-derived baits for samples preserved fresh. Numbers in parentheses above x-axis denote sample size. Single, double, and triple asterisks denote statistically significant pairwise comparisons, with p < 0.050, p < 0.010, and p < 0.001, respectively. See Tables S2–S4 for abbreviations.

## Notes

### Competing Interest Statement

The authors have declared no competing interest.

### Summary of Updates

Statistical analyses and section on novelty of study added; title updated; figures and tables revised; supplementary files updated.

