## Appendix 1 for "Low hybridization temperatures improve target capture success of invertebrate loci"

Appendix 1: Effect of hybridization temperature on legacy locus recovery from off-target sequence data

Michael Forthman^1,2*^, Eric R. L. Gordon^3^, Rebecca T. Kimball^4^

^1^ California State Collection of Arthropods, Plant Pest Diagnostics Branch, California Department of Food and Agriculture, 3294 Meadowview Road, Sacramento, CA 95832, USA

^2^ Entomology & Nematology Department, University of Florida, 1881 Natural Area Drive, Gainesville, FL 32611, USA

^3^ Department of Ecology & Evolutionary Biology, University of Connecticut, 75 N. Eagleville Road, Unit 3043, Storrs, CT 06269, USA

^4^ Department of Biology, University of Florida, 876 Newell Drive, Gainesville, FL 32611, USA

*Corresponding author: California State Collection of Arthropods, Plant Pest Diagnostics Branch, California Department of Food and Agriculture, 3294 Meadowview Road, Sacramento, CA 95832, USA;

Background

Lower hybridization temperatures relax specificity between baits and targets, which may increase the number of off-target sequences (e.g., Cruz-Dávalos et al., 2017) and thus reduce read numbers for targeted regions. However, off-targets reads may contain sequences from loci traditionally used in phylogenetic studies (herein referred to as “legacy loci”) (e.g., Amaral et al., 2015; Wang et al., 2017; Simon et al., 2019; Miller et al., 2022), and there has been recent interest in complementing sequence capture data with legacy loci (e.g., Blaimer et al., 2015; Branstetter et al., 2017; Derkarabetian et al., 2019; Simon et al., 2019; Zhang, Deng, et al. , 2019; Branstetter et al., 2021; Hughes et al., 2021). Integrating legacy loci with target capture datasets has benefits, such as increasing the resolution power for phylogenetic inference and the inclusion of rare species with existing legacy data that are difficult to sample repeatedly for molecular studies (Branstetter et al., 2017; Derkarabetian et al., 2019; Zhang, Williams, et al., 2019).

While legacy locus data can be integrated with target capture data by designing baits from legacy loci (Branstetter et al., 2017; Simon et al., 2019; Hughes et al., 2021), this may increase the cost of custom probe kits because more baits may be required across more species due to higher substitution rates of some loci (e.g., mtDNA) and/or these baits may be included in a separate kit to prevent high copy number loci, like those on the mtDNA genome or the rRNA operon, from dominating capture data (Ströher et al., 2016; Pierce et al., 2017; Allio et al., 2020; Branstetter et al., 2021; Miller et al., 2022). Extracting legacy loci from off-target reads can circumvent some of these issues (Miller et al., 2022), and they have been successfully integrated with capture data despite their often-fewer numbers (compared to, e.g., 1000+ UCE loci) and/or introduction of large amounts of missing data (e.g., Simon et al., 2019; Miller et al., 2022). Thus, having off-target reads may not always be detrimental in capture studies when designing legacy locus baits is less desirable, as long as the primary targeted regions (e.g., UCEs) are also recovered.

Material and methods

We extracted mtDNA and nuclear rRNA legacy loci from off-target contigs in our coreoid target capture dataset. Briefly, we retrieved sequence data for 15 mtDNA (13 protein-coding and two ribosomal regions) and two nuclear rRNA loci (18S and 28S) from the National Center for Biotechnology Information’s (NCBI) database. While several coreoid nuclear protein-coding loci have been published and made available on NCBI (Tian et al., 2011), a recent study found that these loci were recovered in exceptionally low numbers (in some cases, no loci were recovered) from off-target capture data (Miller et al., 2022), and thus, we excluded a search for these loci in our present study. We used MitoFinder v1.1 (Allio et al., 2020) to extract mtDNA sequences. To identify nuclear legacy loci, we created a local nucleotide database using BLAST for each locus of interest and queried our capture data against them using blastn (e-value set to 1 x 10^-50^). We then calculated the proportion of legacy loci recovered, as well as when sequencing depth was controlled (see main text for information on subsampling reads with Seqtk, which did not include dried samples). To test for statistically significant differences in the proportion of legacy loci recovered across capture conditions, we constructed a generalized linear mixed model (GLMMs) with a binomial error structure for freshly preserved samples (i.e., ethanol, frozen, or silica beads) in our entire dataset (with an observation-level random effect also included due to overdispersion); we excluded dried samples from analyses due to their exceptionally low sample size for some target capture protocols (but we highlight trends in Results). We also constructed a GLMM with a binomial error structure for freshly preserved samples in our subsampled dataset. We performed simultaneous tests for general linear hypotheses using Tukey contrasts for multiple comparisons of means.

Results

When comparing overall legacy locus recovery across target capture protocols, as well as in pairwise comparisons of protocols for each taxon, lower hybridization temperatures (50°C and 55°C) significantly improved legacy locus recovery from off-target data of freshly preserved samples (Table A1; Fig. A1). This was particularly true for mtDNA legacy loci, in which we observed a drastic increase in the number of loci recovered for most samples subjected to the TD-50 protocol instead of the standard one. While rRNA legacy loci appeared to be recovered more often at lower hybridization temperatures, this was not as consistent among samples compared to mtDNA legacy loci. For the dried samples, we did not observe any distinctive trends between touchdown capture protocols, but these protocols recovered legacy loci compared to the standard protocol (Fig. A1). For freshly preserved samples, we also found congruent patterns in legacy locus recovery when sequencing depth was controlled, with TD-50 being the most significant (Table A2; Fig. A2).

Discussion

While off-target data may be considered undesirable, lower hybridization temperatures during capture can provide a greater chance to extract more legacy loci than what might be acquired under standard capture conditions. This can be advantageous for phylogenomic studies that seek to generate more comprehensive datasets (taxon and character sampling) by integrating capture data with well-known markers of historical use in molecular phylogenetic studies but are constrained by the added costs of generating additional custom baits or sequencing genomes, even at low read depth. Several target capture studies, especially in vertebrates, have extracted legacy loci from off-target sequences in capture data (e.g., Meiklejohn et al., 2014; Amaral et al., 2015; Wang et al., 2017; Derkarabetian et al., 2019; Simon et al., 2019; Miller et al., 2022). Legacy locus data extracted for focal taxa can then be combined with loci retrieved from genetic depositories for other taxa of interest prior to phylogenetic analysis, as well as to assess possible errors and/or contamination. While the relatively low number of legacy loci available for taxa in depositories does not always provide enough information for perfect phylogenetic resolution, sometimes these loci are the only available data for rare taxa. If such taxa are closely related to others in a target capture experiment, they can be accurately placed in phylogenetic analyses despite the large amounts of missing data (e.g., Kieran et al., 2021; Miller et al., 2022). Given the sequencing depth of a standard Illumina HiSeq or NextSeq lane often used for sequencing target capture data and typical pooling of 60–96 samples for sequencing, the presence of some usable off-target data may be desirable compared to the limited range of on-target loci sequenced at very high, sometimes excessive, depth.

Table A1. Summary data for legacy loci (15 mitochondrial [mtDNA] and two nuclear ribosomal [rRNA] loci targeted). Abbreviations: mtDNA, mitochondrial DNA; rRNA, nuclear ribosomal DNA; TD-60, touchdown hybridization approach starting at 65°C for 12 hrs followed by 62°C for 12 hrs and ending at 60°C for 12 hrs; TD-55, touchdown hybridization approach starting at 65°C for 12 hrs followed by 60°C for 12 hrs and ending at 55°C for 12 hrs; TD-50, touchdown hybridization approach starting at 65°C for 9 hrs followed by 60°C for 9 hrs, 55°C for 9 hrs, and ending at 50°C for 9 hrs.

| **Taxon** | **Target capture protocol** | **No. mtDNA loci recovered** | **% mtDNA loci recovered** | **No. rRNA loci recovered** | **% rRNA loci recovered** |
| --- | --- | --- | --- | --- | --- |
| *Acanthocoris sordidus* | Standard | 12 | 80.00 | 0 | 0.00 |
|  | TD-50 | 13 | 86.67 | 2 | 100.00 |
| *Anasa scorbutica* | Standard | 9 | 60.00 | 2 | 100..00 |
|  | TD-60 | 9 | 60.00 | 2 | 100.00 |
| *Anasa varicornis* | Standard | 1 | 6.67 | 1 | 50.00 |
|  | TD-60 | 1 | 6.67 | 2 | 100.00 |
| *Anisoscelis gradadius* | Standard | 0 | 0.00 | 2 | 100.00 |
|  | TD-60 | 2 | 13.33 | 2 | 100.00 |
| *Anoplocnemis curvipes* | Standard | 6 | 40.00 | 2 | 100.00 |
|  | TD-50 | 15 | 100.00 | 2 | 100.00 |
| *Catorhintha texana* | Standard | 0 | 0.00 | 0 | 0.00 |
|  | TD-55 | 9 | 60.00 | 2 | 100.00 |
| *Cebrenis supina* | Standard | 0 | 0.00 | 0 | 0.00 |
|  | TD-60 | 14 | 93.33 | 1 | 50.00 |
| *Chariesterus antennator* | Standard | 2 | 13.33 | 0 | 0.00 |
|  | TD-60 | 2 | 13.33 | 1 | 50.00 |
| *Chelinidea vittiger* | Standard | 3 | 20.00 | 2 | 100.00 |
|  | TD-55 | 11 | 73.33 | 2 | 100.00 |
| *Cletus ochraceus* | Standard | 5 | 33.33 | 0 | 0.00 |
|  | TD-55 | 14 | 93.33 | 2 | 100.00 |
| *Darmistus* sp. | Standard | 1 | 6.67 | 2 | 100.00 |
|  | TD-50 | 13 | 86.67 | 2 | 100.00 |
| Dasynini sp. | Standard | 2 | 13.33 | 0 | 0.00 |
|  | TD-50 | 15 | 100.00 | 2 | 100.00 |
| *Dysdercus mimus* | Standard | 3 | 20.00 | 2 | 100.00 |
|  | TD-50 | 15 | 100.00 | 2 | 100.00 |
| *Dysdercus suturellus* | Standard | 2 | 13.33 | 2 | 100.00 |
|  | TD-60 | 2 | 13.33 | 2 | 100.00 |
| *Holhymenia* sp. | Standard | 2 | 13.33 | 2 | 100.00 |
|  | TD-55 | 14 | 93.33 | 2 | 100.00 |
| *Hypselonotus bitrianguliger* | Standard | 1 | 6.67 | 1 | 50.00 |
|  | TD-55 | 12 | 80.00 | 2 | 100.00 |
| *Hypselonotus lineatus* | Standard | 0 | 0.00 | 2 | 100.00 |
|  | TD-55 | 4 | 26.67 | 2 | 100.00 |
| *Laminiceps festivus* | TD-60 | 6 | 40.00 | 1 | 50.00 |
|  | TD-55 | 4 | 26.67 | 1 | 50.00 |
| *Laminiceps obscurior* | TD-60 | 15 | 100.00 | 1 | 50.00 |
|  | TD-50 | 15 | 100.00 | 2 | 100.00 |
| *Largus* sp. | Standard | 3 | 20.00 | 2 | 100.00 |
|  | TD-55 | 9 | 60.00 | 2 | 100.00 |
| *Leptoglossus clypealis* | Standard | 1 | 6.67 | 2 | 100.00 |
|  | TD-60 | 0 | 0.00 | 2 | 100.00 |
| *Leptoscelis quadrisignatus* | Standard | 0 | 0.00 | 2 | 100.00 |
|  | TD-60 | 2 | 13.33 | 2 | 100.00 |
| *Melanacanthus margineguttatus* | Standard | 9 | 60.00 | 2 | 100.00 |
|  | TD-60 | 3 | 20.00 | 2 | 100.00 |
| *Myla* sp. | Standard | 1 | 6.67 | 2 | 100.00 |
|  | TD-55 | 9 | 60.00 | 2 | 100.00 |
| *Nematopus lepidus* | Standard | 3 | 20.00 | 1 | 50.00 |
|  | TD-55 | 8 | 53.33 | 2 | 100.00 |
| *Neomegalotomus rufipes* | Standard | 5 | 33.33 | 2 | 100.00 |
|  | TD-60 | 5 | 33.33 | 2 | 100.00 |
| *Omanocoris versicolor* | Standard | NA | NA | NA | NA |
|  | TD-60 | 0 | 0.00 | 0 | 0.00 |
| *Paralycambes pronotalis* | TD-60 | 15 | 100.00 | 2 | 100.00 |
|  | TD-55 | 15 | 100.00 | 2 | 100.00 |
| *Petascelis remipes* | Standard | 11 | 73.33 | 2 | 100.00 |
|  | TD-50 | 15 | 100.00 | 2 | 100.00 |
| *Phthia lunata* | Standard | 0 | 0.00 | 0 | 0.00 |
|  | TD-60 | 0 | 0.00 | 2 | 100.00 |
| *Phthiacnemia picta* | Standard | 2 | 13.33 | 2 | 100.00 |
|  | TD-50 | 15 | 100.00 | 2 | 100.00 |
| *Physomerus grossipes* | Standard | 2 | 13.33 | 0 | 0.00 |
|  | TD-50 | 15 | 100.00 | 2 | 100.00 |
| *Plapigus abdominalis* | Standard | 1 | 6.67 | 1 | 50.00 |
|  | TD-50 | 15 | 100.00 | 2 | 100.00 |
| *Plectropoda* sp. | Standard | 2 | 13.33 | 2 | 100.00 |
|  | TD-55 | 11 | 73.33 | 2 | 100.00 |
| *Salapia nigra* | TD-60 | 15 | 100.00 | 2 | 100.00 |
|  | TD-50 | 15 | 100.00 | 2 | 100.00 |
| *Sciophyrella neodiminuta* | TD-60 | 1 | 6.67 | 2 | 100.00 |
|  | TD-50 | 7 | 46.67 | 2 | 100.00 |
| *Spartocera fusca* | Standard | 10 | 66.67 | 2 | 100.00 |
|  | TD-60 | 5 | 33.33 | 1 | 50.00 |
| *Sphictyrtus pretiosus* | TD-60 | 12 | 80.00 | 2 | 100.00 |
|  | TD-55 | 12 | 80.00 | 2 | 100.00 |
| *Stenocoris* sp. | Standard | 7 | 46.67 | 1 | 50.00 |
|  | TD-50 | 15 | 100.00 | 2 | 100.00 |

Table A2. Summary data for legacy loci (15 mtDNA and two rRNA loci targeted) from 24 taxa that had 2,000,000 million raw reads subsampled to equalize sequencing depth across capture conditions. Abbreviations: see Table A1.

| **Taxon** | **Target capture protocol** | **No. mtDNA loci recovered** | **% mtDNA loci recovered** | **No. rRNA loci recovered** | **% rRNA loci recovered** |
| --- | --- | --- | --- | --- | --- |
| *Acanthocoris sordidus* | Standard | 12 | 80.00 | 0 | 0.00 |
|  | TD-50 | 15 | 100.00 | 2 | 100.00 |
| *Anasa scorbutica* | Standard | 6 | 40.00 | 1 | 50.00 |
|  | TD-60 | 5 | 33.33 | 2 | 100.00 |
| *Anisoscelis gradadius* | Standard | 0 | 0.00 | 1 | 50.00 |
|  | TD-60 | 1 | 6.67 | 2 | 100.00 |
| *Anoplocnemis curvipes* | Standard | 3 | 20.00 | 2 | 100.00 |
|  | TD-50 | 7 | 46.67 | 2 | 100.00 |
| *Catorhintha texana* | Standard | 0 | 0.00 | 0 | 0.00 |
|  | TD-55 | 6 | 40.00 | 2 | 100.00 |
| *Chelinidea vittiger* | Standard | 2 | 13.33 | 1 | 50.00 |
|  | TD-55 | 6 | 40.00 | 2 | 100.00 |
| *Darmistus* sp. | Standard | 1 | 6.67 | 2 | 100.00 |
|  | TD-50 | 2 | 13.33 | 2 | 100.00 |
| Dasynini sp. | Standard | 1 | 6.67 | 0 | 0.00 |
|  | TD-50 | 14 | 93.33 | 2 | 100.00 |
| *Dysdercus suturellus* | Standard | 1 | 6.67 | 2 | 100.00 |
|  | TD-60 | 1 | 6.67 | 2 | 100.00 |
| *Holhymenia* sp. | Standard | 1 | 6.67 | 2 | 100.00 |
|  | TD-55 | 5 | 33.33 | 2 | 100.00 |
| *Hypselonotus bitrianguliger* | Standard | 1 | 6.67 | 0 | 0.00 |
|  | TD-55 | 6 | 40.00 | 2 | 100.00 |
| *Hypselonotus lineatus* | Standard | 0 | 0.00 | 2 | 100.00 |
|  | TD-55 | 3 | 20.00 | 2 | 100.00 |
| *Leptoglossus clypealis* | Standard | 1 | 6.67 | 2 | 100.00 |
|  | TD-60 | 0 | 0.00 | 2 | 100.00 |
| *Leptoscelis quadrisignatus* | Standard | 0 | 0.00 | 2 | 100.00 |
|  | TD-60 | 0 | 0.00 | 2 | 100.00 |
| *Melanacanthus margineguttatus* | Standard | 3 | 20.00 | 2 | 100.00 |
|  | TD-60 | 2 | 13.33 | 2 | 100.00 |
| *Myla* sp. | Standard | 1 | 6.67 | 2 | 100.00 |
|  | TD-55 | 2 | 13.33 | 2 | 100.00 |
| *Nematopus lepidus* | Standard | 3 | 20.00 | 1 | 50.00 |
|  | TD-55 | 6 | 40.00 | 2 | 100.00 |
| *Neomegalotomus rufipes* | Standard | 1 | 6.67 | 2 | 100.00 |
|  | TD-60 | 1 | 6.67 | 2 | 100.00 |
| *Petascelis remipes* | Standard | 0 | 0.00 | 2 | 100.00 |
|  | TD-50 | 11 | 73.33 | 2 | 100.00 |
| *Physomerus grossipes* | Standard | 2 | 13.33 | 0 | 0.00 |
|  | TD-50 | 15 | 100.00 | 2 | 100.00 |
| *Plapigus abdominalis* | Standard | 1 | 6.67 | 1 | 50.00 |
|  | TD-50 | 12 | 80.00 | 2 | 100.00 |
| *Plectropoda* sp. | Standard | 1 | 6.67 | 2 | 100.00 |
|  | TD-55 | 3 | 20.00 | 2 | 100.00 |
| *Spartocera fusca* | Standard | 6 | 40.00 | 2 | 100.00 |
|  | TD-60 | 2 | 13.33 | 1 | 50.00 |
| *Stenocoris* sp. | Standard | 6 | 40.00 | 1 | 50.00 |
|  | TD-50 | 13 | 86.67 | 2 | 100.00 |

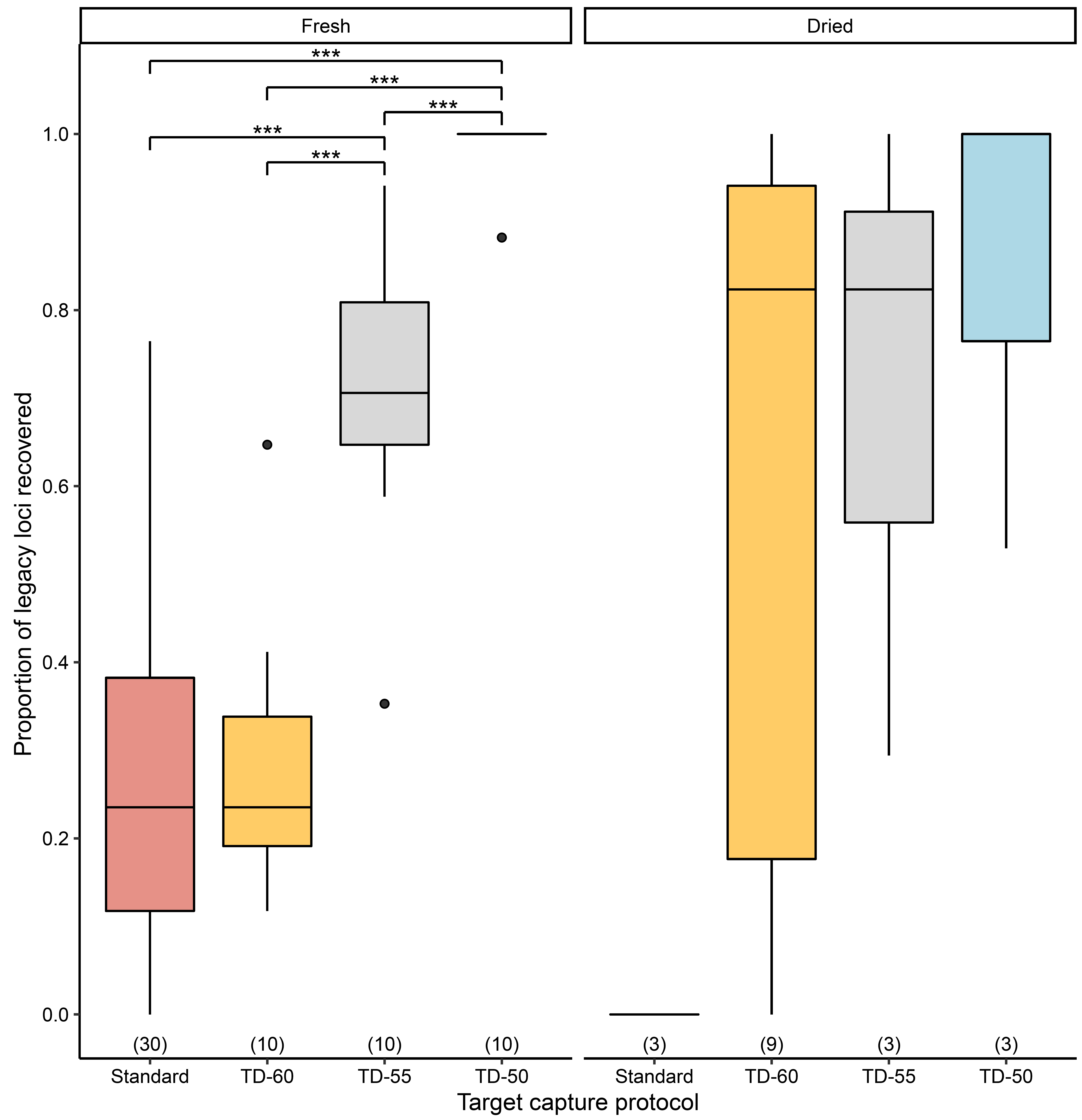

Figure A1. Effects of target capture protocols on the proportion of legacy loci recovered, separated by sample preservation method. Numbers in parentheses above x-axis denote sample size. Triple asterisks denote statistically significant pairwise comparisons, with p < 0.001 (statistical analyses not performed on dried samples due to low sample sizes). See Table A1 additional abbreviations.

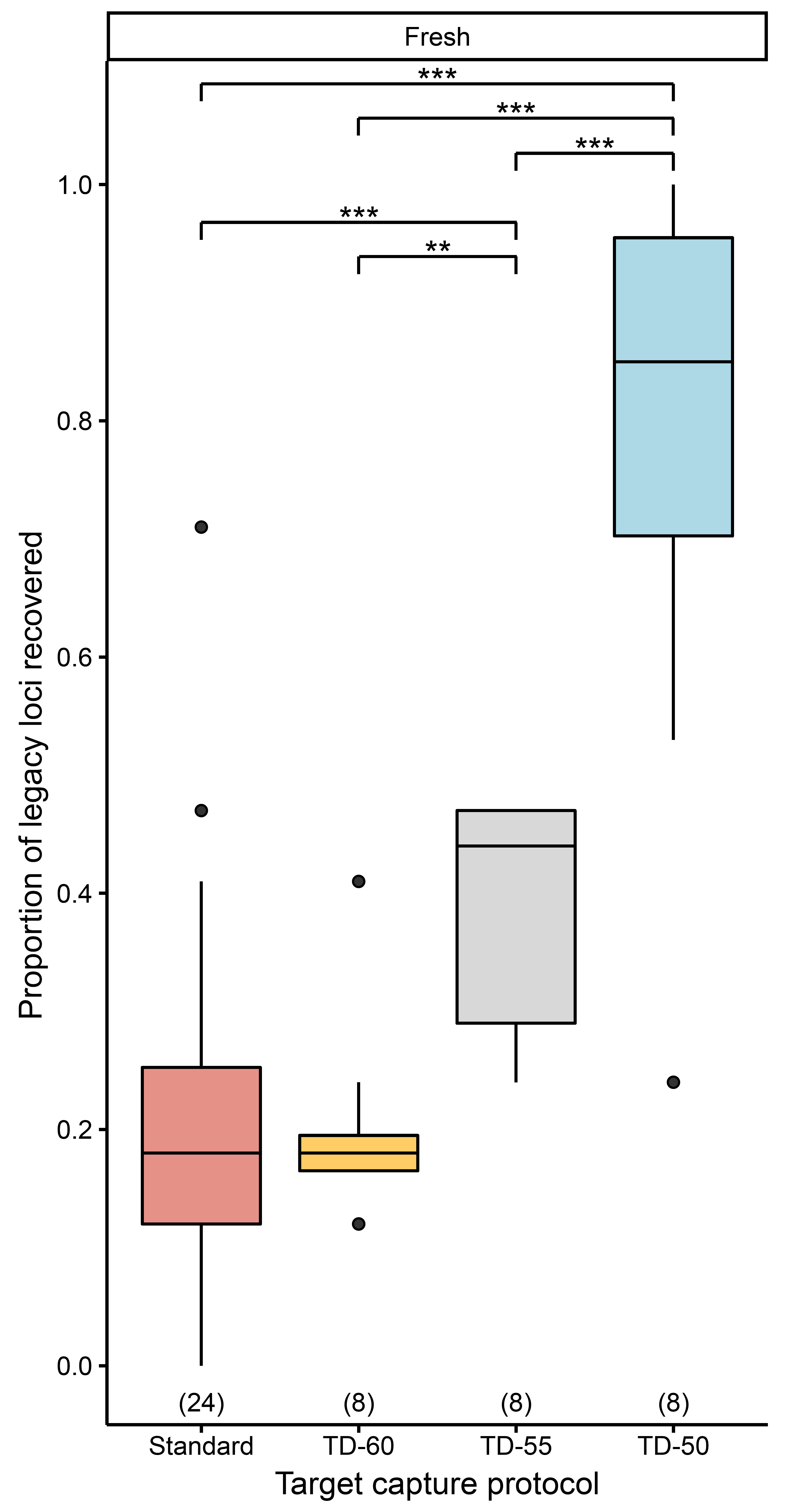

Figure A2. Effects of target capture protocols on the proportion of legacy loci recovered when controlling for sequencing depth for samples preserved fresh. Numbers in parentheses above x-axis denote sample size. Double and triple asterisks denote statistically significant pairwise comparisons, with p < 0.01 and p < 0.001, respectively. See Table A1 additional abbreviations.
