## Supplementary Methods, Tables, and Figures for "Low hybridization temperatures improve target capture success of invertebrate loci"

*Target capture baits*

We used our previously published custom myBaits kit (Forthman et al., 2019), which subsampled a Hemiptera-wide derived UCE bait set (at ~1.33x tiling) designed by Faircloth (2017) to only include two pentatomomorphan taxa that are more closely-related to but not included in our ingroup taxa (herein, referred to as “Pentatomomorpha-derived baits”; Table 2; Fig. 2). This kit also included an independently designed set of baits that were derived from coreoid transcriptomes, but these have not yet been introduced in the literature prior to this study. Thus, we introduce our coreoid bait design procedures here (herein, collectively referred to as “Coreoidea-derived baits”) and assess the effectiveness of these baits. Below, we describe two bait design strategies for our Coreoidea-derived baits, wherein baits were designed from individual exons while others were designed across entire transcripts.

For baits designed from individual coreoid exon sequences (herein, “exon-derived baits”, which is a subset of the Coreoidea-derived baits; Table 2; Fig. 2), we first retrieved an annotated draft genome of *Oncopeltus fasciatus* (Dallas, 1852) (Lygaeidae) from the Baylor College of Medicine – Human Genome Sequencing Center (https://www.hgsc.bcm.edu/arthropods/milkweed-bug-genome-project). We extracted exon sequences from the *O. fasciatus* genome using BEDTools v2.29.0 (Quinlan & Hall, 2010). Exon sequences were then filtered to exclude those that were <200 bp in length or that had a GC-content <30% or >70%.

We then obtained sequence reads for five published coreoid transcriptomes that have not been annotated: *Alydus pilosulus* Herrich-Schӓffer, 1847 (Johnson et al., 2018; NCBI BioProject PRJNA272214), *Anasa tristis* (De Geer, 1773) (Johnson et al., 2018; NCBI BioProject PRJNA272215), *Anoplocnemis curvipes* (Fabricius, 1781) (Agunbiade et al., 2013; NCBI BioProject PRJNA192258), *Boisea trivittata* (Say, 1825) (Johnson et al., 2018; NCBI BioProject PRJNA272221)*,* and *Clavigralla tomentosicollis* Stål, 1855 (Agunbiade et al., 2013; NCBI BioProject PRJNA192261). Sequence reads were processed using PRINSEQ-lite v0.20.4 (Schmieder & Edwards, 2011) and QuorUM v1.1.0 (Marçais et al., 2015), as well as *de novo* assembled in Trinity (Grabherr et al., 2011), following Forthman et al. (2019). For each coreoid transcriptome, a localized reciprocal blastn search using *O. fasciatus* individual exon sequences was performed (e-value threshold set to 1e-20 and percent identity to 60%).

The best reciprocal hit was extracted from each transcriptome, with sequences from multiple transcriptomes corresponding to the same exon grouped together in a single fasta file. These exons were then searched against all coreoid transcriptome sequences using blastn to confirm orthology and to identify additional sequences that may not have been found in reciprocal blast hits with the more distant *O. fasciatus* genome. We then used RepeatMasker (https://repeatmasker.org) (options: rmblast search engine and *Drosophila melanogaster* DNA source [all other options at default]) to exclude exons with low complexity sequences and simple repeats. Sequences were then aligned with MAFFT v7.305b (Katoh et al., 2002; Katoh & Standley, 2013) using the G-INS-i algorithm, and alignments were visually inspected in Geneious v9 to further confirm orthology. Sequences for a total of 456 individual exons were then selected for part of the Coreoidea-derived bait set.

We also used a pipeline from Portik et al. (2016) to design baits across transcript sequences (i.e., RNAseq-derived sequences that may include more than one exon, with baits potentially spanning across one or more introns; herein, “transcript-derived baits”; Table 2; Fig. 2). We first used Portik et al.’s (2016) 4-Annotation.pl script (with some modifications to process our data) and amino acid sequences of the *O. fasciatus* genome to annotate the assembled coreoid transcriptomes by transcript ID (e-value threshold set to 1e-20 and percent identity to 60%). We then used their 6-MarkerSelectionTRANS.pl script with default settings to find orthologous transcript sequences across our transcriptomes. This script requires an orthologous sequence to be present across all transcriptomes to be selected for bait design. Few transcript sequences were selected when all five of our transcriptomes were used because of this requirement; in attempting this, we found that the inclusion of the *B. trivittata* transcriptome was associated with the low number of transcript sequences selected. Thus, we excluded the *B. trivittata* transcriptome from subsequent searches. Furthermore, to maximize the number of transcript sequences selected for bait design while “allowing” for missing taxa, we performed two separate searches that excluded the *Ano. curvipes* or *Ana. tristis* transcriptomes, respectively. Transcript sequences were aligned and visually inspected as described above for exon-derived baits. Based on the annotations, 141 transcripts (out of 172 initially selected) did not correspond to any of the exon-derived baits. Of these, we selected 81 transcripts for part of the Coreoidea-derived baits that were found across most of our coreoid transcriptomes and that ranged between 256 bp and 1000 bp to reduce the number of baits potentially targeting multiple exons interspersed by long introns.

The final Coreoidea-derived bait sequences (i.e., exon- and transcript-derived baits) were submitted to Arbor Biosciences (Ann Arbor, MI) to produce 120 bp baits with ~2x tiling density. We also preliminarily compared our Coreoidea-derived baits against the Pentatomomorpha-derived baits using blastn (e-value threshold set to 1e-20) to determine whether any were potentially associated with a locus already targeted by the latter set of baits. Those Coreoidea-derived bait sequences matching to Pentatomomorpha-derived baits were not removed from the final selection of baits, as their inclusion could allow for some targeted loci to be captured by more baits from more closely-related species (herein, “Pentatomomorpha-Coreoidea [PC] dual baits”; Table 2; Fig. 2).

*DNA extraction and library preparation*

See references in Table S1 for details on library preparation and target capture for samples previously published. For new samples, genomic DNA was extracted from any part of the body or the entire body from EtOH-preserved, silica-bead preserved, frozen, or dried specimens to sample similar amounts of tissue across taxa, where possible (Table S1). Freshly preserved specimens were extracted with either the Gentra Puregene Tissue or Qiagen DNeasy Blood and Tissue kit (hereafter DNeasy) (Table S1). For the Puregene kit, we followed the manufacturer’s protocol for 5–10 mg tissue and optional recommendations, but we made the following modifications: 10 µL of proteinase K was added to samples; samples were incubated for 24–48 hr; 600 µL of 100% EtOH was used for the first wash, and the sample was then centrifuged for 10 mins; and 50–100 µL of molecular grade water or Puregene DNA Hydration Solution was used to resuspend isolated DNA. For the DNeasy kit, we also followed the manufacturer’s protocol but with fewer modifications: tissue was incubated in 180–190 µL Buffer ATL and 10–20 µL proteinase K for 24–48 hr, and depending on the source of the tissue, DNA was eluted once or twice with 50 µL Buffer AE.

For degraded museum specimens, DNA was extracted using a modified version of the DNeasy protocol, following Knyshov et al. (2019) (i.e., Qiagen DNeasy Blood and Tissue kit coupled with Qiagen QIAquick PCR purification kit; hereafter DNQIA) (Table S1). The protocol is designed to extract DNA >100 bp in length. The DNQIA protocol follows the DNeasy protocol to the first centrifugation step, but a QIAquick spin column is used. The samples are then subjected to the manufacturer’s Qiagen QIAquick PCR purification protocol by replacing AW1 and AW2 washes with PE buffer. Samples were then eluted in 30 µL EB buffer.

We assessed DNA quality and quantity with 1% agarose gel electrophoresis and a Qubit 2.0 fluorometer, respectively. Samples were normalized to 10–20 ng/µL. High molecular weight samples were then fragmented into 200–1000 bp using a Bioruptor UCD-300 sonication device (4–10 cycles of 30 s on/30 s) or a Covaris M220 Focused-ultrasonicator (20–60 s) (Table S1).

Libraries were constructed with a modified KAPA Hyper Prep Kit protocol following Forthman et al. (2019). Briefly, we used half volume reactions for all steps. iTru universal adapter stubs and 8 bp dual indexes were used (Glenn et al., 2019). Library amplification conditions involved initial denaturation at 98°C for 3 min; 14 cycles of 98°C for 30 s, 60°C for 30 s, and 72°C for 30 s; and a final extension at 72°C for 5 min. Amplified libraries quality and quantity were assess with gel electrophoresis and Qubit, respectively. Libraries were then combined into 1000 ng pools using equimolar amounts, dried at 60°C, and resuspended in 14 µL IDTE.

*Statistical analyses*

Statistical analyses using linear mixed models (LMMs), generalized linear models (GLMs), and generalized linear mixed models (GLMMs) were performed using *lme4* v1.1.30 (Bates et al., 2015) in R v4.1.2 (R Core Team, 2022). We excluded dried samples from statistical analyses due to very low sample size for some target capture protocols, with the exception that they were included with fresh samples when analyzing bait-target divergences since this was independent of target capture protocol. We refrained from treating library quality as a factor in analyses since this was used as an *a posteriori* criterion for selecting samples to be included in a second capture protocol. We also did not include the different sequencing efforts across samples as a random effect since it has been suggested that random effect terms should have at least five levels for inclusion (Harrison, 2015; but see Gomes, 2022). However, to account for potential effects of different sequencing efforts, we subsampled our data as described in the previous section to normalize sequencing depth across sequencing efforts. For LMM and GLMM analyses, we treated target capture protocol as a fixed effect. Because the same sample was subjected to two capture protocols, we included replication as a random effect. The GLM analyses treated bait design strategy as a fixed effect. For all analyses, we performed simultaneous tests for general linear hypotheses using Tukey contrasts for multiple comparisons of means using the *multcomp* v1.4.20 R package (Hothorn et al., 2008).

*Specific details of the statistical analyses for the entire dataset.* We conducted LMM analyses on read depth across all targeted loci regardless of bait design strategy, as well as when data were partitioned by bait design strategy. The response variable was log-transformed to achieve a normal distribution.

The response variables for GLM analyses were the average minimum bait-target divergences and the average read depth per locus across loci with low bait-target divergences (i.e., tiling strategy), with the latter log-transformed to achieve a more normal distribution. We used a binomial error structure with the average minimum bait-target divergences, while a Gaussian structure was used for the analysis investigating bait tiling strategy.

For GLMM analyses, we used a Poisson error structure with the following response variables: median lengths of loci targeted by Pentatomomorpha-derived baits and number of putative paralogs of loci targeted by transcript- and Pentatomomorpha-derived baits, as well as by PC dual baits. Due to evidence of overdispersion when using a Poisson error structure, we used the negative binomial family for the following response variables: total number of raw reads; total number of contigs; median contig length; median lengths of loci targeted by exon-, transcript-, and Coreoidea-derived baits, as well as PC dual baits; and number of putative paralogs of loci targeted by exon- and Coreoidea-derived baits. A binomial family was used for analyses involving the proportion of on-target reads (with an observation-level random effect due to evidence of overdispersed data [Williams, 1982]), proportion of target loci recovered (with an observation-level random effect for those targeted by exon-, Coreoidea, and Pentatomomorpha-derived baits), and proportion of legacy loci recovered.

*Specific details of the statistical analyses for the subsampled dataset.* For our dataset that subsampled reads, we performed statistical analyses similar to those described above for the entire dataset. For LMM analyses, only read depth data from the Pentatomomorpha-derived bait treatment were log-transformed to achieve a normal distribution (other treatments were normally distributed).

For GLMM analyses, many of our response variables were analyzed similarly as in the previous section. Here, we only highlight cases where four GLMMs were constructed slightly differently. First, we did not include an observation-level random effect for GLMMs with the proportion of target loci recovered by exon- and Coreoidea-derived baits, as there was no evidence of overdispersed data. Second—and for similar reasoning—, we used a Poisson error structure with the number of putative paralogs of loci targeted by exon- and Coreoidea-derived baits.

Table S1. Information regarding sample age, preservation method, and DNA extraction and library preparation protocols used. Abbreviations: DNeasy, Qiagen, DNeasy Blood and Tissue Kit; DNQIA, DNeasy with Qiagen QIAquick PCR Purification Kit.

| **Taxon** | **Collection year** | **Preservation method** | **Sampled structures** | **DNA extraction method** | **Shearing method** | **PreCR treatment** | **Reference, if published** |
| --- | --- | --- | --- | --- | --- | --- | --- |
| *Acanthocoris sordidus* CMF_0276 | 2016 | ethanol | whole body | Puregene | Bioruptor | No | Forthman et al. (2020) |
| *Anasa scorbutica* CMF_0169 | 2016 | ethanol | legs, abdomen | Puregene | Bioruptor | No | Emberts et al. (2020) |
| *Anasa varicornis* CMF_0005 | 2013 | ethanol | legs, abdomen | Puregene | Bioruptor | No | Forthman et al. (2020) |
| *Anisoscelis gradadius* CMF_0137 | 2013 | ethanol | legs, abdomen | Puregene | Bioruptor | No | Forthman et al. (2020) |
| *Anoplocnemis curvipes* CMF_0332 | 2015 | frozen | legs | Puregene | Bioruptor | No | Emberts et al. (2020) |
| *Catorhintha texana* CMF_0070 | 2015 | ethanol | legs, abdomen | Puregene | Bioruptor | No | Forthman et al. (2020) |
| *Cebrenis supina* CMF_1027 | 2017 | dried | whole body | DNQIA | Covaris | Yes |  |
| *Chariesterus antennator* CMF_0103 | 2016 | ethanol | legs, abdomen | Puregene | Bioruptor | No | Forthman et al. (2020) |
| *Chelinidea vittiger* CMF_0220 | 2016 | ethanol | abdomen | DNeasy | Bioruptor | No | Forthman et al. (2020) |
| *Cletus ochraceus* CMF_0152 | 2008 | ethanol | whole body | Puregene | Bioruptor | No | Forthman et al. (2020) |
| *Darmistus* sp. CMF_0161 | 2013 | ethanol | whole body | DNeasy | Bioruptor | No |  |
| Dasynini sp. CMF_0065 | 2013 | ethanol | abdomen | Puregene | Bioruptor | No |  |
| *Dysdercus mimus* CMF_0110 | 2009 | ethanol | whole body | DNeasy | Bioruptor | No | Forthman et al. (2019) |
| *Dysdercus suturellus* CMF_0305 | 2016 | ethanol | abdomen | DNeasy | Bioruptor | No | Forthman et al. (2019) |
| *Holhymenia* sp. CMF_0051 | 2010 | ethanol | legs | Puregene | Bioruptor | No | Forthman et al. (2020) |
| *Hypselonotus bitrianguliger* CMF_0073 | 2014 | ethanol | legs, abdomen | Puregene | Bioruptor | No | Forthman et al. (2020) |
| *Hypselonotus lineatus* CMF_0027 | 2010 | ethanol | legs, abdomen | Puregene | Bioruptor | No | Emberts et al. (2020) |
| *Laminiceps festivus* CMF_0605 | 1961 | dried | thorax, abdomen | DNQIA | Covaris | Yes |  |
| *Laminiceps obscurior* CMF_1020 | 2017 | dried | thorax, legs, abdomen | DNQIA | Covaris | Yes |  |
| *Largus* sp. CMF_0230 | 2016 | ethanol | whole body | DNeasy | Bioruptor | No | Forthman et al. (2019) |
| *Leptoglossus clypealis* CMF_0463 | 2016 | ethanol | legs, abdomen | DNeasy | Bioruptor | No | Forthman et al. (2020) |
| *Leptoscelis quadrisignatus* CMF_0076 | 2008 | ethanol | legs | Puregene | Bioruptor | No | Emberts et al. (2020) |
| *Melanacanthus margineguttatus* CMF_0418 | 2017 | ethanol | whole body | DNeasy | Bioruptor | No | Emberts et al. (2020) |
| *Myla* sp. CMF_0091 | 2008 | ethanol | whole body | DNeasy | Bioruptor | No | Forthman et al. (2019) |
| *Nematopus lepidus* CMF_0071 | 2013 | ethanol | legs | Puregene | Bioruptor | No | Forthman et al. (2020) |
| *Neomegalotomus rufipes* CMF_0098 | 2016 | ethanol | whole body | DNeasy | Bioruptor | No | Forthman et al. (2019) |
| *Omanocoris versicolor* CMF_1081 | 1935 | dried | whole body | DNQIA | Covaris | Yes |  |
| *Paralycambes pronotalis* CMF_1018 | 2017 | dried | thorax, legs, abdomen | DNQIA | Covaris | Yes |  |
| *Petascelis remipes* CMF_0330 | 2015 | silica beads | legs | Puregene | Bioruptor | No | Forthman et al. (2020) |
| *Phthia lunata* CMF_1085 | 1961 | dried | whole body | DNQIA | Covaris | Yes |  |
| *Phthiacnemia picta* CMF_0350 | 2017 | ethanol | abdomen | DNeasy | Bioruptor | No | Forthman et al. (2020) |
| *Physomerus grossipes* CMF_0175 | 2016 | ethanol | legs | Puregene | Bioruptor | No | Emberts et al. (2020) |
| *Plapigus abdominalis* CMF_0002 | 2013 | ethanol | legs, abdomen | Puregene | Bioruptor | No | Forthman et al. (2020) |
| *Plectropoda* sp. CMF_0022 | 2013 | ethanol | legs | Puregene | Bioruptor | No | Forthman et al. (2020) |
| *Salapia nigra* CMF_0604 | 1980(?) | dried | thorax, abdomen | DNQIA | Covaris | Yes |  |
| *Sciophyrella neodiminuta* CMF_0608 | 1944 | dried | whole body | DNQIA | Covaris | Yes |  |
| *Spartocera fusca* CMF_0526 | 2017 | ethanol | head, thorax, legs | DNeasy | Bioruptor | No | Forthman et al. (2020) |
| *Sphictyrtus pretiosus* CMF_1082 | 1941 | dried | whole body | DNQIA | Covaris | Yes |  |
| *Stenocoris* sp. CMF_0315 | 2010 | ethanol | legs, abdomen | Puregene | Bioruptor | No |  |

Table S2. Target capture experimental design. Freshly preserved samples or samples preserved dried were subjected to the standard and TD-60 protocols, respectively, prior to the start of this study. Abbreviations: TD-60, touchdown hybridization approach starting at 65°C for 12 hrs followed by 62°C for 12 hrs and ending at 60°C for 12 hrs; TD-55, touchdown hybridization approach starting at 65°C for 12 hrs followed by 60°C for 12 hrs and ending at 55°C for 12 hrs; TD-50, touchdown hybridization approach starting at 65°C for 9 hrs followed by 60°C for 9 hrs, 55°C for 9 hrs, and ending at 50°C for 9 hrs.

|  |  | **Library quality** | | |
| --- | --- | --- | --- | --- |
| **Preservation method** | **Target capture protocol** | **Best** | **Moderate** | **Marginal** |
| Dried | Standard | *Cebrenis supina* | *Phthia lunata* | *Omanocoris versicolor* |
|  | TD-55 | *Sphictyrtus pretiosus* | *Paralycambes pronotalis* | *Laminiceps festivus* |
|  | TD-50 | *Salapia nigra* | *Sciophyrella neodiminuta* | *Laminiceps obscurior* |
| Fresh | TD-60 | *Leptoglossus clypealis* | *Leptoscelis quadrisignatus* | *Melanacanthus margineguttatus* |
|  |  | *Neomegalotomus rufipes* | *Anasa scorbutica* | *Anasa varicornis* |
|  |  | *Anisoscelis gradadius* | *Dysdercus suturellus* | *Chariesterus antennator* |
|  |  | *Spartocera fusca* |  |  |
|  | TD-55 | *Plectropoda* sp. | *Hypselonotus bitrianguliger* | *Nematopus lepidus* |
|  |  | *Myla* sp. | *Catorhintha texana* | *Largus* sp. |
|  |  | *Holhymenia* sp. | *Chelinidea vittiger* | *Hypselonotus lineatus* |
|  |  |  | *Cletus ochraceus* |  |
|  | TD-50 | *Petascelis remipes* | *Dysdercus mimus* | Dasynini sp. |
|  |  | *Stenocoris* sp. | *Anoplocnemis curvipes* | *Phthiacnemia picta* |
|  |  | *Acanthocoris sordidus* | *Darmistus* sp. | *Physomerus grossipes* |
|  |  |  |  | *Plapigus abdominalis* |

Table S3. Summary data for raw and filtered sequence reads, reads on-target and read depth across all targeted loci, and contigs. Abbreviations: FRO, filtered reads on-target; see Table S2 for additional abbreviations.

| **Taxon** | **Target capture protocol** | **No. raw reads** | **No. filtered reads** | **No. total FRO** | **% Total FRO** | **Overall read depth** | **No. contigs** | **Total bp across contigs** | **Mean contig length** | **Median contig length** | **Min contig length** | **Max contig length** |
| --- | --- | --- | --- | --- | --- | --- | --- | --- | --- | --- | --- | --- |
| *Acanthocoris sordidus* | Standard | 7212888 | 2702481 | 1117130 | 41.34 | 92.73 | 8549 | 3897031 | 455.85 | 347.0 | 56 | 3047 |
|  | TD-50 | 29890222 | 15287910 | 1468804 | 9.61 | 56.64 | 300406 | 122185175 | 406.73 | 315.0 | 55 | 9887 |
| *Anasa scorbutica* | Standard | 8552484 | 3240607 | 1532900 | 47.30 | 107.95 | 14554 | 4620980 | 317.51 | 211.0 | 56 | 4246 |
|  | TD-60 | 3549878 | 1851730 | 831849 | 44.92 | 49.05 | 18643 | 6545202 | 351.08 | 236.0 | 56 | 5909 |
| *Anasa varicornis* | Standard | 7434086 | 2719521 | 842488 | 30.98 | 92.71 | 11148 | 2599509 | 233.18 | 92.0 | 56 | 3063 |
|  | TD-60 | 2058368 | 1296037 | 543053 | 41.90 | 36.73 | 15173 | 5105065 | 336.46 | 228.0 | 56 | 4041 |
| *Anisoscelis gradadius* | Standard | 7341354 | 4125411 | 1766322 | 42.82 | 123.11 | 28775 | 7608953 | 264.43 | 137.0 | 56 | 3846 |
|  | TD-60 | 5865360 | 3812435 | 1715426 | 45.00 | 97.31 | 33179 | 10964894 | 330.48 | 245.0 | 56 | 4780 |
| *Anoplocnemis curvipes* | Standard | 8379830 | 3695282 | 1251889 | 33.88 | 87.04 | 20700 | 5535191 | 267.40 | 111.0 | 56 | 5356 |
|  | TD-50 | 32370506 | 17639724 | 1290744 | 7.32 | 45.41 | 474803 | 125890086 | 265.14 | 158.0 | 55 | 14720 |
| *Catorhintha texana* | Standard | 7861266 | 2423631 | 1165774 | 48.10 | 112.27 | 6093 | 2113821 | 346.93 | 208.0 | 56 | 3420 |
|  | TD-55 | 4877766 | 2558288 | 908590 | 35.52 | 48.53 | 29009 | 10696524 | 368.73 | 259.0 | 56 | 6022 |
| *Cebrenis supina* | Standard | 130 | 44 | 0 | 0.00 | NA | 1 | 61 | 61.00 | 61.0 | 61 | 61 |
|  | TD-60 | 6665212 | 4304088 | 935034 | 21.72 | 131.67 | 134483 | 11022179 | 81.96 | 60.0 | 56 | 5902 |
| *Chariesterus antennator* | Standard | 3157454 | 1651822 | 368838 | 22.33 | 49.58 | 19639 | 3007671 | 153.15 | 73.0 | 56 | 2757 |
|  | TD-60 | 1938478 | 1280855 | 391606 | 30.57 | 35.41 | 20960 | 4555210 | 217.33 | 114.0 | 56 | 2827 |
| *Chelinidea vittiger* | Standard | 4606720 | 2052586 | 776013 | 37.81 | 67.54 | 13020 | 3737366 | 287.05 | 206.0 | 56 | 3029 |
|  | TD-55 | 5949378 | 3482097 | 831442 | 23.88 | 50.50 | 62187 | 15563883 | 250.28 | 213.0 | 56 | 4834 |
| *Cletus ochraceus* | Standard | 2987604 | 1571786 | 328254 | 20.88 | 35.33 | 19249 | 3927816 | 204.05 | 87.0 | 56 | 3452 |
|  | TD-55 | 11472838 | 7054064 | 823951 | 11.68 | 46.35 | 119278 | 29388020 | 246.38 | 138.0 | 56 | 7963 |
| *Darmistus* sp. | Standard | 6918570 | 4415144 | 431757 | 9.78 | 44.87 | 40646 | 5299412 | 130.38 | 66.0 | 56 | 8531 |
|  | TD-50 | 22240678 | 16105203 | 909446 | 5.65 | 47.09 | 353899 | 71189520 | 201.16 | 97.0 | 56 | 20305 |
| Dasynini sp. | Standard | 4518480 | 1710378 | 380760 | 22.26 | 62.48 | 11457 | 1914810 | 167.13 | 78.0 | 56 | 2707 |
|  | TD-50 | 5999540 | 3909865 | 469759 | 12.02 | 32.82 | 99579 | 24229291 | 243.32 | 174.0 | 56 | 7757 |
| *Dysdercus mimus* | Standard | 1820144 | 820915 | 272415 | 33.18 | 31.87 | 5841 | 2285316 | 391.25 | 276.0 | 56 | 4563 |
|  | TD-50 | 18983254 | 9850119 | 964278 | 9.79 | 39.68 | 198231 | 76186448 | 384.33 | 280.0 | 55 | 15330 |
| *Dysdercus suturellus* | Standard | 4146928 | 1823353 | 634728 | 34.81 | 59.43 | 9507 | 4103624 | 431.64 | 312.0 | 56 | 4171 |
|  | TD-60 | 4146154 | 2480417 | 1001799 | 40.39 | 57.91 | 23742 | 10796355 | 454.74 | 348.0 | 56 | 4281 |
| *Holhymenia* sp. | Standard | 5937966 | 3065667 | 1257403 | 41.02 | 85.19 | 22537 | 6160010 | 273.33 | 121.0 | 56 | 4675 |
|  | TD-55 | 12593576 | 7177770 | 1670268 | 23.27 | 75.45 | 100000 | 27356654 | 273.57 | 208.0 | 56 | 6504 |
| *Hypselonotus bitrianguliger* | Standard | 4697458 | 1953506 | 759963 | 38.90 | 65.65 | 11363 | 3324326 | 292.56 | 144.0 | 56 | 6231 |
|  | TD-55 | 7963868 | 4434477 | 945007 | 21.31 | 51.23 | 65629 | 18975279 | 289.13 | 224.0 | 56 | 9993 |
| *Hypselonotus lineatus* | Standard | 5723868 | 2286214 | 578938 | 25.32 | 76.21 | 13670 | 2429936 | 177.76 | 73.0 | 56 | 3349 |
|  | TD-55 | 3603512 | 2035392 | 407527 | 20.02 | 28.45 | 34428 | 9641335 | 280.04 | 210.0 | 56 | 6558 |
| *Laminiceps festivus* | TD-60 | 738862 | 264053 | 12963 | 4.91 | 26.99 | 3271 | 447833 | 136.91 | 80.0 | 56 | 1157 |
|  | TD-55 | 3499418 | 635726 | 23272 | 3.66 | 64.55 | 3123 | 488025 | 156.27 | 109.0 | 56 | 2955 |
| *Laminiceps obscurior* | TD-60 | 867644 | 345587 | 7124 | 2.06 | 15.23 | 3774 | 886008 | 234.77 | 220.0 | 56 | 15628 |
|  | TD-50 | 9535370 | 2965437 | 15510 | 0.52 | 24.41 | 16205 | 4951075 | 305.53 | 228.0 | 56 | 32319 |
| *Largus* sp. | Standard | 2219122 | 1014223 | 219662 | 21.66 | 29.04 | 11008 | 2415502 | 219.43 | 85.0 | 56 | 5370 |
|  | TD-55 | 8697084 | 4782325 | 618750 | 12.94 | 35.45 | 78837 | 23185792 | 294.10 | 221.0 | 56 | 14241 |
| *Leptoglossus clypealis* | Standard | 7089366 | 3349221 | 1359592 | 40.59 | 99.22 | 22303 | 5201335 | 233.21 | 100.0 | 56 | 3709 |
|  | TD-60 | 3682732 | 2313350 | 983901 | 42.53 | 60.15 | 25021 | 7342952 | 293.47 | 215.0 | 56 | 4825 |
| *Leptoscelis quadrisignatus* | Standard | 8373644 | 3481066 | 1595665 | 45.84 | 106.84 | 20646 | 5111669 | 247.59 | 98.0 | 56 | 4258 |
|  | TD-60 | 6842486 | 3907501 | 1858093 | 47.55 | 86.12 | 35537 | 11348726 | 319.35 | 223.0 | 56 | 4890 |
| *Melanacanthus margineguttatus* | Standard | 8421192 | 4104664 | 1158070 | 28.21 | 93.95 | 23027 | 5614154 | 243.81 | 94.0 | 56 | 3990 |
|  | TD-60 | 4358098 | 2736799 | 1132212 | 41.37 | 66.67 | 22821 | 7959814 | 348.79 | 255.0 | 56 | 5712 |
| *Myla* sp. | Standard | 6619040 | 2888756 | 918226 | 31.79 | 70.73 | 14087 | 4602614 | 326.73 | 213.0 | 56 | 3295 |
|  | TD-55 | 14281326 | 7824872 | 1845911 | 23.59 | 78.21 | 102490 | 33884626 | 330.61 | 252.0 | 56 | 7137 |
| *Nematopus lepidus* | Standard | 10602714 | 5778383 | 508195 | 8.80 | 73.96 | 62272 | 6557227 | 105.30 | 64.0 | 56 | 3187 |
|  | TD-55 | 4818490 | 3006296 | 484931 | 16.13 | 34.98 | 53635 | 12121257 | 226.00 | 121.0 | 56 | 5438 |
| *Neomegalotomus rufipes* | Standard | 13121288 | 7247447 | 1508737 | 20.82 | 127.35 | 63861 | 8773229 | 137.38 | 66.0 | 56 | 3950 |
|  | TD-60 | 6803472 | 4570535 | 1611504 | 35.26 | 85.56 | 39826 | 10433595 | 261.98 | 114.0 | 56 | 4315 |
| *Omanocoris versicolor* | Standard | 4 | 0 | NA | NA | NA | 0 | 0 | 0.00 | 0.0 | 0 | 0 |
|  | TD-60 | 4234334 | 3141929 | 322874 | 10.28 | 163.35 | 141744 | 9772165 | 68.94 | 59.0 | 56 | 1765 |
| *Paralycambes pronotalis* | TD-60 | 4182412 | 2239022 | 110464 | 4.93 | 23.42 | 40407 | 5872762 | 145.34 | 76.0 | 56 | 15646 |
|  | TD-55 | 28442850 | 7512454 | 265534 | 3.54 | 56.76 | 58170 | 9231201 | 158.69 | 83.0 | 56 | 10143 |
| *Petascelis remipes* | Standard | 17420490 | 7823970 | 2632917 | 33.65 | 153.67 | 32110 | 9754835 | 303.79 | 187.0 | 56 | 8384 |
|  | TD-50 | 30006824 | 16134549 | 1460435 | 9.05 | 49.61 | 363683 | 141324625 | 388.59 | 284.0 | 56 | 15548 |
| *Phthia lunata* | Standard | 520 | 49 | 0 | 0.00 | NA | 2 | 113 | 56.50 | 56.5 | 56 | 57 |
|  | TD-60 | 3110532 | 1739891 | 51245 | 2.95 | 30.07 | 42531 | 3823922 | 89.91 | 61.0 | 56 | 1381 |
| *Phthiacnemia picta* | Standard | 2619550 | 1060764 | 389457 | 36.72 | 45.49 | 7642 | 2315165 | 302.95 | 157.0 | 56 | 48358 |
|  | TD-50 | 9445080 | 6162334 | 538060 | 8.73 | 33.50 | 130992 | 32867274 | 250.91 | 209.0 | 56 | 137243 |
| *Physomerus grossipes* | Standard | 4025938 | 2173257 | 524126 | 24.12 | 48.54 | 24874 | 4751208 | 191.01 | 79.0 | 56 | 4525 |
|  | TD-50 | 31049194 | 17665754 | 1025473 | 5.81 | 39.39 | 572984 | 115466074 | 201.52 | 90.0 | 56 | 15957 |
| *Plapigus abdominalis* | Standard | 6559670 | 2442042 | 562318 | 23.03 | 75.06 | 12602 | 2351639 | 186.61 | 77.0 | 56 | 2931 |
|  | TD-50 | 7322818 | 4950119 | 570181 | 11.52 | 34.69 | 116427 | 25433058 | 218.45 | 135.0 | 56 | 15902 |
| *Plectropoda* sp. | Standard | 4411586 | 2174687 | 957399 | 44.03 | 80.50 | 13034 | 4025437 | 308.84 | 215.0 | 56 | 3653 |
|  | TD-55 | 9952498 | 5617383 | 1240880 | 22.09 | 70.92 | 91206 | 24678818 | 270.58 | 221.0 | 56 | 7280 |
| *Salapia nigra* | TD-60 | 11691090 | 5352453 | 1325267 | 24.76 | 145.56 | 47011 | 7115842 | 151.37 | 76.0 | 56 | 7562 |
|  | TD-50 | 32956504 | 12293458 | 1339590 | 10.90 | 133.64 | 153173 | 31604878 | 206.33 | 215.0 | 56 | 13782 |
| *Sciophyrella neodiminuta* | TD-60 | 10247064 | 6696007 | 200360 | 2.99 | 61.63 | 248822 | 17895782 | 71.92 | 59.0 | 56 | 3350 |
|  | TD-50 | 31541592 | 14018423 | 347764 | 2.48 | 78.75 | 275903 | 27404951 | 99.33 | 66.0 | 56 | 4754 |
| *Spartocera fusca* | Standard | 8369548 | 3670141 | 1296853 | 35.34 | 88.21 | 16113 | 6799220 | 421.97 | 292.0 | 56 | 4693 |
|  | TD-60 | 3987128 | 2427427 | 994628 | 40.98 | 56.53 | 25055 | 10037032 | 400.60 | 295.0 | 56 | 4818 |
| *Sphictyrtus pretiosus* | TD-60 | 12178496 | 7759212 | 2577895 | 33.22 | 335.29 | 162399 | 14090476 | 86.76 | 61.0 | 56 | 2562 |
|  | TD-55 | 36530412 | 11479705 | 935525 | 8.15 | 128.20 | 115865 | 11926606 | 102.94 | 65.0 | 56 | 2979 |
| *Stenocoris* sp. | Standard | 5699092 | 2278471 | 785345 | 34.47 | 67.39 | 11755 | 3372784 | 286.92 | 134.0 | 56 | 3477 |
|  | TD-50 | 24721398 | 13725320 | 1767921 | 12.88 | 65.58 | 303221 | 90590648 | 298.76 | 235.0 | 56 | 15462 |

Table S4. Summary data for on-target reads and read depth of targeted loci, partitioned based on type of baits used. Abbreviations: C-baits, loci targeted by exon- or transcript-derived baits (i.e., Coreoidea-derived baits); exon, loci targeted by exon-derived baits; P-baits, loci targeted only by Pentatomomorpha-derived UCE baits; PC dual, loci targeted by both Pentatomomorpha ultraconserved element (UCE) baits and Coreoidea-derived baits (i.e., Pentatomomorpha-Coreoidea dual baits); RD, read depth; transcript, loci targeted by transcript-derived baits; see Tables S2 and S3 and Table 1 of main text for explanation of terms used.

| **Taxon** | **Target capture protocol** | **No. FRO exon** | **% FRO exon** | **RD exon** | **No. FRO transcript** | **% FRO transcript** | **RD transcript** | **No. FRO**  **C-baits** | **% FRO**  **C-baits** | **RD**  **C-baits** | **No. FRO**  **PC dual** | **% FRO**  **PC dual** | **RD**  **PC dual** | **No. FRO**  **P-baits** | **% FRO**  **P-baits** | **RD**  **P-baits** |
| --- | --- | --- | --- | --- | --- | --- | --- | --- | --- | --- | --- | --- | --- | --- | --- | --- |
| *Acanthocoris sordidus* | Standard | 596805 | 22.08 | 162.41 | 45825 | 1.70 | 162.92 | 642290 | 23.77 | 162.38 | 183373 | 6.79 | 197.59 | 373285 | 13.81 | 51.83 |
|  | TD-50 | 556521 | 3.64 | 101.14 | 74377 | 0.49 | 125.13 | 612782 | 4.01 | 101.35 | 206291 | 1.35 | 146.29 | 911101 | 5.96 | 46.79 |
| *Anasa scorbutica* | Standard | 846658 | 26.13 | 172.38 | 41903 | 1.29 | 120.04 | 887662 | 27.39 | 168.73 | 319437 | 9.86 | 240.81 | 523467 | 16.15 | 66.95 |
|  | TD-60 | 431831 | 23.32 | 93.95 | 26952 | 1.46 | 73.76 | 456377 | 24.65 | 92.12 | 150007 | 8.10 | 119.98 | 318351 | 17.19 | 29.16 |
| *Anasa varicornis* | Standard | 501323 | 18.43 | 129.25 | 28935 | 1.06 | 107.43 | 530051 | 19.49 | 127.78 | 173283 | 6.37 | 171.24 | 211721 | 7.79 | 53.73 |
|  | TD-60 | 285130 | 22.00 | 65.15 | 16848 | 1.30 | 55.38 | 301521 | 23.27 | 64.46 | 98664 | 7.61 | 90.44 | 194456 | 15.00 | 21.17 |
| *Anisoscelis gradadius* | Standard | 967935 | 23.46 | 211.33 | 52989 | 1.28 | 147.97 | 1020584 | 24.74 | 206.59 | 333140 | 8.08 | 298.44 | 740673 | 17.95 | 85.03 |
|  | TD-60 | 913413 | 23.96 | 192.41 | 62142 | 1.63 | 151.55 | 971697 | 25.49 | 188.61 | 287571 | 7.54 | 257.82 | 713631 | 18.72 | 62.59 |
| *Anoplocnemis curvipes* | Standard | 667505 | 18.06 | 77.46 | 61259 | 1.66 | 95.37 | 728430 | 19.71 | 140.11 | 237664 | 6.43 | 190.52 | 406496 | 11.00 | 50.83 |
|  | TD-50 | 481657 | 2.45 | 139.20 | 48583 | 0.28 | 151.16 | 522364 | 2.96 | 78.08 | 165184 | 0.94 | 106.89 | 835318 | 4.74 | 38.98 |
| *Catorhintha texana* | Standard | 664507 | 27.42 | 171.29 | 42761 | 1.76 | 140.92 | 702598 | 28.99 | 167.99 | 186439 | 7.69 | 196.39 | 396774 | 16.37 | 74.37 |
|  | TD-55 | 431112 | 16.85 | 86.75 | 49533 | 1.94 | 97.79 | 471795 | 18.44 | 86.69 | 143808 | 5.62 | 116.02 | 407026 | 15.91 | 32.41 |
| *Cebrenis supina* | Standard | NA | NA | NA | NA | NA | NA | NA | NA | NA | NA | NA | NA | NA | NA | NA |
|  | TD-60 | 173475 | 4.03 | 79.97 | 7924 | 0.18 | 66.50 | 181135 | 4.21 | 79.21 | 55982 | 1.30 | 93.14 | 746007 | 17.33 | 179.50 |
| *Chariesterus antennator* | Standard | 185066 | 11.20 | 63.59 | 7664 | 0.46 | 49.72 | 192661 | 11.66 | 62.83 | 51226 | 3.10 | 68.59 | 167895 | 10.16 | 44.45 |
|  | TD-60 | 217081 | 16.95 | 63.40 | 6016 | 0.47 | 45.59 | 222950 | 17.41 | 62.69 | 61696 | 4.82 | 75.87 | 161486 | 12.61 | 23.15 |
| *Chelinidea vittiger* | Standard | 454320 | 22.13 | 110.73 | 24239 | 1.18 | 94.56 | 478119 | 23.29 | 109.75 | 152604 | 7.43 | 152.44 | 241747 | 11.78 | 38.55 |
|  | TD-55 | 413666 | 11.88 | 92.66 | 26383 | 0.76 | 86.06 | 439007 | 12.61 | 92.15 | 146616 | 4.21 | 132.21 | 339163 | 9.74 | 30.95 |
| *Cletus ochraceus* | Standard | 175765 | 11.18 | 54.48 | 5743 | 0.37 | 41.78 | 181506 | 11.55 | 53.92 | 47332 | 3.01 | 62.85 | 128662 | 8.19 | 23.93 |
|  | TD-55 | 284493 | 4.03 | 78.31 | 5156 | 0.07 | 57.55 | 289476 | 4.10 | 77.76 | 66960 | 0.95 | 95.45 | 572824 | 8.12 | 41.50 |
| *Darmistus* sp. | Standard | 147541 | 3.34 | 64.67 | 13218 | 0.30 | 54.57 | 160686 | 3.64 | 63.67 | 37962 | 0.86 | 63.47 | 257730 | 5.84 | 39.09 |
|  | TD-50 | 292302 | 1.82 | 72.44 | 17566 | 0.11 | 60.36 | 308377 | 1.92 | 71.44 | 133386 | 0.83 | 127.05 | 566841 | 3.52 | 38.86 |
| Dasynini sp. | Standard | 216156 | 12.64 | 86.62 | 14978 | 0.88 | 85.89 | 229349 | 13.41 | 86.02 | 78280 | 4.58 | 110.88 | 104388 | 6.10 | 38.09 |
|  | TD-50 | 211724 | 5.42 | 60.06 | 16823 | 0.43 | 70.12 | 222137 | 5.68 | 59.289 | 72863 | 1.86 | 74.45 | 246891 | 6.31 | 24.27 |
| *Dysdercus mimus* | Standard | 96556 | 11.76 | 48.63 | 6396 | 0.78 | 29.91 | 102894 | 12.53 | 46.75 | 37426 | 4.56 | 46.84 | 152971 | 18.63 | 27.42 |
|  | TD-50 | 300005 | 3.05 | 61.79 | 36641 | 0.37 | 58.65 | 323960 | 3.29 | 60.23 | 111711 | 1.13 | 75.75 | 746371 | 7.58 | 40.01 |
| *Dysdercus suturellus* | Standard | 159693 | 8.76 | 86.87 | 13956 | 0.77 | 64.12 | 173271 | 9.50 | 84.41 | 61546 | 3.38 | 115.47 | 426150 | 23.37 | 52.54 |
|  | TD-60 | 299738 | 12.08 | 91.54 | 28599 | 1.15 | 72.78 | 327053 | 13.19 | 89.41 | 144002 | 5.81 | 130.28 | 626315 | 25.25 | 49.50 |
| *Holhymenia* sp. | Standard | 714912 | 23.32 | 153.52 | 58569 | 1.91 | 127.83 | 766893 | 25.02 | 150.58 | 195802 | 6.39 | 186.22 | 484918 | 15.82 | 54.01 |
|  | TD-55 | 826583 | 11.52 | 151.81 | 51771 | 0.72 | 124.34 | 877357 | 12.22 | 149.67 | 262885 | 3.66 | 217.67 | 803006 | 11.19 | 51.67 |
| *Hypselonotus bitrianguliger* | Standard | 408532 | 20.91 | 103.54 | 23462 | 1.20 | 89.71 | 431246 | 22.08 | 102.56 | 160968 | 8.24 | 139.25 | 254430 | 13.02 | 40.51 |
|  | TD-55 | 432017 | 9.74 | 96.24 | 32931 | 0.74 | 93.98 | 457909 | 10.33 | 95.09 | 173422 | 3.91 | 132.79 | 448225 | 10.11 | 35.11 |
| *Hypselonotus lineatus* | Standard | 348820 | 15.26 | 107.99 | 16853 | 0.74 | 85.55 | 365327 | 15.98 | 106.62 | 87910 | 3.85 | 116.31 | 168280 | 7.36 | 49.65 |
|  | TD-55 | 193748 | 9.52 | 47.89 | 13792 | 0.68 | 48.29 | 205683 | 10.11 | 47.64 | 66850 | 3.28 | 62.87 | 176899 | 8.69 | 19.09 |
| *Laminiceps festivus* | TD-60 | 11968 | 4.53 | 45.50 | 5979 | 2.26 | 75.78 | 12330 | 4.67 | 39.15 | 6255 | 2.37 | 61.13 | 8082 | 3.06 | 56.50 |
|  | TD-55 | 19268 | 3.03 | 98.94 | 7791 | 1.23 | 95.33 | 20795 | 3.27 | 81.27 | 7223 | 1.14 | 100.50 | 6202 | 0.98 | 81.66 |
| *Laminiceps obscurior* | TD-60 | 4854 | 1.40 | 17.81 | 421 | 0.12 | 12.32 | 5270 | 1.53 | 17.11 | 2400 | 0.69 | 25.29 | 744 | 0.22 | 8.48 |
|  | TD-50 | 8173 | 0.28 | 26.79 | 734 | 0.02 | 14.59 | 8792 | 0.30 | 24.22 | 4857 | 0.16 | 31.98 | 10217 | 0.34 | 48.83 |
| *Largus* sp. | Standard | 75734 | 7.47 | 41.33 | 5258 | 0.52 | 29.02 | 80961 | 7.98 | 40.23 | 31731 | 3.13 | 45.04 | 119314 | 11.76 | 24.53 |
|  | TD-55 | 185326 | 3.88 | 57.85 | 16525 | 0.35 | 58.19 | 200835 | 4.20 | 57.72 | 49604 | 1.04 | 68.98 | 410180 | 8.58 | 30.47 |
| *Leptoglossus clypealis* | Standard | 725164 | 21.65 | 161.96 | 33958 | 1.01 | 136.78 | 759030 | 22.66 | 160.58 | 223470 | 6.67 | 216.45 | 545866 | 16.30 | 65.52 |
|  | TD-60 | 464737 | 20.09 | 108.64 | 26310 | 1.14 | 90.71 | 490975 | 21.22 | 107.48 | 172086 | 7.44 | 155.93 | 425437 | 18.39 | 38.83 |
| *Leptoscelis quadrisignatus* | Standard | 855050 | 24.56 | 166.15 | 63406 | 1.82 | 147.10 | 917249 | 26.35 | 164.52 | 322355 | 9.26 | 234.13 | 536402 | 15.41 | 65.52 |
|  | TD-60 | 895443 | 22.92 | 164.58 | 80317 | 2.06 | 148.73 | 972887 | 24.90 | 162.89 | 355837 | 9.11 | 231.54 | 731924 | 18.73 | 51.15 |
| *Melanacanthus margineguttatus* | Standard | 485515 | 11.83 | 164.42 | 29350 | 0.72 | 175.30 | 513710 | 12.52 | 164.75 | 93874 | 2.29 | 166.84 | 636870 | 15.52 | 72.91 |
|  | TD-60 | 483845 | 17.68 | 116.65 | 35282 | 1.29 | 130.40 | 516712 | 18.88 | 117.11 | 178927 | 6.54 | 173.27 | 568046 | 20.76 | 48.41 |
| *Myla* sp. | Standard | 406458 | 14.07 | 118.71 | 32200 | 1.11 | 149.36 | 438380 | 15.18 | 120.49 | 107234 | 3.71 | 150.46 | 448513 | 15.53 | 51.73 |
|  | TD-55 | 743373 | 9.50 | 147.44 | 62191 | 0.79 | 163.93 | 805149 | 10.29 | 148.56 | 296473 | 3.79 | 242.09 | 933047 | 11.92 | 53.98 |
| *Nematopus lepidus* | Standard | 247164 | 4.28 | 103.32 | 16131 | 0.28 | 86.32 | 263220 | 4.56 | 102.05 | 89852 | 1.56 | 140.88 | 188804 | 3.27 | 51.77 |
|  | TD-55 | 207111 | 6.89 | 58.86 | 17077 | 0.57 | 60.90 | 223574 | 7.44 | 58.94 | 81891 | 2.72 | 85.89 | 226362 | 7.53 | 24.15 |
| *Neomegalotomus rufipes* | Standard | 523590 | 7.22 | 167.51 | 71285 | 0.98 | 208.11 | 594302 | 8.20 | 171.46 | 161155 | 2.22 | 218.69 | 868926 | 11.99 | 115.82 |
|  | TD-60 | 680895 | 14.90 | 157.05 | 59513 | 1.30 | 175.29 | 739342 | 16.18 | 158.26 | 284201 | 6.22 | 251.34 | 769173 | 16.83 | 58.28 |
| *Omanocoris versicolor* | Standard | NA | NA | NA | NA | NA | NA | NA | NA | NA | NA | NA | NA | NA | NA | NA |
|  | TD-60 | 71473 | 2.27 | 61.84 | 23274 | 0.74 | 151.70 | 58664 | 1.87 | 58.01 | 26178 | 0.83 | 85.02 | 285339 | 9.08 | 491.23 |
| *Paralycambes pronotalis* | TD-60 | 56552 | 2.53 | 35.79 | 3180 | 0.14 | 31.05 | 59669 | 2.67 | 35.46 | 15677 | 0.70 | 36.90 | 46693 | 2.09 | 17.16 |
|  | TD-55 | 130187 | 1.73 | 83.56 | 7453 | 0.10 | 71.29 | 137557 | 1.83 | 82.73 | 46788 | 0.62 | 97.46 | 101645 | 1.35 | 39.52 |
| *Petascelis remipes* | Standard | 1338819 | 17.11 | 275.21 | 127321 | 1.63 | 265.19 | 1466038 | 18.74 | 274.37 | 451248 | 5.77 | 383.15 | 970642 | 12.41 | 89.68 |
|  | TD-50 | 590524 | 3.66 | 92.17 | 82275 | 0.51 | 157.24 | 647219 | 4.01 | 93.32 | 220318 | 1.37 | 135.15 | 895067 | 5.55 | 40.67 |
| *Phthia lunata* | Standard | NA | NA | NA | NA | NA | NA | NA | NA | NA | NA | NA | NA | NA | NA | NA |
|  | TD-60 | 33748 | 1.94 | 36.01 | 18809 | 1.08 | 130.85 | 35661 | 2.05 | 34.86 | 23751 | 1.37 | 72.66 | 25818 | 1.48 | 41.19 |
| *Phthiacnemia picta* | Standard | 235876 | 22.24 | 70.22 | 9648 | 0.91 | 54.29 | 245491 | 23.14 | 69.41 | 74662 | 7.04 | 84.37 | 119018 | 11.22 | 27.86 |
|  | TD-50 | 227020 | 3.68 | 56.55 | 8444 | 0.14 | 45.59 | 235020 | 3.81 | 55.97 | 72962 | 1.18 | 74.47 | 323195 | 5.24 | 27.65 |
| *Physomerus grossipes* | Standard | 261970 | 12.05 | 63.82 | 13928 | 0.64 | 52.35 | 275832 | 12.69 | 63.08 | 104763 | 4.82 | 92.11 | 209647 | 9.65 | 37.81 |
|  | TD-50 | 400688 | 2.27 | 63.23 | 21103 | 0.12 | 47.97 | 419874 | 2.38 | 62.04 | 167472 | 0.95 | 111.49 | 663857 | 3.76 | 35.04 |
| *Plapigus abdominalis* | Standard | 312765 | 12.81 | 101.40 | 12958 | 0.53 | 92.25 | 324130 | 13.27 | 100.99 | 76505 | 3.13 | 118.19 | 201109 | 8.24 | 54.68 |
|  | TD-50 | 261091 | 5.27 | 57.02 | 14841 | 0.30 | 52.64 | 268701 | 5.43 | 56.55 | 82787 | 1.67 | 77.56 | 318060 | 6.43 | 27.85 |
| *Plectropoda* sp. | Standard | 555385 | 25.54 | 142.53 | 36136 | 1.66 | 120.15 | 591421 | 27.20 | 140.88 | 170959 | 7.86 | 188.72 | 297156 | 13.66 | 43.48 |
|  | TD-55 | 587576 | 10.46 | 138.18 | 37767 | 0.67 | 145.20 | 624366 | 11.12 | 138.50 | 229675 | 4.09 | 213.32 | 526432 | 9.37 | 42.92 |
| *Salapia nigra* | TD-60 | 662031 | 12.37 | 275.85 | 33469 | 0.63 | 257.09 | 695102 | 12.99 | 274.80 | 241407 | 4.51 | 415.87 | 502753 | 9.39 | 82.98 |
|  | TD-50 | 590777 | 4.81 | 233.62 | 30657 | 0.25 | 246.05 | 619359 | 5.04 | 233.61 | 164033 | 1.33 | 281.56 | 718013 | 5.84 | 97.93 |
| *Sciophyrella neodiminuta* | TD-60 | 122592 | 1.83 | 81.38 | 48496 | 0.72 | 320.47 | 133809 | 2.00 | 82.83 | 67621 | 1.01 | 144.81 | 79506 | 1.19 | 53.90 |
|  | TD-50 | 224290 | 1.60 | 130.16 | 165520 | 1.18 | 1134.46 | 229940 | 1.64 | 126.37 | 233589 | 1.67 | 360.68 | 223011 | 1.59 | 90.15 |
| *Spartocera fusca* | Standard | 646114 | 17.60 | 155.38 | 65562 | 1.79 | 185.27 | 711476 | 19.39 | 157.72 | 192116 | 5.23 | 195.88 | 507507 | 13.83 | 54.63 |
|  | TD-60 | 464884 | 19.15 | 104.32 | 36304 | 1.50 | 96.07 | 500961 | 20.64 | 103.66 | 181037 | 7.46 | 161.32 | 394852 | 16.27 | 33.71 |
| *Sphictyrtus pretiosus* | TD-60 | 454973 | 5.86 | 164.31 | 16140 | 0.21 | 109.83 | 470654 | 6.07 | 161.11 | 135916 | 1.75 | 197.51 | 2143446 | 27.62 | 461.75 |
|  | TD-55 | 493518 | 4.30 | 223.42 | 24081 | 0.21 | 186.42 | 516587 | 4.50 | 221.04 | 170006 | 1.48 | 287.27 | 332674 | 2.90 | 74.30 |
| *Stenocoris* sp. | Standard | 314466 | 13.80 | 106.52 | 22051 | 0.97 | 92.62 | 335413 | 14.72 | 105.34 | 93909 | 4.12 | 128.76 | 418226 | 18.36 | 52.94 |
|  | TD-50 | 737683 | 5.37 | 123.34 | 70084 | 0.51 | 129.41 | 779688 | 5.68 | 120.96 | 321558 | 2.34 | 179.86 | 1268518 | 9.24 | 63.34 |

Table S5. Summary data for captured loci targeted by exon-derived baits (376 loci targeted). Abbreviations: see Tables S2–S4.

| **Taxon** | **Target capture protocol** | **No. loci** | **% loci recovered** | **Mean locus length** | **Median locus length** | **Min locus length** | **Max locus length** | **No. putative paralogs** |
| --- | --- | --- | --- | --- | --- | --- | --- | --- |
| *Acanthocoris sordidus* | Standard | 328 | 87.23 | 1080.97 | 1035.5 | 216 | 2525 | 30 |
|  | TD-50 | 342 | 90.96 | 1399.65 | 1275.0 | 437 | 3362 | 32 |
| *Anasa scorbutica* | Standard | 337 | 89.63 | 1368.91 | 1307.0 | 307 | 3295 | 26 |
|  | TD-60 | 332 | 88.30 | 1351.42 | 1293.5 | 346 | 3044 | 37 |
| *Anasa varicornis* | Standard | 328 | 87.23 | 1118.20 | 1059.0 | 225 | 3063 | 24 |
|  | TD-60 | 336 | 89.36 | 1273.23 | 1224.5 | 214 | 2979 | 28 |
| *Anisoscelis gradadius* | Standard | 330 | 87.77 | 1312.25 | 1270.5 | 228 | 3191 | 36 |
|  | TD-60 | 335 | 89.10 | 1382.77 | 1310.0 | 213 | 3869 | 33 |
| *Anoplocnemis curvipes* | Standard | 326 | 86.70 | 1395.93 | 1375.0 | 228 | 2888 | 35) |
|  | TD-50 | 330 | 87.77 | 1576.99 | 1514.0 | 250 | 3764 | 41 |
| *Catorhintha texana* | Standard | 305 | 81.12 | 1192.76 | 1132.0 | 218 | 2829 | 26 |
|  | TD-55 | 332 | 88.30 | 1406.20 | 1363.0 | 219 | 6022 | 28 |
| *Cebrenis supina* | Standard | 0 | 0.00 | 0 | 0 | 0 | 0 | NA |
|  | TD-60 | 297 | 78.99 | 644.53 | 570.0 | 226 | 2271 | 43 |
| *Chariesterus antennator* | Standard | 301 | 80.05 | 850.40 | 782.0 | 210 | 2649 | 36 |
|  | TD-60 | 322 | 85.64 | 976.19 | 906.5 | 228 | 2827 | 40 |
| *Chelinidea vittiger* | Standard | 329 | 87.50 | 1195.12 | 1157.0 | 222 | 3029 | 33 |
|  | TD-55 | 337 | 89.63 | 1267.05 | 1215.0 | 240 | 3711 | 31 |
| *Cletus ochraceus* | Standard | 292 | 77.66 | 1002.89 | 988.0 | 206 | 2535 | 48 |
|  | TD-55 | 268 | 71.28 | 1214.97 | 1192.5 | 235 | 3105 | 96 |
| *Darmistus* sp. | Standard | 238 | 63.30 | 1168.27 | 1092.0 | 207 | 5014 | 66 |
|  | TD-50 | 315 | 83.78 | 914.16 | 907.0 | 217 | 2354 | 35 |
| Dasynini sp. | Standard | 297 | 78.99 | 799.19 | 736.0 | 215 | 2707 | 25 |
|  | TD-50 | 328 | 87.23 | 964.39 | 901.5 | 209 | 3033 | 35 |
| *Dysdercus mimus* | Standard | 184 | 48.94 | 995.17 | 983.0 | 209 | 2722 | 26 |
|  | TD-50 | 281 | 74.73 | 1330.33 | 1287.0 | 209 | 4616 | 23 |
| *Dysdercus suturellus* | Standard | 178 | 47.34 | 988.88 | 991.5 | 206 | 4171 | 72 |
|  | TD-60 | 263 | 69.95 | 1208.44 | 1183.0 | 215 | 4281 | 23 |
| *Holhymenia* sp. | Standard | 333 | 88.56 | 1349.51 | 1332.0 | 241 | 2958 | 32 |
|  | TD-55 | 343 | 91.22 | 1471.51 | 1429.0 | 297 | 3562 | 27 |
| *Hypselonotus bitrianguliger* | Standard | 333 | 88.56 | 1131.82 | 1068.0 | 215 | 2762 | 25 |
|  | TD-55 | 336 | 89.36 | 1252.20 | 1201.0 | 225 | 3088 | 33 |
| *Hypselonotus lineatus* | Standard | 303 | 80.59 | 977.94 | 938.0 | 208 | 2502 | 33 |
|  | TD-55 | 330 | 87.77 | 1167.65 | 1131.5 | 207 | 3037 | 30 |
| *Laminiceps festivus* | TD-60 | 56 | 14.89 | 278.80 | 251.0 | 206 | 681 | 18 |
|  | TD-55 | 49 | 13.03 | 258.20 | 249.0 | 207 | 399 | 6 |
| *Laminiceps obscurior* | TD-60 | 71 | 18.88 | 305.59 | 288.0 | 206 | 576 | 13 |
|  | TD-50 | 69 | 18.35 | 277.22 | 261.0 | 208 | 429 | 13 |
| *Largus* sp. | Standard | 188 | 50.00 | 932.44 | 927.5 | 206 | 2422 | 36 |
|  | TD-55 | 257 | 68.35 | 1184.34 | 1165.0 | 211 | 3132 | 51 |
| *Leptoglossus clypealis* | Standard | 335 | 89.10 | 1251.14 | 1216.0 | 227 | 3168 | 26 |
|  | TD-60 | 338 | 89.89 | 1230.58 | 1198.0 | 236 | 2971 | 33 |
| *Leptoscelis quadrisignatus* | Standard | 346 | 92.02 | 1414.01 | 1386.5 | 210 | 3174 | 18 |
|  | TD-60 | 340 | 90.43 | 1556.68 | 1509.0 | 303 | 3567 | 32 |
| *Melanacanthus margineguttatus* | Standard | 243 | 64.63 | 1139.49 | 1145.0 | 206 | 2971 | 78 |
|  | TD-60 | 313 | 83.24 | 1279.77 | 1240.0 | 253 | 3354 | 33 |
| *Myla* sp. | Standard | 272 | 72.34 | 1197.13 | 1233.5 | 212 | 2754 | 67 |
|  | TD-55 | 333 | 88.56 | 1460.68 | 1387.0 | 275 | 3387 | 32 |
| *Nematopus lepidus* | Standard | 282 | 75.00 | 785.82 | 753.5 | 206 | 2321 | 41 |
|  | TD-55 | 327 | 86.97 | 1024.85 | 993.0 | 278 | 2805 | 31 |
| *Neomegalotomus rufipes* | Standard | 255 | 67.82 | 1171.11 | 1186.0 | 237 | 3105 | 77 |
|  | TD-60 | 320 | 85.11 | 1322.08 | 1280.5 | 240 | 3570 | 34 |
| *Omanocoris versicolor* | Standard | NA | NA | NA | NA | NA | NA | NA |
|  | TD-60 | 200 | 53.19 | 337.44 | 286.0 | 206 | 1192 | 43 |
| *Paralycambes pronotalis* | TD-60 | 266 | 70.74 | 502.48 | 456.5 | 209 | 1675 | 40 |
|  | TD-55 | 274 | 72.87 | 511.35 | 446.0 | 206 | 1805 | 30 |
| *Petascelis remipes* | Standard | 322 | 85.64 | 1001.49 | 932.0 | 213 | 2865 | 44 |
|  | TD-50 | 341 | 90.69 | 1056.11 | 1008.5 | 226 | 4195 | 31 |
| *Phthia lunata* | Standard | 0 | 0 | 0 | 0 | 0 | 0 | NA |
|  | TD-60 | 192 | 51.06 | 346.04 | 291.0 | 206 | 1148 | 42 |
| *Phthiacnemia picta* | Standard | 315 | 83.78 | 594.24 | 537.0 | 206 | 2705 | 25 |
|  | TD-50 | 328 | 87.23 | 674.45 | 646.0 | 206 | 2846 | 40 |
| *Physomerus grossipes* | Standard | 325 | 86.44 | 1152.35 | 1160.0 | 213 | 2687 | 24 |
|  | TD-50 | 341 | 90.69 | 1498.37 | 1453.0 | 225 | 3402 | 31 |
| *Plapigus abdominalis* | Standard | 285 | 75.80 | 1016.26 | 998.0 | 207 | 2443 | 47 |
|  | TD-50 | 328 | 87.23 | 1211.08 | 1162.0 | 242 | 3726 | 34 |
| *Plectropoda* sp. | Standard | 328 | 87.23 | 1123.21 | 1091.0 | 207 | 2591 | 35 |
|  | TD-55 | 336 | 89.36 | 1209.00 | 1157.5 | 207 | 3007 | 34 |
| *Salapia nigra* | TD-60 | 334 | 88.83 | 655.88 | 583.0 | 232 | 2156 | 35 |
|  | TD-50 | 335 | 89.10 | 651.41 | 570.0 | 220 | 2193 | 33 |
| *Sciophyrella neodiminuta* | TD-60 | 288 | 76.60 | 398.99 | 322.0 | 195 | 1978 | 37 |
|  | TD-50 | 287 | 76.33 | 403.47 | 323.0 | 172 | 1893 | 37 |
| *Spartocera fusca* | Standard | 313 | 83.24 | 1287.99 | 1273.0 | 263 | 2884 | 47 |
|  | TD-60 | 338 | 89.89 | 1301.13 | 1256.5 | 215 | 2924 | 30 |
| *Sphictyrtus pretiosus* | TD-60 | 332 | 88.30 | 685.35 | 601.5 | 217 | 2395 | 34 |
|  | TD-55 | 326 | 86.70 | 625.66 | 546.5 | 208 | 2310 | 31 |
| *Stenocoris* sp. | Standard | 265 | 70.48 | 1051.71 | 1060.0 | 231 | 2968 | 61 |
|  | TD-50 | 335 | 89.10 | 1439.48 | 1358.0 | 332 | 4394 | 34 |

Table S6. Summary data for captured loci targeted by transcript-derived baits (58 loci targeted). Abbreviations: see Tables S2–S4.

| **Taxon** | **Target capture protocol** | **No. loci** | **% loci recovered** | **Mean locus length** | **Median locus length** | **Min locus length** | **Max locus length** | **No. putative paralogs** |
| --- | --- | --- | --- | --- | --- | --- | --- | --- |
| *Acanthocoris sordidus* | Standard | 22 | 37.93 | 1227.91 | 1278.0 | 445 | 1941 | 36 |
|  | TD-50 | 25 | 43.10 | 1922.24 | 1624.0 | 889 | 4075 | 33 |
| *Anasa scorbutica* | Standard | 19 | 32.76 | 1747.63 | 1557.0 | 1207 | 3869 | 39 |
|  | TD-60 | 19 | 32.76 | 1841.00 | 1682.0 | 1196 | 3092 | 39 |
| *Anasa varicornis* | Standard | 21 | 36.21 | 1220.81 | 1190.0 | 454 | 2150 | 37 |
|  | TD-60 | 19 | 32.76 | 1563.16 | 1446.0 | 976 | 2284 | 39 |
| *Anisoscelis gradadius* | Standard | 21 | 36.21 | 1659.90 | 1577.0 | 779 | 2697 | 37 |
|  | TD-60 | 22 | 37.93 | 1813.82 | 1696.0 | 1140 | 3243 | 36 |
| *Anoplocnemis curvipes* | Standard | 20 | 34.48 | 1939.75 | 1730.0 | 1103 | 3650 | 37 |
|  | TD-50 | 19 | 32.76 | 2286.53 | 1858.0 | 1049 | 5170 | 39 |
| *Catorhintha texana* | Standard | 19 | 32.76 | 1503.05 | 1468.0 | 844 | 2558 | 37 |
|  | TD-55 | 24 | 41.38 | 1879.21 | 1755.0 | 595 | 3473 | 34 |
| *Cebrenis supina* | Standard | 0 | 0 | 0 | 0 | 0 | 0 | NA |
|  | TD-60 | 16 | 27.59 | 670.13 | 674.0 | 232 | 996 | 42 |
| *Chariesterus antennator* | Standard | 15 | 25.86 | 978.67 | 992.0 | 270 | 1411 | 42 |
|  | TD-60 | 12 | 20.69 | 1060.67 | 1053.5 | 767 | 1320 | 46 |
| *Chelinidea vittiger* | Standard | 18 | 31.03 | 1360.89 | 1396.0 | 831 | 2341 | 40 |
|  | TD-55 | 20 | 34.48 | 1468.80 | 1351.5 | 672 | 2569 | 38 |
| *Cletus ochraceus* | Standard | 13 | 22.41 | 1025.08 | 1095.0 | 444 | 1472 | 44 |
|  | TD-55 | 7 | 12.07 | 1214.00 | 1205.0 | 825 | 1694 | 51 |
| *Darmistus* sp. | Standard | 20 | 34.48 | 1169.35 | 1020.0 | 433 | 2770 | 33 |
|  | TD-50 | 18 | 31.03 | 1457.83 | 1274.0 | 640 | 2867 | 39 |
| Dasynini sp. | Standard | 20 | 34.48 | 726.65 | 634.5 | 430 | 1156 | 37 |
|  | TD-50 | 16 | 27.59 | 1313.63 | 1050.0 | 626 | 5003 | 42 |
| *Dysdercus mimus* | Standard | 26 | 44.83 | 783.77 | 727.0 | 230 | 1480 | 14 |
|  | TD-50 | 29 | 50.00 | 1474.38 | 1411.0 | 640 | 3467 | 21 |
| *Dysdercus suturellus* | Standard | 23 | 39.66 | 894.39 | 995.0 | 222 | 1931 | 22 |
|  | TD-60 | 28 | 48.28 | 1341.57 | 1233.5 | 667 | 3496 | 22 |
| *Holhymenia* sp. | Standard | 20 | 34.48 | 1946.60 | 1671.0 | 1149 | 3799 | 38 |
|  | TD-55 | 20 | 34.48 | 2009.10 | 1781.0 | 1083 | 4105 | 38 |
| *Hypselonotus bitrianguliger* | Standard | 19 | 32.76 | 1303.37 | 1368.0 | 482 | 2018 | 39 |
|  | TD-55 | 19 | 32.76 | 1612.05 | 1579.0 | 946 | 2980 | 39 |
| *Hypselonotus lineatus* | Standard | 16 | 27.59 | 1141.44 | 1148.5 | 473 | 2173 | 39 |
|  | TD-55 | 18 | 31.03 | 1447.94 | 1403.0 | 328 | 3301 | 40 |
| *Laminiceps festivus* | TD-60 | 15 | 25.86 | 239.93 | 231.0 | 207 | 323 | 5 |
|  | TD-55 | 19 | 32.76 | 239.79 | 230.0 | 207 | 374 | 2 |
| *Laminiceps obscurior* | TD-60 | 13 | 22.41 | 240.31 | 242.0 | 206 | 280 | 7 |
|  | TD-50 | 18 | 31.03 | 259.28 | 233.0 | 213 | 535 | 2 |
| *Largus* sp. | Standard | 21 | 36.21 | 817.67 | 847.0 | 302 | 1484 | 23 |
|  | TD-55 | 22 | 37.93 | 1155.91 | 1200.5 | 285 | 1770 | 33 |
| *Leptoglossus clypealis* | Standard | 16 | 27.59 | 1502.25 | 1447.0 | 806 | 2759 | 42 |
|  | TD-60 | 17 | 29.31 | 1683.71 | 1546.0 | 993 | 3315 | 41 |
| *Leptoscelis quadrisignatus* | Standard | 22 | 37.93 | 1876.27 | 1760.0 | 1310 | 2820 | 36 |
|  | TD-60 | 24 | 41.38 | 2172.08 | 1901.0 | 1425 | 4609 | 34 |
| *Melanacanthus margineguttatus* | Standard | 11 | 18.97 | 1433.18 | 1319.0 | 726 | 2535 | 47 |
|  | TD-60 | 17 | 29.31 | 1518.24 | 1555.0 | 759 | 2412 | 41 |
| *Myla* sp. | Standard | 15 | 25.86 | 1389.07 | 1386.0 | 994 | 1911 | 43 |
|  | TD-55 | 19 | 32.76 | 1955.00 | 1756.0 | 692 | 3784 | 39 |
| *Nematopus lepidus* | Standard | 19 | 32.76 | 931.32 | 909.0 | 297 | 1428 | 38 |
|  | TD-55 | 20 | 34.48 | 1294.25 | 1181.5 | 737 | 2512 | 38 |
| *Neomegalotomus rufipes* | Standard | 17 | 29.31 | 1941.76 | 1690.0 | 817 | 3572 | 41 |
|  | TD-60 | 18 | 31.03 | 1842.33 | 1705.0 | 975 | 3369 | 40 |
| *Omanocoris versicolor* | Standard | NA | NA | NA | NA | NA | NA | NA |
|  | TD-60 | 22 | 37.93 | 305.64 | 243.0 | 209 | 592 | 30 |
| *Paralycambes pronotalis* | TD-60 | 20 | 34.48 | 478.45 | 436.0 | 251 | 863 | 36 |
|  | TD-55 | 20 | 34.48 | 496.75 | 440.0 | 262 | 879 | 35 |
| *Petascelis remipes* | Standard | 25 | 43.10 | 1847.60 | 1680.0 | 1324 | 3355 | 33 |
|  | TD-50 | 25 | 43.10 | 2167.92 | 1931.0 | 1303 | 3820 | 33 |
| *Phthia lunata* | Standard | 0 | 0 | 0 | 0 | 0 | 0 | NA |
|  | TD-60 | 23 | 39.66 | 310.70 | 267.0 | 217 | 866 | 25 |
| *Phthiacnemia picta* | Standard | 16 | 27.59 | 1067.44 | 1030.5 | 642 | 1658 | 41 |
|  | TD-50 | 14 | 24.14 | 1260.07 | 1281.0 | 794 | 1590 | 44 |
| *Physomerus grossipes* | Standard | 21 | 36.21 | 1217.10 | 1237.0 | 328 | 1903 | 37 |
|  | TD-50 | 21 | 36.21 | 1810.10 | 1706.0 | 1138 | 2802 | 37 |
| *Plapigus abdominalis* | Standard | 13 | 22.41 | 1025.00 | 1069.0 | 533 | 1383 | 43 |
|  | TD-50 | 17 | 29.31 | 1381.29 | 1448.0 | 328 | 2817 | 41 |
| *Plectropoda* sp. | Standard | 19 | 32.76 | 1535.26 | 1432.0 | 824 | 3124 | 39 |
|  | TD-55 | 17 | 29.31 | 1471.76 | 1488.0 | 837 | 2262 | 41 |
| *Salapia nigra* | TD-60 | 17 | 29.31 | 714.53 | 782.0 | 462 | 975 | 41 |
|  | TD-50 | 17 | 29.31 | 667.12 | 585.0 | 375 | 1013 | 41 |
| *Sciophyrella neodiminuta* | TD-60 | 20 | 34.48 | 422.75 | 333.5 | 219 | 918 | 36 |
|  | TD-50 | 17 | 29.31 | 433.65 | 401.0 | 215 | 751 | 40 |
| *Spartocera fusca* | Standard | 21 | 36.21 | 1639.10 | 1537.0 | 908 | 3017 | 36 |
|  | TD-60 | 22 | 37.93 | 1692.23 | 1538.5 | 1024 | 3133 | 36 |
| *Sphictyrtus pretiosus* | TD-60 | 18 | 31.03 | 759.94 | 797.5 | 424 | 1131 | 40 |
|  | TD-55 | 17 | 29.31 | 706.29 | 755.0 | 278 | 976 | 41 |
| *Stenocoris* sp. | Standard | 17 | 29.31 | 1284.47 | 1290.0 | 714 | 2051 | 38 |
|  | TD-50 | 23 | 39.66 | 1769.83 | 1848.0 | 522 | 4899 | 35 |

Table S7. Summary data for captured loci targeted by Pentatomomorpha-Coreoidea dual baits (103 loci targeted). Abbreviations: see Tables S2–S4.

| **Taxon** | **Target capture protocol** | **No. loci** | **% loci recovered** | **Mean locus length** | **Median locus length** | **Min locus length** | **Max locus length** | **No. putative paralogs** |
| --- | --- | --- | --- | --- | --- | --- | --- | --- |
| *Acanthocoris sordidus* | Standard | 70 | 67.96 | 1279.23 | 1092.5 | 514 | 3047 | 33 |
|  | TD-50 | 71 | 68.93 | 1686.20 | 1515.0 | 783 | 3637 | 32 |
| *Anasa scorbutica* | Standard | 74 | 71.84 | 1705.66 | 1519.0 | 642 | 4246 | 29 |
|  | TD-60 | 72 | 69.90 | 1676.19 | 1407.5 | 721 | 4095 | 31 |
| *Anasa varicornis* | Standard | 68 | 66.02 | 1397.97 | 1245.0 | 540 | 2970 | 36 |
|  | TD-60 | 68 | 66.02 | 1569.09 | 1369.0 | 609 | 3109 | 36 |
| *Anisoscelis gradadius* | Standard | 67 | 65.05 | 1542.18 | 1392.0 | 592 | 3118 | 37 |
|  | TD-60 | 65 | 63.11 | 1671.15 | 1442.0 | 515 | 3924 | 38 |
| *Anoplocnemis curvipes* | Standard | 70 | 67.96 | 1702.67 | 1595.0 | 480 | 4791 | 33 |
|  | TD-50 | 66 | 64.08 | 1957.94 | 1722.0 | 674 | 4998 | 38 |
| *Catorhintha texana* | Standard | 67 | 65.05 | 1322.37 | 1238.0 | 317 | 3010 | 34 |
|  | TD-55 | 69 | 66.99 | 1674.59 | 1484.0 | 274 | 3358 | 32 |
| *Cebrenis supina* | Standard | 0 | 0 | 0 | 0 | 0 | 0 | NA |
|  | TD-60 | 67 | 65.05 | 833.76 | 661.0 | 287 | 2341 | 34 |
| *Chariesterus antennator* | Standard | 69 | 66.99 | 1023.20 | 843.0 | 295 | 2757 | 35 |
|  | TD-60 | 64 | 62.14 | 1198.03 | 1028.0 | 430 | 2750 | 40 |
| *Chelinidea vittiger* | Standard | 66 | 64.08 | 1472.83 | 1335.5 | 340 | 2954 | 36 |
|  | TD-55 | 68 | 66.02 | 1585.94 | 1383.0 | 474 | 3245 | 35 |
| *Cletus ochraceus* | Standard | 63 | 61.17 | 1130.16 | 1018.0 | 208 | 2865 | 40 |
|  | TD-55 | 46 | 44.66 | 1357.93 | 1312.0 | 476 | 3243 | 59 |
| *Darmistus* sp. | Standard | 55 | 53.40 | 1047.42 | 1012.0 | 242 | 2784 | 42 |
|  | TD-50 | 66 | 64.08 | 1459.94 | 1271.0 | 580 | 3163 | 35 |
| Dasynini sp. | Standard | 70 | 67.96 | 957.80 | 843.5 | 210 | 2484 | 28 |
|  | TD-50 | 70 | 67.96 | 1314.10 | 1105.5 | 418 | 5445 | 32 |
| *Dysdercus mimus* | Standard | 67 | 65.05 | 1124.25 | 985.0 | 216 | 2593 | 21 |
|  | TD-50 | 67 | 65.05 | 1650.19 | 1441.0 | 452 | 4814 | 36 |
| *Dysdercus suturellus* | Standard | 46 | 44.66 | 1039.26 | 1062.0 | 265 | 2018 | 46 |
|  | TD-60 | 67 | 65.05 | 1583.67 | 1356.0 | 215 | 3258 | 32 |
| *Holhymenia* sp. | Standard | 62 | 60.19 | 1621.19 | 1403.5 | 721 | 3428 | 41 |
|  | TD-55 | 63 | 61.17 | 1796.17 | 1546.0 | 699 | 4905 | 41 |
| *Hypselonotus bitrianguliger* | Standard | 79 | 76.70 | 1401.65 | 1277.0 | 254 | 3005 | 24 |
|  | TD-55 | 76 | 73.79 | 1613.59 | 1378.5 | 459 | 3326 | 27 |
| *Hypselonotus lineatus* | Standard | 64 | 62.14 | 1113.45 | 1034.5 | 245 | 2860 | 36 |
|  | TD-55 | 70 | 67.96 | 1432.89 | 1193.0 | 498 | 4133 | 33 |
| *Laminiceps festivus* | TD-60 | 20 | 19.42 | 256.05 | 228.0 | 209 | 506 | 13 |
|  | TD-55 | 16 | 15.53 | 254.88 | 240.0 | 210 | 346 | 12 |
| *Laminiceps obscurior* | TD-60 | 19 | 18.45 | 292.32 | 283.0 | 208 | 388 | 10 |
|  | TD-50 | 26 | 25.24 | 277.19 | 256.0 | 208 | 636 | 10 |
| *Largus* sp. | Standard | 60 | 58.25 | 1121.45 | 1101.0 | 293 | 2756 | 27 |
|  | TD-55 | 48 | 46.60 | 1403.23 | 1344.0 | 254 | 3262 | 53 |
| *Leptoglossus clypealis* | Standard | 65 | 63.11 | 1534.09 | 1367.0 | 343 | 3086 | 38 |
|  | TD-60 | 68 | 66.02 | 1586.71 | 1348.5 | 583 | 3313 | 35 |
| *Leptoscelis quadrisignatus* | Standard | 76 | 73.79 | 1743.18 | 1630.5 | 606 | 3210 | 27 |
|  | TD-60 | 78 | 75.73 | 1926.12 | 1731.0 | 757 | 3494 | 25 |
| *Melanacanthus margineguttatus* | Standard | 41 | 39.81 | 1316.93 | 1230.0 | 263 | 2368 | 57 |
|  | TD-60 | 62 | 60.19 | 1618.16 | 1481.0 | 406 | 3519 | 38 |
| *Myla* sp. | Standard | 53 | 51.46 | 1288.40 | 1268.0 | 283 | 3295 | 48 |
|  | TD-55 | 66 | 64.08 | 1781.24 | 1592.5 | 315 | 3704 | 37 |
| *Nematopus lepidus* | Standard | 62 | 60.19 | 971.02 | 810.0 | 236 | 2552 | 35 |
|  | TD-55 | 70 | 67.96 | 1287.80 | 1097.5 | 351 | 2900 | 32 |
| *Neomegalotomus rufipes* | Standard | 56 | 54.37 | 1244.18 | 1142.5 | 315 | 3329 | 45 |
|  | TD-60 | 67 | 65.05 | 1642.37 | 1452.0 | 472 | 4315 | 36 |
| *Omanocoris versicolor* | Standard | NA | NA | NA | NA | NA | NA | NA |
|  | TD-60 | 46 | 44.66 | 395.13 | 295.0 | 214 | 1111 | 38 |
| *Paralycambes pronotalis* | TD-60 | 62 | 60.19 | 615.69 | 476.0 | 223 | 2251 | 35 |
|  | TD-55 | 65 | 63.11 | 704.92 | 479.0 | 214 | 2317 | 30 |
| *Petascelis remipes* | Standard | 66 | 64.08 | 1715.06 | 1613.5 | 695 | 3485 | 40 |
|  | TD-50 | 72 | 69.90 | 1918.25 | 1787.5 | 484 | 3883 | 34 |
| *Phthia lunata* | Standard | 0 | 0 | 0 | 0 | 0 | 0 | NA |
|  | TD-60 | 49 | 47.57 | 373.18 | 288.0 | 214 | 1356 | 34 |
| *Phthiacnemia picta* | Standard | 69 | 66.99 | 1236.75 | 1084.0 | 396 | 2705 | 34 |
|  | TD-50 | 64 | 62.14 | 1407.44 | 1153.0 | 413 | 2821 | 40 |
| *Physomerus grossipes* | Standard | 73 | 70.87 | 1448.41 | 1390.0 | 396 | 2942 | 28 |
|  | TD-50 | 68 | 66.02 | 1853.78 | 1715.0 | 729 | 4819 | 35 |
| *Plapigus abdominalis* | Standard | 57 | 55.34 | 1024.68 | 981.0 | 219 | 2287 | 43 |
|  | TD-50 | 64 | 62.14 | 1459.97 | 1217.5 | 356 | 3091 | 37 |
| *Plectropoda* sp. | Standard | 66 | 64.08 | 1335.23 | 1152.0 | 430 | 3184 | 38 |
|  | TD-55 | 68 | 66.02 | 1532.78 | 1330.5 | 578 | 3186 | 36 |
| *Salapia nigra* | TD-60 | 62 | 60.19 | 800.95 | 621.0 | 249 | 2441 | 41 |
|  | TD-50 | 63 | 61.17 | 860.83 | 632.0 | 262 | 2538 | 40 |
| *Sciophyrella neodiminuta* | TD-60 | 59 | 57.28 | 492.83 | 340.0 | 206 | 2031 | 40 |
|  | TD-50 | 62 | 60.19 | 589.16 | 362.5 | 209 | 2306 | 38 |
| *Spartocera fusca* | Standard | 58 | 56.31 | 1642.93 | 1433.0 | 806 | 3322 | 41 |
|  | TD-60 | 64 | 62.14 | 1734.19 | 1580.5 | 633 | 3309 | 35 |
| *Sphictyrtus pretiosus* | TD-60 | 68 | 66.02 | 913.44 | 703.5 | 342 | 2562 | 35 |
|  | TD-55 | 68 | 66.02 | 820.79 | 623.0 | 271 | 2382 | 35 |
| *Stenocoris* sp. | Standard | 58 | 56.31 | 1169.38 | 1115.0 | 229 | 2739 | 45 |
|  | TD-50 | 70 | 67.96 | 1947.11 | 1851.5 | 544 | 5042 | 33 |

Table S8. Summary data for captured loci targeted by Pentatomomorpha-derived baits (2566 loci targeted). Abbreviations: see Tables S2–S4.

| **Taxon** | **Target capture protocol** | **No. loci** | **% loci recovered** | **Mean locus length** | **Median locus length** | **Min locus length** | **Max locus length** | **No. putative paralogs** |
| --- | --- | --- | --- | --- | --- | --- | --- | --- |
| *Acanthocoris sordidus* | Standard | 1011 | 39.40 | 681.53 | 672.0 | 101 | 1999 | 5 |
|  | TD-50 | 1674 | 65.24 | 978.59 | 953.5 | 206 | 4247 | 21 |
| *Anasa scorbutica* | Standard | 918 | 35.78 | 778.41 | 755.5 | 206 | 3869 | 9 |
|  | TD-60 | 1327 | 51.71 | 788.06 | 767.0 | 207 | 3158 | 14 |
| *Anasa varicornis* | Standard | 612 | 23.85 | 605.50 | 552.0 | 206 | 2771 | 5 |
|  | TD-60 | 1189 | 46.34 | 736.95 | 710.0 | 208 | 3290 | 14 |
| *Anisoscelis gradadius* | Standard | 1005 | 39.17 | 766.69 | 747.0 | 161 | 3191 | 16 |
|  | TD-60 | 1376 | 53.62 | 826.35 | 808.5 | 161 | 3924 | 22 |
| *Anoplocnemis curvipes* | Standard | 902 | 35.15 | 835.63 | 811.0 | 207 | 5356 | 12 |
|  | TD-50 | 1619 | 63.09 | 1055.50 | 1007.0 | 208 | 5762 | 37 |
| *Catorhintha texana* | Standard | 705 | 27.47 | 694.34 | 646.0 | 206 | 2651 | 6 |
|  | TD-55 | 1341 | 52.26 | 850.93 | 837.0 | 206 | 2910 | 18 |
| *Cebrenis supina* | Standard | 0 | 0 | 0 | 0 | 0 | 0 | NA |
|  | TD-60 | 921 | 35.89 | 372.80 | 338.0 | 165 | 2100 | 11 |
| *Chariesterus antennator* | Standard | 626 | 24.40 | 479.03 | 421.0 | 207 | 2649 | 14 |
|  | TD-60 | 1127 | 43.92 | 546.04 | 499.0 | 200 | 2688 | 39 |
| *Chelinidea vittiger* | Standard | 866 | 33.75 | 670.86 | 613.5 | 206 | 2960 | 6 |
|  | TD-55 | 1443 | 56.24 | 701.38 | 662.0 | 206 | 4043 | 17 |
| *Cletus ochraceus* | Standard | 740 | 28.84 | 622.62 | 587.0 | 208 | 2380 | 13 |
|  | TD-55 | 1460 | 56.90 | 803.21 | 783.5 | 206 | 3269 | 66 |
| *Darmistus* sp. | Standard | 890 | 34.68 | 659.32 | 626.5 | 206 | 2784 | 10 |
|  | TD-50 | 1554 | 60.56 | 787.53 | 776.0 | 142 | 3699 | 29 |
| Dasynini sp. | Standard | 535 | 20.85 | 481.28 | 431.0 | 206 | 1949 | 5 |
|  | TD-50 | 1377 | 53.66 | 633.24 | 603.0 | 206 | 2957 | 23 |
| *Dysdercus mimus* | Standard | 737 | 28.72 | 710.91 | 669.0 | 207 | 2439 | 10 |
|  | TD-50 | 1484 | 57.83 | 976.56 | 968.5 | 209 | 4238 | 43 |
| *Dysdercus suturellus* | Standard | 924 | 36.01 | 836.40 | 821.5 | 208 | 2623 | 22 |
|  | TD-60 | 1325 | 51.64 | 921.98 | 920.0 | 208 | 3258 | 33 |
| *Holhymenia* sp. | Standard | 971 | 37.84 | 817.19 | 808.0 | 206 | 3214 | 10 |
|  | TD-55 | 1521 | 59.28 | 937.19 | 925.0 | 120 | 3551 | 19 |
| *Hypselonotus bitrianguliger* | Standard | 867 | 33.79 | 681.34 | 660.0 | 206 | 2606 | 7 |
|  | TD-55 | 1464 | 57.05 | 793.53 | 778.5 | 206 | 3326 | 19 |
| *Hypselonotus lineatus* | Standard | 559 | 21.78 | 570.09 | 508.0 | 206 | 2538 | 8 |
|  | TD-55 | 1203 | 46.88 | 699.37 | 659.0 | 206 | 3037 | 18 |
| *Laminiceps festivus* | TD-60 | 29 | 1.13 | 252.93 | 236.0 | 70 | 672 | 1 |
|  | TD-55 | 17 | 0.66 | 240.53 | 231.0 | 209 | 286 | 0 |
| *Laminiceps obscurior* | TD-60 | 31 | 1.21 | 255.87 | 251.0 | 206 | 399 | 0 |
|  | TD-50 | 40 | 1.56 | 260.75 | 254.0 | 206 | 369 | 0 |
| *Largus* sp. | Standard | 687 | 26.77 | 670.06 | 643.0 | 206 | 2756 | 9 |
|  | TD-55 | 1455 | 56.70 | 858.17 | 848.0 | 206 | 3262 | 43 |
| *Leptoglossus clypealis* | Standard | 990 | 38.58 | 736.55 | 706.0 | 206 | 3709 | 9 |
|  | TD-60 | 1358 | 52.92 | 755.35 | 724.5 | 157 | 4825 | 18 |
| *Leptoscelis quadrisignatus* | Standard | 925 | 36.05 | 815.21 | 778.0 | 207 | 3878 | 10 |
|  | TD-60 | 1454 | 56.66 | 938.90 | 932.5 | 207 | 3977 | 9 |
| *Melanacanthus margineguttatus* | Standard | 1008 | 39.28 | 806.99 | 774.0 | 60 | 3371 | 52 |
|  | TD-60 | 1357 | 52.88 | 823.37 | 789.0 | 165 | 3519 | 73 |
| *Myla* sp. | Standard | 1019 | 39.71 | 810.78 | 800.0 | 131 | 3295 | 17 |
|  | TD-55 | 1632 | 63.60 | 994.15 | 991.5 | 215 | 3704 | 39 |
| *Nematopus lepidus* | Standard | 665 | 25.92 | 516.34 | 474.0 | 201 | 2326 | 4 |
|  | TD-55 | 1353 | 52.73 | 644.00 | 623.0 | 206 | 2900 | 8 |
| *Neomegalotomus rufipes* | Standard | 1063 | 41.43 | 820.08 | 780.0 | 206 | 3950 | 30 |
|  | TD-60 | 1450 | 56.51 | 877.91 | 871.0 | 206 | 3984 | 40 |
| *Omanocoris versicolor* | Standard | NA | NA | NA | NA | NA | NA | NA |
|  | TD-60 | 145 | 5.65 | 253.11 | 229.0 | 206 | 953 | 2 |
| *Paralycambes pronotalis* | TD-60 | 713 | 27.79 | 334.31 | 304.0 | 206 | 1539 | 9 |
|  | TD-55 | 709 | 27.63 | 334.17 | 312.0 | 206 | 1494 | 6 |
| *Petascelis remipes* | Standard | 1124 | 43.80 | 893.43 | 883.0 | 206 | 3485 | 6 |
|  | TD-50 | 1652 | 64.38 | 1114.55 | 1095.0 | 209 | 3596 | 17 |
| *Phthia lunata* | Standard | 0 | 0 | 0 | 0 | 0 | 0 | NA |
|  | TD-60 | 134 | 5.22 | 277.51 | 230.5 | 206 | 1356 | 1 |
| *Phthiacnemia picta* | Standard | 691 | 26.93 | 567.55 | 519.0 | 206 | 2829 | 5 |
|  | TD-50 | 1399 | 54.52 | 664.68 | 638.0 | 206 | 4195 | 23 |
| *Physomerus grossipes* | Standard | 686 | 26.73 | 702.04 | 655.5 | 206 | 2524 | 4 |
|  | TD-50 | 1542 | 60.09 | 963.65 | 967.0 | 206 | 4236 | 24 |
| *Plapigus abdominalis* | Standard | 578 | 22.53 | 582.03 | 533.0 | 206 | 2754 | 13 |
|  | TD-50 | 1354 | 52.77 | 698.99 | 664.0 | 207 | 4044 | 16 |
| *Plectropoda* sp. | Standard | 967 | 37.69 | 672.37 | 648.0 | 206 | 3653 | 9 |
|  | TD-55 | 1517 | 59.12 | 747.63 | 729.0 | 73 | 4253 | 26 |
| *Salapia nigra* | TD-60 | 1393 | 54.29 | 387.07 | 374.0 | 206 | 2079 | 9 |
|  | TD-50 | 1519 | 59.20 | 388.01 | 366.0 | 157 | 2908 | 13 |
| *Sciophyrella neodiminuta* | TD-60 | 396 | 15.43 | 262.00 | 235.5 | 206 | 1660 | 4 |
|  | TD-50 | 607 | 23.66 | 266.69 | 241.0 | 202 | 1893 | 5 |
| *Spartocera fusca* | Standard | 1087 | 42.36 | 815.85 | 803.0 | 210 | 3322 | 17 |
|  | TD-60 | 1395 | 54.36 | 820.77 | 814.0 | 206 | 3309 | 23 |
| *Sphictyrtus pretiosus* | TD-60 | 1213 | 47.27 | 401.12 | 378.0 | 206 | 2391 | 12 |
|  | TD-55 | 1087 | 42.36 | 373.40 | 344.0 | 206 | 2286 | 12 |
| *Stenocoris* sp. | Standard | 1017 | 39.63 | 704.56 | 665.0 | 206 | 3062 | 13 |
|  | TD-50 | 1630 | 63.52 | 996.92 | 965.5 | 213 | 5558 | 28 |

Table S9. Summary data for raw and filtered sequence reads, reads on-target and read depth across all targeted loci, and contigs from 24 taxa that had 2,000,000 million raw reads subsampled to equalize sequencing depth across capture conditions. Abbreviations: see Tables S2 and S3.

| **Taxon** | **Target capture protocol** | **No. filtered reads** | **No. total FRO** | **% Total FRO** | **Overall read depth** | **No. contigs** | **Total bp across contigs** | **Mean contig length** | **Median contig length** | **Min contig length** | **Max contig length** |
| --- | --- | --- | --- | --- | --- | --- | --- | --- | --- | --- | --- |
| *Acanthocoris sordidus* | Standard | 1075872 | 573549 | 53.31 | 52.58 | 6292 | 3009400 | 478.29 | 385.5 | 56 | 2693 |
|  | TD-50 | 1209143 | 237160 | 19.61 | 20.02 | 34324 | 11528337 | 335.87 | 272 | 56 | 13992 |
| *Anasa scorbutica* | Standard | 1048842 | 572806 | 54.61 | 47.48 | 7693 | 2986228 | 388.17 | 247 | 56 | 4047 |
|  | TD-60 | 1171167 | 570430 | 48.71 | 36.70 | 13204 | 4800827 | 363.59 | 241 | 56 | 5640 |
| *Anisoscelis gradadius* | Standard | 1474581 | 733420 | 49.74 | 62.15 | 12140 | 4032746 | 332.19 | 236 | 56 | 2994 |
|  | TD-60 | 1531468 | 748818 | 48.90 | 51.44 | 15949 | 5649107 | 354.2 | 258 | 56 | 4776 |
| *Anoplocnemis curvipes* | Standard | 1179798 | 547895 | 46.44 | 45.91 | 9693 | 3244754 | 334.75 | 189 | 56 | 5332 |
|  | TD-50 | 1128259 | 142357 | 12.62 | 13.90 | 38913 | 8344468 | 214.44 | 123 | 56 | 8048 |
| *Catorhintha texana* | Standard | 806301 | 451325 | 55.98 | 44.90 | 4408 | 1827259 | 414.53 | 248 | 56 | 3421 |
|  | TD-55 | 1212405 | 495582 | 40.88 | 31.14 | 16520 | 6363459 | 385.2 | 269 | 56 | 6607 |
| *Chelinidea vittiger* | Standard | 1107317 | 497889 | 44.96 | 48.41 | 8639 | 2798634 | 323.95 | 224 | 56 | 2960 |
|  | TD-55 | 1317593 | 357913 | 27.16 | 29.12 | 28921 | 7252679 | 250.78 | 211 | 56 | 3956 |
| *Darmistus* sp. | Standard | 1531115 | 213636 | 13.95 | 24.64 | 19474 | 3255993 | 167.2 | 75 | 56 | 4487 |
|  | TD-50 | 1489119 | 85005 | 5.71 | 10.73 | 49158 | 8866622 | 180.37 | 105 | 56 | 7819 |
| Dasynini sp. | Standard | 864548 | 223026 | 25.80 | 36.72 | 8727 | 1677526 | 192.22 | 86 | 56 | 2690 |
|  | TD-50 | 1352965 | 182984 | 13.53 | 17.88 | 44475 | 10216945 | 229.72 | 136 | 56 | 4536 |
| *Dysdercus suturellus* | Standard | 1041752 | 485822 | 46.64 | 44.42 | 7264 | 3328375 | 458.2 | 341.5 | 56 | 3946 |
|  | TD-60 | 1342251 | 589718 | 43.94 | 39.29 | 15153 | 6998339 | 461.85 | 362 | 56 | 4300 |
| *Holhymenia* sp. | Standard | 1270723 | 595948 | 46.90 | 46.47 | 11490 | 3784462 | 329.37 | 215 | 56 | 3692 |
|  | TD-55 | 1369718 | 418936 | 30.59 | 27.84 | 26996 | 7794267 | 288.72 | 211 | 56 | 5328 |
| *Hypselonotus bitrianguliger* | Standard | 1009005 | 454399 | 45.03 | 41.87 | 8115 | 2750572 | 338.95 | 217 | 56 | 6494 |
|  | TD-55 | 1334691 | 348942 | 26.14 | 25.53 | 25513 | 7676040 | 300.87 | 225 | 56 | 5773 |
| *Hypselonotus lineatus* | Standard | 938236 | 282013 | 30.06 | 37.83 | 8356 | 1910318 | 228.62 | 90 | 56 | 3349 |
|  | TD-55 | 1234652 | 270865 | 21.94 | 21.71 | 24378 | 6963996 | 285.67 | 213 | 56 | 7004 |
| *Leptoglossus clypealis* | Standard | 1265842 | 615297 | 48.61 | 52.07 | 10561 | 3206977 | 303.66 | 193 | 56 | 3518 |
|  | TD-60 | 1404007 | 618494 | 44.05 | 42.33 | 16992 | 5212070 | 306.74 | 220 | 56 | 4772 |
| *Leptoscelis quadrisignatus* | Standard | 1180206 | 654451 | 55.45 | 54.26 | 9507 | 3134759 | 329.73 | 211 | 56 | 3835 |
|  | TD-60 | 1420166 | 763178 | 53.74 | 46.43 | 16038 | 5381561 | 335.55 | 225 | 56 | 4498 |
| *Melanacanthus margineguttatus* | Standard | 1342674 | 589174 | 43.88 | 50.73 | 10033 | 3352344 | 334.13 | 217 | 56 | 3428 |
|  | TD-60 | 1451198 | 634045 | 43.69 | 44.23 | 14176 | 5201947 | 366.95 | 265 | 56 | 4428 |
| *Myla* sp. | Standard | 1159931 | 543563 | 46.86 | 44.15 | 7932 | 3170688 | 399.73 | 263 | 56 | 3200 |
|  | TD-55 | 1298981 | 413165 | 31.81 | 27.39 | 22938 | 7816755 | 340.78 | 253 | 56 | 4118 |
| *Nematopus lepidus* | Standard | 1320616 | 188334 | 14.26 | 27.28 | 19889 | 3173176 | 159.54 | 75 | 56 | 2606 |
|  | TD-55 | 1360762 | 280873 | 20.64 | 24.51 | 27569 | 6588103 | 238.97 | 144 | 56 | 5208 |
| *Neomegalotomus rufipes* | Standard | 1529108 | 616728 | 40.33 | 52.21 | 14721 | 3631875 | 246.71 | 96 | 56 | 4610 |
|  | TD-60 | 1605689 | 637480 | 39.70 | 42.70 | 17424 | 5361172 | 307.69 | 207 | 56 | 4069 |
| *Petascelis remipes* | Standard | 1398376 | 707935 | 50.63 | 56.48 | 10223 | 3948810 | 386.27 | 273 | 56 | 4101 |
|  | TD-50 | 1043457 | 205874 | 19.73 | 17.33 | 32315 | 9325503 | 288.58 | 236 | 56 | 8970 |
| *Physomerus grossipes* | Standard | 1197327 | 327830 | 27.38 | 32.86 | 15501 | 3251965 | 209.8 | 85 | 56 | 3363 |
|  | TD-50 | 1256370 | 95366 | 7.59 | 11.57 | 47635 | 7736243 | 162.41 | 93 | 56 | 13478 |
| *Plapigus abdominalis* | Standard | 908197 | 301385 | 33.19 | 38.36 | 7184 | 1783295 | 248.23 | 101 | 56 | 2890 |
|  | TD-50 | 1421927 | 174465 | 12.27 | 16.97 | 42343 | 8946416 | 211.28 | 137 | 56 | 7083 |
| *Plectropoda* sp. | Standard | 1223284 | 631590 | 51.63 | 57.74 | 8967 | 3061673 | 341.44 | 233 | 56 | 2992 |
|  | TD-55 | 1328198 | 368149 | 27.72 | 29.54 | 30568 | 8127462 | 265.88 | 213 | 56 | 6071 |
| *Spartocera fusca* | Standard | 1262499 | 642129 | 50.86 | 51.44 | 9246 | 4127458 | 446.4 | 331 | 56 | 4557 |
|  | TD-60 | 1357712 | 603270 | 44.43 | 39.85 | 16486 | 6601949 | 400.46 | 297 | 56 | 4622 |
| *Stenocoris* sp. | Standard | 1056011 | 496566 | 47.02 | 43.40 | 7552 | 2624496 | 347.52 | 230 | 56 | 3431 |
|  | TD-50 | 1371135 | 264974 | 19.33 | 20.42 | 37773 | 10128467 | 268.14 | 225 | 56 | 5604 |

Table S10. Summary data for on-target reads and read depth of targeted loci, partitioned based on type of baits used for 24 taxa that had 2,000,000 million raw reads subsampled to equalize sequencing depth across capture conditions. Abbreviations: see Tables S2–S4 and Table 1 of main text.

| **Taxon** | **Target capture protocol** | **No. FRO exon** | **% FRO exon** | **RD exon** | **No. FRO transcript** | **% FRO transcript** | **RD transcript** | **No. FRO**  **C-baits** | **% FRO**  **C-baits** | **RD**  **C-baits** | **No. FRO**  **PC dual** | **% FRO**  **PC dual** | **RD**  **PC dual** | **No. FRO**  **P-baits** | **% FRO**  **P-baits** | **RD**  **P-baits** |
| --- | --- | --- | --- | --- | --- | --- | --- | --- | --- | --- | --- | --- | --- | --- | --- | --- |
| *Acanthocoris sordidus* | Standard | 305490 | 28.40 | 87.79 | 20257 | 1.88 | 79.13 | 325622 | 30.27 | 87.18 | 119378 | 11.10 | 123.10 | 176219 | 16.38 | 28.23 |
|  | TD-50 | 104057 | 8.61 | 33.30 | 7097 | 0.59 | 31.64 | 110826 | 9.17 | 33.13 | 40756 | 3.37 | 46.28 | 108985 | 9.01 | 13.69 |
| *Anasa scorbutica* | Standard | 338770 | 32.30 | 77.53 | 13940 | 1.33 | 57.52 | 352637 | 33.62 | 76.46 | 132221 | 12.61 | 103.90 | 163632 | 15.60 | 25.61 |
|  | TD-60 | 299571 | 25.58 | 67.20 | 18297 | 1.56 | 53.44 | 316733 | 27.04 | 66.05 | 113567 | 9.70 | 88.52 | 209729 | 17.91 | 21.66 |
| *Anisoscelis gradadius* | Standard | 476030 | 32.28 | 113.40 | 19626 | 1.33 | 76.08 | 495640 | 33.61 | 111.13 | 127003 | 8.61 | 134.62 | 184684 | 12.53 | 28.08 |
|  | TD-60 | 401497 | 26.22 | 96.84 | 23284 | 1.52 | 72.91 | 424395 | 27.71 | 95.07 | 128352 | 8.38 | 133.86 | 278109 | 18.16 | 29.47 |
| *Anoplocnemis curvipes* | Standard | 306621 | 25.99 | 72.38 | 26991 | 2.29 | 73.64 | 333463 | 28.26 | 72.47 | 115337 | 9.78 | 97.78 | 154074 | 13.06 | 24.91 |
|  | TD-50 | 65037 | 5.76 | 19.89 | 5795 | 0.51 | 19.89 | 70157 | 6.22 | 19.73 | 25399 | 2.25 | 27.22 | 61954 | 5.49 | 10.35 |
| *Catorhintha texana* | Standard | 246421 | 30.56 | 65.07 | 17251 | 2.14 | 54.12 | 263236 | 32.65 | 64.15 | 95557 | 11.85 | 87.25 | 137107 | 17.00 | 27.67 |
|  | TD-55 | 237754 | 19.61 | 53.65 | 25153 | 2.08 | 60.00 | 259567 | 21.41 | 53.76 | 80941 | 6.68 | 72.05 | 224922 | 18.55 | 21.62 |
| *Chelinidea vittiger* | Standard | 293907 | 26.54 | 76.69 | 14455 | 1.31 | 63.41 | 308191 | 27.83 | 75.93 | 104311 | 9.42 | 107.65 | 139469 | 12.60 | 25.83 |
|  | TD-55 | 182459 | 13.85 | 50.08 | 10772 | 0.82 | 43.45 | 192572 | 14.62 | 49.61 | 66095 | 5.02 | 69.11 | 135872 | 10.31 | 17.59 |
| *Darmistus* sp. | Standard | 98072 | 6.41 | 39.45 | 6277 | 0.41 | 26.62 | 104297 | 6.81 | 38.34 | 33822 | 2.21 | 43.67 | 92038 | 6.01 | 17.62 |
|  | TD-50 | 33159 | 2.23 | 15.38 | 2116 | 0.14 | 12.17 | 35243 | 2.37 | 15.14 | 12456 | 0.84 | 19.99 | 43841 | 2.94 | 8.60 |
| Dasynini sp. | Standard | 126270 | 14.61 | 50.08 | 7587 | 0.88 | 45.96 | 132713 | 15.35 | 49.52 | 44707 | 5.17 | 62.76 | 62826 | 7.27 | 23.04 |
|  | TD-50 | 83934 | 6.20 | 28.72 | 5385 | 0.40 | 32.26 | 88344 | 6.53 | 28.66 | 28512 | 2.11 | 36.42 | 91082 | 6.73 | 13.48 |
| *Dysdercus suturellus* | Standard | 166033 | 15.94 | 71.27 | 10562 | 1.01 | 43.70 | 176340 | 16.93 | 68.68 | 66591 | 6.39 | 79.84 | 280914 | 26.97 | 37.16 |
|  | TD-60 | 178064 | 13.27 | 61.96 | 14932 | 1.11 | 46.25 | 192643 | 14.35 | 60.35 | 86846 | 6.47 | 84.36 | 359503 | 26.78 | 33.05 |
| *Holhymenia* sp. | Standard | 328643 | 25.86 | 74.58 | 20165 | 1.59 | 54.40 | 348406 | 27.42 | 72.96 | 107604 | 8.47 | 106.87 | 212632 | 16.73 | 29.26 |
|  | TD-55 | 212009 | 15.48 | 49.70 | 12156 | 0.89 | 41.11 | 224064 | 16.36 | 49.10 | 73291 | 5.35 | 70.95 | 179673 | 13.12 | 18.18 |
| *Hypselonotus bitrianguliger* | Standard | 254794 | 25.25 | 67.62 | 13502 | 1.34 | 55.59 | 268046 | 26.57 | 66.86 | 92808 | 9.20 | 84.00 | 144825 | 14.35 | 25.04 |
|  | TD-55 | 162365 | 12.17 | 43.38 | 10590 | 0.79 | 44.31 | 171831 | 12.87 | 43.28 | 70076 | 5.25 | 62.10 | 154166 | 11.55 | 17.45 |
| *Hypselonotus lineatus* | Standard | 173435 | 18.49 | 53.20 | 8290 | 0.88 | 40.94 | 181657 | 19.36 | 52.46 | 48488 | 5.17 | 58.69 | 74280 | 7.92 | 23.55 |
|  | TD-55 | 134099 | 10.86 | 35.98 | 9237 | 0.75 | 34.41 | 141969 | 11.50 | 35.65 | 45664 | 3.70 | 48.55 | 112304 | 9.10 | 14.43 |
| *Leptoglossus clypealis* | Standard | 324601 | 25.64 | 81.76 | 19185 | 1.52 | 79.88 | 343456 | 27.13 | 81.60 | 123113 | 9.73 | 121.48 | 213173 | 16.84 | 31.00 |
|  | TD-60 | 310016 | 22.08 | 77.33 | 15884 | 1.13 | 65.07 | 325872 | 23.21 | 76.61 | 111572 | 7.95 | 108.76 | 243726 | 17.36 | 25.47 |
| *Leptoscelis quadrisignatus* | Standard | 345245 | 29.25 | 79.48 | 24352 | 2.06 | 77.89 | 369508 | 31.31 | 79.36 | 132900 | 11.26 | 113.95 | 215053 | 18.22 | 34.69 |
|  | TD-60 | 385144 | 27.12 | 84.59 | 27019 | 1.90 | 76.16 | 411317 | 28.96 | 83.89 | 153357 | 10.80 | 118.50 | 278427 | 19.61 | 26.77 |
| *Melanacanthus margineguttatus* | Standard | 262175 | 19.53 | 80.81 | 17545 | 1.21 | 82.46 | 283420 | 21.11 | 81.07 | 103939 | 7.74 | 114.83 | 268608 | 20.01 | 36.75 |
|  | TD-60 | 270280 | 18.63 | 75.19 | 21432 | 1.60 | 84.75 | 287237 | 19.79 | 75.49 | 101865 | 7.02 | 108.43 | 309448 | 21.32 | 31.83 |
| *Myla* sp. | Standard | 261616 | 22.55 | 72.67 | 18586 | 1.60 | 83.24 | 280084 | 24.15 | 73.27 | 101113 | 8.72 | 104.23 | 209675 | 18.08 | 27.55 |
|  | TD-55 | 174618 | 13.44 | 47.59 | 10642 | 0.82 | 50.55 | 185213 | 14.26 | 47.75 | 69504 | 5.35 | 73.11 | 194949 | 15.01 | 18.71 |
| *Nematopus lepidus* | Standard | 94381 | 7.15 | 37.35 | 5321 | 0.40 | 27.56 | 99644 | 7.55 | 36.64 | 40451 | 3.06 | 51.17 | 66400 | 5.03 | 19.13 |
|  | TD-55 | 121075 | 8.90 | 39.34 | 9867 | 0.73 | 38.67 | 130544 | 9.59 | 39.24 | 46037 | 3.38 | 53.84 | 132617 | 9.75 | 17.50 |
| *Neomegalotomus rufipes* | Standard | 298045 | 19.49 | 86.46 | 22999 | 1.50 | 89.04 | 320832 | 20.98 | 86.61 | 119992 | 7.85 | 121.77 | 276952 | 18.11 | 37.54 |
|  | TD-60 | 277467 | 17.28 | 76.53 | 23559 | 1.47 | 82.53 | 300838 | 18.74 | 76.95 | 124592 | 7.76 | 121.54 | 285894 | 17.81 | 28.25 |
| *Petascelis remipes* | Standard | 371600 | 26.57 | 91.98 | 31487 | 2.25 | 95.20 | 402800 | 28.80 | 92.19 | 140428 | 10.04 | 132.88 | 227048 | 16.24 | 31.30 |
|  | TD-50 | 88943 | 8.52 | 26.76 | 7539 | 0.72 | 26.46 | 96071 | 9.21 | 26.69 | 38082 | 3.65 | 38.06 | 96695 | 9.27 | 12.62 |
| *Physomerus grossipes* | Standard | 166469 | 13.90 | 43.72 | 9054 | 0.76 | 36.34 | 175483 | 14.66 | 43.25 | 74030 | 6.18 | 65.49 | 114823 | 9.59 | 22.67 |
|  | TD-50 | 45048 | 3.59 | 15.83 | 2530 | 0.20 | 13.22 | 47570 | 3.79 | 15.65 | 19252 | 1.53 | 22.18 | 40676 | 3.24 | 8.98 |
| *Plapigus abdominalis* | Standard | 171899 | 18.93 | 51.83 | 7885 | 0.87 | 43.03 | 179729 | 19.79 | 51.37 | 58065 | 6.39 | 62.23 | 88248 | 9.72 | 25.25 |
|  | TD-50 | 86884 | 6.11 | 26.00 | 4023 | 0.28 | 22.51 | 90632 | 6.37 | 25.78 | 23469 | 1.65 | 30.58 | 82265 | 5.79 | 12.80 |
| *Plectropoda* sp. | Standard | 352313 | 28.80 | 95.23 | 25323 | 2.07 | 89.63 | 377557 | 30.86 | 94.82 | 137693 | 11.26 | 142.54 | 183740 | 15.02 | 30.25 |
|  | TD-55 | 175363 | 13.20 | 51.19 | 11802 | 0.89 | 48.80 | 186974 | 14.08 | 51.02 | 70891 | 5.34 | 79.06 | 146914 | 11.06 | 18.14 |
| *Spartocera fusca* | Standard | 335775 | 26.60 | 85.88 | 27072 | 2.14 | 85.82 | 362818 | 28.74 | 85.87 | 131198 | 10.39 | 124.06 | 197780 | 15.67 | 27.31 |
|  | TD-60 | 284711 | 20.97 | 71.01 | 20735 | 1.53 | 66.53 | 305359 | 22.49 | 70.68 | 112038 | 8.25 | 109.08 | 230002 | 16.94 | 23.32 |
| *Stenocoris* sp. | Standard | 220887 | 20.92 | 68.15 | 15802 | 1.50 | 51.42 | 236508 | 22.40 | 66.67 | 94879 | 8.99 | 96.14 | 218701 | 20.71 | 30.83 |
|  | TD-50 | 112317 | 8.19 | 33.74 | 10143 | 0.74 | 30.43 | 119884 | 8.74 | 33.10 | 42875 | 3.13 | 43.66 | 151145 | 11.02 | 16.79 |

Table S11. Summary data for captured loci targeted by exon-derived baits (376 loci targeted) from 24 taxa that had 2,000,000 million raw reads subsampled to equalize sequencing depth across capture conditions. Abbreviations: see Tables S2–S4.

| **Taxon** | **Target capture protocol** | **No. loci** | **% loci recovered** | **Mean locus length** | **Median locus length** | **Min locus length** | **Max locus length** | **No. putative paralogs** |
| --- | --- | --- | --- | --- | --- | --- | --- | --- |
| *Acanthocoris sordidus* | Standard | 331 | 88.03 | 1028.64 | 965 | 216 | 2693 | 24 |
|  | TD-50 | 328 | 87.23 | 904.02 | 821 | 219 | 2704 | 30 |
| *Anasa scorbutica* | Standard | 338 | 89.89 | 1229.72 | 1188 | 273 | 2925 | 21 |
|  | TD-60 | 338 | 89.89 | 1279.75 | 1209 | 210 | 3003 | 29 |
| *Anisoscelis gradadius* | Standard | 337 | 89.63 | 1171.20 | 1121 | 213 | 2994 | 26 |
|  | TD-60 | 335 | 89.10 | 1206.15 | 1122 | 277 | 3521 | 32 |
| *Anoplocnemis curvipes* | Standard | 328 | 87.23 | 1228.81 | 1181 | 214 | 2652 | 25 |
|  | TD-50 | 314 | 83.51 | 962.59 | 906.5 | 208 | 2587 | 26 |
| *Catorhintha texana* | Standard | 310 | 82.45 | 1159.29 | 1115.5 | 218 | 2817 | 19 |
|  | TD-55 | 327 | 86.97 | 1277.37 | 1237 | 219 | 3383 | 26 |
| *Chelinidea vittiger* | Standard | 335 | 89.10 | 1108.05 | 1066 | 222 | 2788 | 27 |
|  | TD-55 | 339 | 90.16 | 1042.68 | 989 | 213 | 3082 | 30 |
| *Darmistus* sp. | Standard | 273 | 72.61 | 887.83 | 854 | 208 | 2254 | 21 |
|  | TD-50 | 275 | 73.14 | 747.25 | 676 | 207 | 2174 | 25 |
| Dasynini sp. | Standard | 299 | 79.52 | 800.82 | 717 | 220 | 2690 | 21 |
|  | TD-50 | 320 | 85.11 | 830.47 | 754 | 221 | 2495 | 30 |
| *Dysdercus suturellus* | Standard | 214 | 56.91 | 1056.13 | 1030.5 | 213 | 3946 | 17 |
|  | TD-60 | 257 | 68.35 | 1092.52 | 1068 | 223 | 3986 | 21 |
| *Holhymenia* sp. | Standard | 337 | 89.63 | 1241.29 | 1202 | 208 | 2927 | 23 |
|  | TD-55 | 339 | 90.16 | 1196.98 | 1141 | 229 | 3260 | 28 |
| *Hypselonotus bitrianguliger* | Standard | 330 | 87.77 | 1093.94 | 1045 | 221 | 2726 | 25 |
|  | TD-55 | 334 | 88.83 | 1075.74 | 1044 | 218 | 2822 | 32 |
| *Hypselonotus lineatus* | Standard | 310 | 82.45 | 981.79 | 944.5 | 208 | 2502 | 23 |
|  | TD-55 | 329 | 87.50 | 1069.90 | 1023 | 207 | 2970 | 29 |
| *Leptoglossus clypealis* | Standard | 336 | 89.36 | 1150.65 | 1137.5 | 227 | 2848 | 20 |
|  | TD-60 | 342 | 90.96 | 1143.71 | 1115.5 | 236 | 2838 | 27 |
| *Leptoscelis quadrisignatus* | Standard | 345 | 91.76 | 1229.23 | 1195 | 218 | 3055 | 17 |
|  | TD-60 | 342 | 90.96 | 1311.28 | 1259.5 | 250 | 3067 | 26 |
| *Melanacanthus margineguttatus* | Standard | 280 | 74.47 | 1121.79 | 1104 | 240 | 3018 | 30 |
|  | TD-60 | 306 | 81.38 | 1144.26 | 1097.5 | 213 | 2946 | 34 |
| *Myla* sp. | Standard | 308 | 81.91 | 1144.10 | 1115.5 | 242 | 3150 | 25 |
|  | TD-55 | 321 | 85.37 | 1121.94 | 1070 | 214 | 3195 | 30 |
| *Nematopus lepidus* | Standard | 301 | 80.05 | 793.30 | 746 | 206 | 2206 | 18 |
|  | TD-55 | 322 | 85.64 | 900.20 | 862 | 209 | 2606 | 26 |
| *Neomegalotomus rufipes* | Standard | 297 | 78.99 | 1112.13 | 1077 | 213 | 3028 | 23 |
|  | TD-60 | 308 | 81.91 | 1157.59 | 1117 | 220 | 3281 | 35 |
| *Petascelis remipes* | Standard | 340 | 90.43 | 1158.57 | 1110 | 210 | 2724 | 23 |
|  | TD-50 | 325 | 86.44 | 957.87 | 898 | 211 | 4002 | 27 |
| *Physomerus grossipes* | Standard | 327 | 86.97 | 1082.01 | 1061 | 207 | 2645 | 21 |
|  | TD-50 | 305 | 81.12 | 840.84 | 791 | 218 | 2439 | 23 |
| *Plapigus abdominalis* | Standard | 305 | 81.12 | 1040.83 | 1009 | 211 | 2724 | 21 |
|  | TD-50 | 319 | 84.84 | 949.54 | 882 | 214 | 2690 | 28 |
| *Plectropoda* sp. | Standard | 330 | 87.77 | 1075.37 | 1050.5 | 207 | 2501 | 28 |
|  | TD-55 | 332 | 88.30 | 1000.86 | 936 | 240 | 2724 | 30 |
| *Spartocera fusca* | Standard | 327 | 86.97 | 1174.35 | 1140 | 238 | 2829 | 26 |
|  | TD-60 | 339 | 90.16 | 1170.64 | 1113 | 213 | 2818 | 28 |
| *Stenocoris* sp. | Standard | 296 | 78.72 | 1041.67 | 1043.5 | 219 | 3062 | 25 |
|  | TD-50 | 313 | 83.24 | 927.77 | 866 | 210 | 2424 | 25 |

Table S12. Summary data for captured loci targeted by transcript-derived baits (58 loci targeted) from 24 taxa that had 2,000,000 million raw reads subsampled to equalize sequencing depth across capture conditions. Abbreviations: see Tables S2–S4.

| **Taxon** | **Target capture protocol** | **No. loci** | **% loci recovered** | **Mean locus length** | **Median locus length** | **Min locus length** | **Max locus length** | **No. putative paralogs** |
| --- | --- | --- | --- | --- | --- | --- | --- | --- |
| *Acanthocoris sordidus* | Standard | 22 | 37.93 | 1137.09 | 1206.5 | 211 | 1754 | 36 |
|  | TD-50 | 21 | 36.21 | 1021.81 | 1024 | 605 | 1693 | 37 |
| *Anasa scorbutica* | Standard | 17 | 29.31 | 1382.41 | 1322 | 540 | 2454 | 41 |
|  | TD-60 | 20 | 34.48 | 1650.55 | 1468 | 870 | 3781 | 38 |
| *Anisoscelis gradadius* | Standard | 19 | 32.76 | 1341.63 | 1284 | 645 | 2557 | 39 |
|  | TD-60 | 20 | 34.48 | 1566.70 | 1388 | 894 | 3158 | 38 |
| *Anoplocnemis curvipes* | Standard | 21 | 36.21 | 1697.52 | 1597 | 934 | 3196 | 36 |
|  | TD-50 | 22 | 37.93 | 1257.77 | 1190 | 540 | 2198 | 35 |
| *Catorhintha texana* | Standard | 21 | 36.21 | 1436.57 | 1366 | 681 | 3045 | 35 |
|  | TD-55 | 23 | 39.66 | 1670.74 | 1488 | 627 | 3088 | 34 |
| *Chelinidea vittiger* | Standard | 18 | 31.03 | 1225.22 | 1227.5 | 698 | 2340 | 40 |
|  | TD-55 | 19 | 32.76 | 1229.63 | 1289 | 434 | 2358 | 39 |
| *Darmistus* sp. | Standard | 22 | 37.93 | 1035.91 | 969 | 384 | 2324 | 31 |
|  | TD-50 | 20 | 34.48 | 811.05 | 841.5 | 236 | 2105 | 36 |
| Dasynini sp. | Standard | 20 | 34.48 | 712.20 | 634.5 | 284 | 1144 | 37 |
|  | TD-50 | 16 | 27.59 | 904.06 | 946 | 454 | 1536 | 42 |
| *Dysdercus suturellus* | Standard | 26 | 44.83 | 885.62 | 884.5 | 228 | 1532 | 17 |
|  | TD-60 | 28 | 48.28 | 1105.00 | 1028.5 | 619 | 2012 | 21 |
| *Holhymenia* sp. | Standard | 21 | 36.21 | 1690.52 | 1438 | 814 | 3692 | 37 |
|  | TD-55 | 18 | 31.03 | 1612.00 | 1370 | 805 | 3690 | 40 |
| *Hypselonotus bitrianguliger* | Standard | 19 | 32.76 | 1220.63 | 1229 | 482 | 2018 | 39 |
|  | TD-55 | 18 | 31.03 | 1178.33 | 1211 | 467 | 2213 | 40 |
| *Hypselonotus lineatus* | Standard | 18 | 31.03 | 1062.00 | 1059.5 | 473 | 1599 | 37 |
|  | TD-55 | 19 | 32.76 | 1265.00 | 1265 | 263 | 2343 | 39 |
| *Leptoglossus clypealis* | Standard | 17 | 29.31 | 1377.59 | 1184 | 806 | 2716 | 41 |
|  | TD-60 | 16 | 27.59 | 1512.56 | 1345 | 993 | 3290 | 42 |
| *Leptoscelis quadrisignatus* | Standard | 20 | 34.48 | 1518.30 | 1422 | 841 | 2142 | 38 |
|  | TD-60 | 21 | 36.21 | 1641.00 | 1517 | 1110 | 2295 | 37 |
| *Melanacanthus margineguttatus* | Standard | 19 | 32.76 | 1291.32 | 1245 | 641 | 2520 | 39 |
|  | TD-60 | 15 | 25.86 | 1378.73 | 1383 | 752 | 2399 | 43 |
| *Myla* sp. | Standard | 17 | 29.31 | 1286.59 | 1280 | 846 | 1912 | 41 |
|  | TD-55 | 16 | 27.59 | 1289.06 | 1353.5 | 725 | 2098 | 42 |
| *Nematopus lepidus* | Standard | 22 | 37.93 | 834.77 | 808.5 | 221 | 1426 | 35 |
|  | TD-55 | 20 | 34.48 | 1168.35 | 1039.5 | 596 | 2417 | 38 |
| *Neomegalotomus rufipes* | Standard | 18 | 31.03 | 1405.44 | 1365 | 699 | 2393 | 40 |
|  | TD-60 | 17 | 29.31 | 1654.06 | 1423 | 877 | 3261 | 41 |
| *Petascelis remipes* | Standard | 23 | 39.66 | 1407.48 | 1447 | 1000 | 1996 | 34 |
|  | TD-50 | 23 | 39.66 | 1135.39 | 1119 | 299 | 1856 | 35 |
| *Physomerus grossipes* | Standard | 21 | 36.21 | 1145.90 | 1247 | 328 | 1903 | 37 |
|  | TD-50 | 20 | 34.48 | 945.90 | 1013.5 | 346 | 1787 | 37 |
| *Plapigus abdominalis* | Standard | 17 | 29.31 | 1023.94 | 1069 | 468 | 1721 | 39 |
|  | TD-50 | 15 | 25.86 | 1091.53 | 1100 | 668 | 1491 | 41 |
| *Plectropoda* sp. | Standard | 20 | 34.48 | 1382.15 | 1339.5 | 693 | 2453 | 38 |
|  | TD-55 | 18 | 31.03 | 1312.28 | 1197 | 837 | 2538 | 40 |
| *Spartocera fusca* | Standard | 21 | 36.21 | 1482.48 | 1366 | 884 | 2776 | 36 |
|  | TD-60 | 21 | 36.21 | 1462.48 | 1332 | 981 | 2852 | 37 |
| *Stenocoris* sp. | Standard | 22 | 37.93 | 1342.36 | 1191 | 652 | 3203 | 33 |
|  | TD-50 | 24 | 41.38 | 1105.08 | 1095.5 | 210 | 1878 | 34 |

Table S13. Summary data for captured loci targeted by Pentatomomorpha-Coreoidea dual baits (103 loci targeted) from 24 taxa that had 2,000,000 million raw reads subsampled to equalize sequencing depth across capture conditions. Abbreviations: see Tables S2–S4.

| **Taxon** | **Target capture protocol** | **No. loci** | **% loci recovered** | **Mean locus length** | **Median locus length** | **Min locus length** | **Max locus length** | **No. putative paralogs** |
| --- | --- | --- | --- | --- | --- | --- | --- | --- |
| *Acanthocoris sordidus* | Standard | 75 | 72.82 | 1264.04 | 1076 | 247 | 2669 | 28 |
|  | TD-50 | 72 | 69.90 | 1170.68 | 992.5 | 212 | 2752 | 31 |
| *Anasa scorbutica* | Standard | 78 | 75.73 | 1552.23 | 1380.5 | 620 | 4047 | 25 |
|  | TD-60 | 75 | 72.82 | 1645.39 | 1360 | 568 | 4095 | 28 |
| *Anisoscelis gradadius* | Standard | 67 | 65.05 | 1383.01 | 1231 | 519 | 2848 | 36 |
|  | TD-60 | 65 | 63.11 | 1453.42 | 1224 | 521 | 2838 | 38 |
| *Anoplocnemis curvipes* | Standard | 75 | 72.82 | 1528.24 | 1427 | 317 | 3045 | 28 |
|  | TD-50 | 72 | 69.90 | 1214.24 | 1056.5 | 244 | 2667 | 29 |
| *Catorhintha texana* | Standard | 76 | 73.79 | 1365.95 | 1271.5 | 227 | 2858 | 24 |
|  | TD-55 | 71 | 68.93 | 1485.23 | 1303 | 274 | 3340 | 22 |
| *Chelinidea vittiger* | Standard | 68 | 66.02 | 1391.94 | 1213.5 | 451 | 2821 | 35 |
|  | TD-55 | 69 | 66.99 | 1357.03 | 1115 | 355 | 2859 | 34 |
| *Darmistus* sp. | Standard | 69 | 66.99 | 1101.10 | 1012 | 206 | 2717 | 28 |
|  | TD-50 | 59 | 57.28 | 998.32 | 839 | 222 | 2637 | 36 |
| Dasynini sp. | Standard | 72 | 69.90 | 954.04 | 773.5 | 210 | 2484 | 27 |
|  | TD-50 | 68 | 66.02 | 1104.66 | 863.5 | 398 | 2641 | 31 |
| *Dysdercus suturellus* | Standard | 64 | 62.14 | 1209.59 | 1142 | 265 | 2928 | 27 |
|  | TD-60 | 69 | 66.99 | 1434.58 | 1255 | 336 | 3102 | 28 |
| *Holhymenia* sp. | Standard | 65 | 63.11 | 1498.11 | 1261 | 708 | 3017 | 38 |
|  | TD-55 | 64 | 62.14 | 1527.47 | 1350 | 551 | 3174 | 39 |
| *Hypselonotus bitrianguliger* | Standard | 79 | 76.70 | 1350.67 | 1168 | 247 | 2947 | 23 |
|  | TD-55 | 78 | 75.73 | 1377.38 | 1204 | 260 | 2961 | 25 |
| *Hypselonotus lineatus* | Standard | 70 | 67.96 | 1127.09 | 996.5 | 284 | 2872 | 30 |
|  | TD-55 | 69 | 66.99 | 1294.71 | 1126 | 342 | 3059 | 33 |
| *Leptoglossus clypealis* | Standard | 69 | 66.99 | 1437.13 | 1209 | 343 | 2766 | 34 |
|  | TD-60 | 67 | 65.05 | 1509.97 | 1217 | 506 | 4132 | 36 |
| *Leptoscelis quadrisignatus* | Standard | 74 | 71.84 | 1535.64 | 1387.5 | 275 | 3004 | 29 |
|  | TD-60 | 77 | 74.76 | 1654.53 | 1434 | 493 | 3224 | 26 |
| *Melanacanthus margineguttatus* | Standard | 60 | 58.25 | 1474.12 | 1299.5 | 263 | 3303 | 37 |
|  | TD-60 | 61 | 59.22 | 1510.33 | 1450 | 323 | 3512 | 39 |
| *Myla* sp. | Standard | 67 | 65.05 | 1410.87 | 1245 | 304 | 3200 | 32 |
|  | TD-55 | 68 | 66.02 | 1364.68 | 1175.5 | 403 | 3148 | 35 |
| *Nematopus lepidus* | Standard | 74 | 71.84 | 1031.34 | 870.5 | 236 | 2606 | 23 |
|  | TD-55 | 71 | 68.93 | 1148.61 | 967 | 216 | 2848 | 32 |
| *Neomegalotomus rufipes* | Standard | 74 | 71.84 | 1297.12 | 1136.5 | 211 | 3260 | 27 |
|  | TD-60 | 71 | 68.93 | 1417.37 | 1381 | 303 | 3345 | 32 |
| *Petascelis remipes* | Standard | 71 | 68.93 | 1458.01 | 1279 | 505 | 2930 | 57 |
|  | TD-50 | 73 | 70.87 | 1250.41 | 1149 | 261 | 2730 | 62 |
| *Physomerus grossipes* | Standard | 75 | 72.82 | 1369.55 | 1285 | 217 | 2865 | 26 |
|  | TD-50 | 73 | 70.87 | 1098.60 | 887 | 208 | 2697 | 27 |
| *Plapigus abdominalis* | Standard | 74 | 71.84 | 1205.22 | 1078 | 219 | 2844 | 26 |
|  | TD-50 | 64 | 62.14 | 1110.86 | 929.5 | 280 | 2749 | 37 |
| *Plectropoda* sp. | Standard | 72 | 69.90 | 1315.51 | 1101 | 430 | 2992 | 31 |
|  | TD-55 | 68 | 66.02 | 1289.60 | 1071.5 | 412 | 3102 | 35 |
| *Spartocera fusca* | Standard | 69 | 66.99 | 1511.13 | 1335 | 613 | 2922 | 32 |
|  | TD-60 | 67 | 65.05 | 1521.00 | 1314 | 597 | 2993 | 34 |
| *Stenocoris* sp. | Standard | 74 | 71.84 | 1269.51 | 1170 | 229 | 2797 | 28 |
|  | TD-50 | 69 | 66.99 | 1245.93 | 1074 | 302 | 4794 | 33 |

Table S14. Summary data for captured loci targeted by Pentatomomorpha-derived baits (2566 loci targeted) from 24 taxa that had 2,000,000 million raw reads subsampled to equalize sequencing depth across capture conditions. Abbreviations: see Tables S2–S4.

| **Taxon** | **Target capture protocol** | **No. loci** | **% loci recovered** | **Mean locus length** | **Median locus length** | **Min locus length** | **Max locus length** | **No. putative paralogs** |
| --- | --- | --- | --- | --- | --- | --- | --- | --- |
| *Acanthocoris sordidus* | Standard | 952 | 37.10 | 636.19 | 622 | 206 | 2349 | 3 |
|  | TD-50 | 1269 | 49.45 | 562.01 | 546 | 207 | 2256 | 6 |
| *Anasa scorbutica* | Standard | 836 | 32.58 | 706.91 | 684 | 206 | 2969 | 8 |
|  | TD-60 | 1251 | 48.75 | 732.77 | 706 | 206 | 3781 | 13 |
| *Anisoscelis gradadius* | Standard | 890 | 34.68 | 686.20 | 660.5 | 161 | 2994 | 14 |
|  | TD-60 | 1251 | 48.75 | 703.22 | 682 | 206 | 3521 | 18 |
| *Anoplocnemis curvipes* | Standard | 799 | 31.14 | 740.13 | 727 | 206 | 2852 | 9 |
|  | TD-50 | 928 | 36.17 | 561.69 | 505 | 206 | 2761 | 9 |
| *Catorhintha texana* | Standard | 686 | 26.73 | 668.41 | 607.5 | 206 | 3358 | 3 |
|  | TD-55 | 1224 | 47.70 | 772.61 | 756 | 207 | 3518 | 14 |
| *Chelinidea vittiger* | Standard | 810 | 31.57 | 621.49 | 560 | 206 | 2960 | 6 |
|  | TD-55 | 1236 | 48.17 | 574.41 | 521 | 206 | 3082 | 14 |
| *Darmistus* sp. | Standard | 814 | 31.72 | 610.45 | 590.5 | 206 | 2717 | 6 |
|  | TD-50 | 952 | 37.10 | 484.43 | 459 | 206 | 2637 | 12 |
| Dasynini sp. | Standard | 513 | 19.99 | 484.46 | 437 | 207 | 2077 | 6 |
|  | TD-50 | 1109 | 43.22 | 521.13 | 490 | 206 | 3491 | 14 |
| *Dysdercus suturellus* | Standard | 907 | 35.35 | 801.02 | 788 | 208 | 2928 | 11 |
|  | TD-60 | 1268 | 49.42 | 831.16 | 832.5 | 209 | 3102 | 32 |
| *Holhymenia* sp. | Standard | 895 | 34.88 | 741.02 | 713 | 212 | 2927 | 6 |
|  | TD-55 | 1253 | 48.83 | 711.63 | 684 | 206 | 3197 | 8 |
| *Hypselonotus bitrianguliger* | Standard | 833 | 32.46 | 661.27 | 642 | 206 | 2534 | 6 |
|  | TD-55 | 1237 | 48.21 | 664.83 | 626 | 206 | 2961 | 15 |
| *Hypselonotus lineatus* | Standard | 535 | 20.85 | 559.29 | 496 | 82 | 2503 | 3 |
|  | TD-55 | 1131 | 44.08 | 633.68 | 589 | 206 | 3059 | 13 |
| *Leptoglossus clypealis* | Standard | 919 | 35.81 | 667.08 | 627 | 206 | 3518 | 7 |
|  | TD-60 | 1304 | 50.82 | 690.10 | 664 | 206 | 4772 | 16 |
| *Leptoscelis quadrisignatus* | Standard | 877 | 34.18 | 711.22 | 662 | 206 | 3835 | 4 |
|  | TD-60 | 1321 | 51.48 | 756.50 | 733 | 207 | 3900 | 7 |
| *Melanacanthus margineguttatus* | Standard | 944 | 36.79 | 732.53 | 695 | 206 | 3303 | 34 |
|  | TD-60 | 1254 | 48.87 | 742.95 | 721 | 60 | 3915 | 61 |
| *Myla* sp. | Standard | 976 | 38.04 | 744.81 | 722 | 131 | 3200 | 14 |
|  | TD-55 | 1400 | 54.56 | 708.71 | 693.5 | 206 | 3299 | 20 |
| *Nematopus lepidus* | Standard | 620 | 24.16 | 511.26 | 468.5 | 201 | 2326 | 2 |
|  | TD-55 | 1198 | 46.69 | 568.22 | 544 | 201 | 4046 | 5 |
| *Neomegalotomus rufipes* | Standard | 958 | 37.33 | 704.61 | 674 | 206 | 4610 | 14 |
|  | TD-60 | 1305 | 50.86 | 749.24 | 732 | 206 | 4069 | 29 |
| *Petascelis remipes* | Standard | 967 | 37.69 | 713.34 | 696 | 206 | 2638 | 7 |
|  | TD-50 | 1148 | 44.74 | 588.47 | 563 | 206 | 2804 | 10 |
| *Physomerus grossipes* | Standard | 652 | 25.41 | 652.69 | 612 | 207 | 2442 | 2 |
|  | TD-50 | 791 | 30.83 | 498.24 | 439 | 206 | 2697 | 3 |
| *Plapigus abdominalis* | Standard | 555 | 21.63 | 576.50 | 525 | 206 | 2676 | 4 |
|  | TD-50 | 997 | 38.85 | 540.01 | 493 | 189 | 4302 | 13 |
| *Plectropoda* sp. | Standard | 910 | 35.46 | 637.67 | 616 | 206 | 2607 | 10 |
|  | TD-55 | 1273 | 49.61 | 597.77 | 578 | 206 | 2725 | 13 |
| *Spartocera fusca* | Standard | 987 | 38.46 | 715.46 | 701 | 206 | 2922 | 12 |
|  | TD-60 | 1320 | 51.44 | 730.70 | 724 | 206 | 2993 | 17 |
| *Stenocoris* sp. | Standard | 982 | 38.27 | 663.91 | 638 | 206 | 3062 | 9 |
|  | TD-50 | 1276 | 49.73 | 588.11 | 557.5 | 206 | 5221 | 20 |

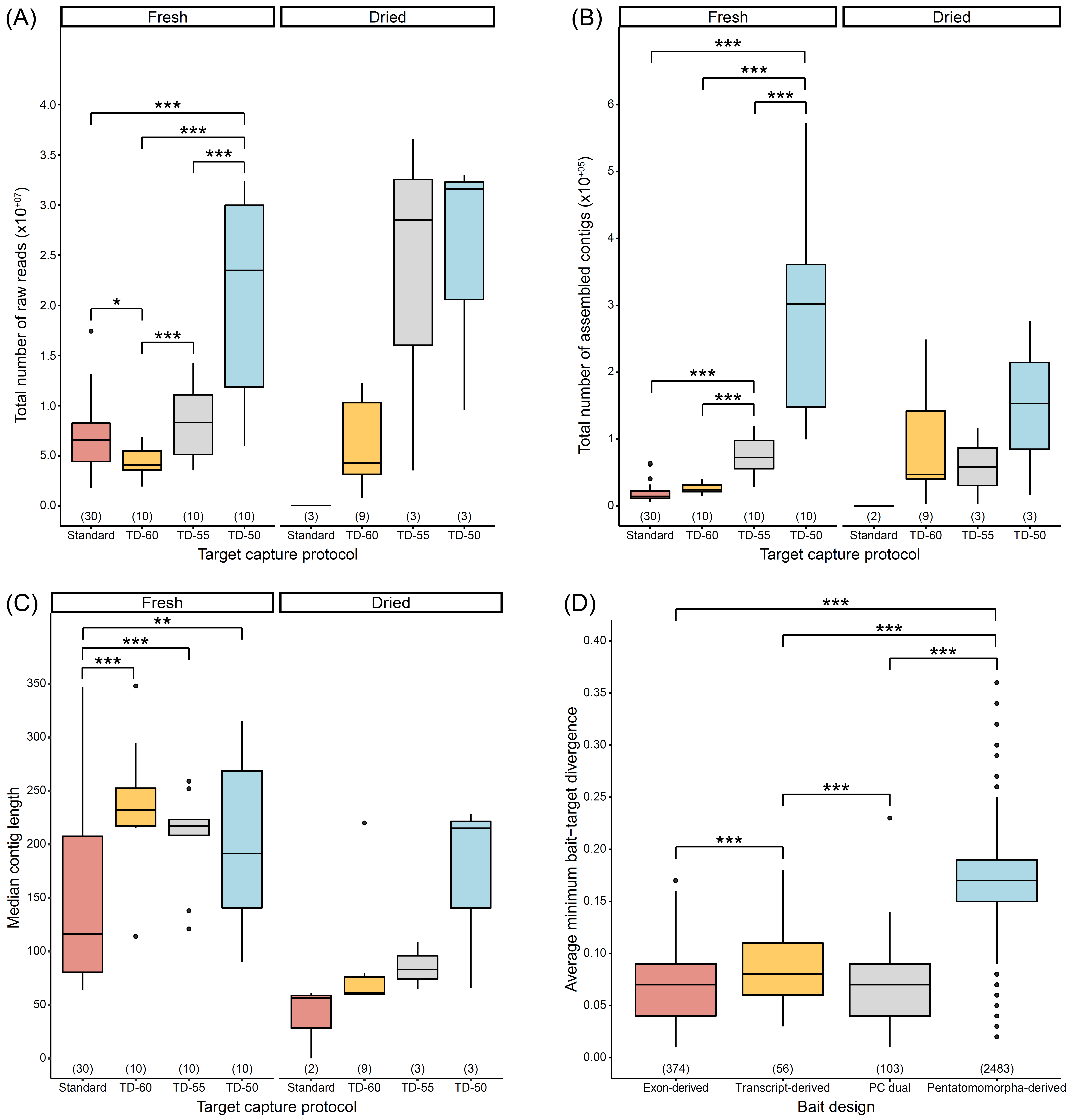

Figure S1. Effects of target capture protocols on the (A) total number of raw reads, (B) assembled contigs, and (C) median contig length, separated by preservation method. (D) Average minimum bait-target divergences by bait design strategy. Numbers in parentheses above x-axis denote sample size. Single, double, and triple asterisks denote statistically significant pairwise comparisons, with p < 0.050, p < 0.010, and p < 0.001, respectively (statistical analyses not performed on dried samples due to low sample sizes). See Tables S2–S4 for abbreviations.

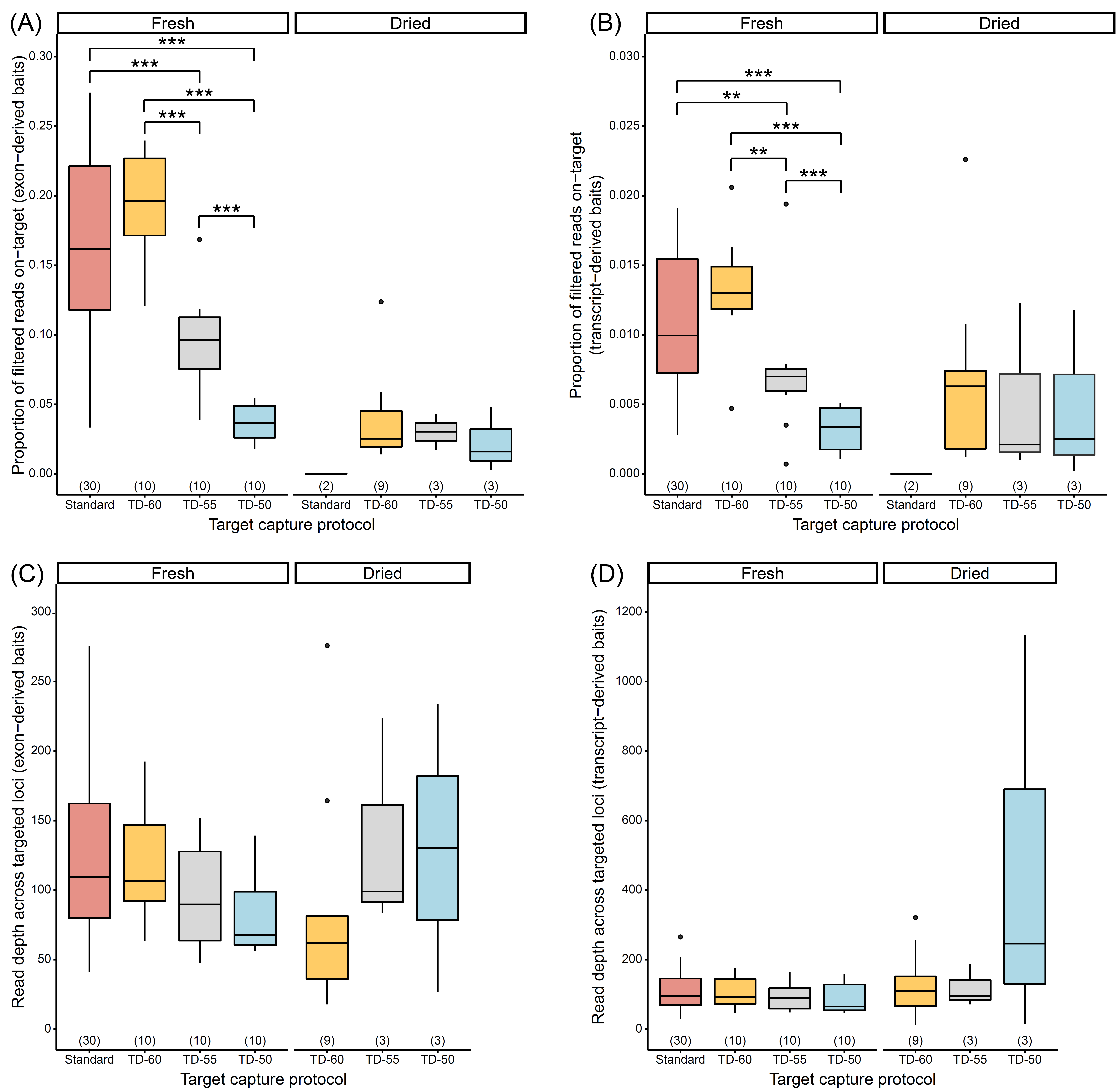

Figure S2. Effects of target capture protocols on (A) the proportion of filtered reads on-target for loci targeted by exon-derived baits, (B) the proportion of filtered reads on-target for loci targeted by transcript-derived baits, (C) read depth across loci targeted by exon-derived baits, and (D) read depth across loci targeted by transcript-derived baits, separated by sample preservation method. Numbers in parentheses above x-axis denote sample size. Double and triple asterisks denote statistically significant pairwise comparisons, with p < 0.010 and p < 0.001, respectively (statistical analyses not performed on dried samples due to low sample sizes). See Tables S2–S4 for abbreviations.

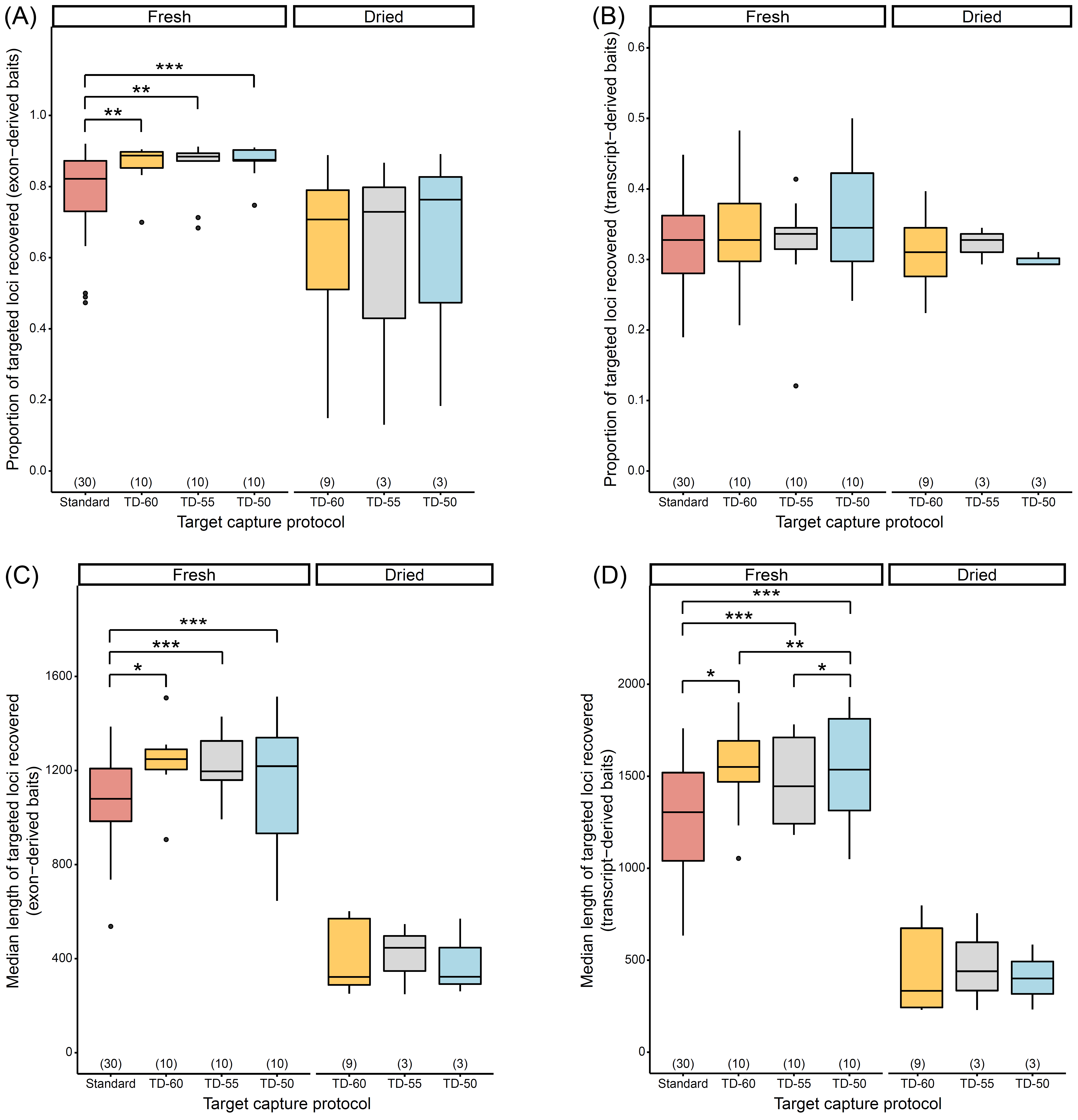

Figure S3. Effects of target capture protocols on (A) the proportion of targeted loci captured by exon-derived baits, (B) the proportion of targeted loci captured by transcript-derived baits, (C) the median lengths of loci targeted by exon-derived baits, and (D) the median lengths of loci targeted by transcript-derived baits, separated by sample preservation method. Numbers in parentheses above x-axis denote sample size. Single, double, and triple asterisks denote statistically significant pairwise comparisons, with p < 0.050, p < 0.010, and p < 0.001, respectively (statistical analyses not performed on dried samples due to low sample sizes). See Tables S2–S4 for abbreviations.

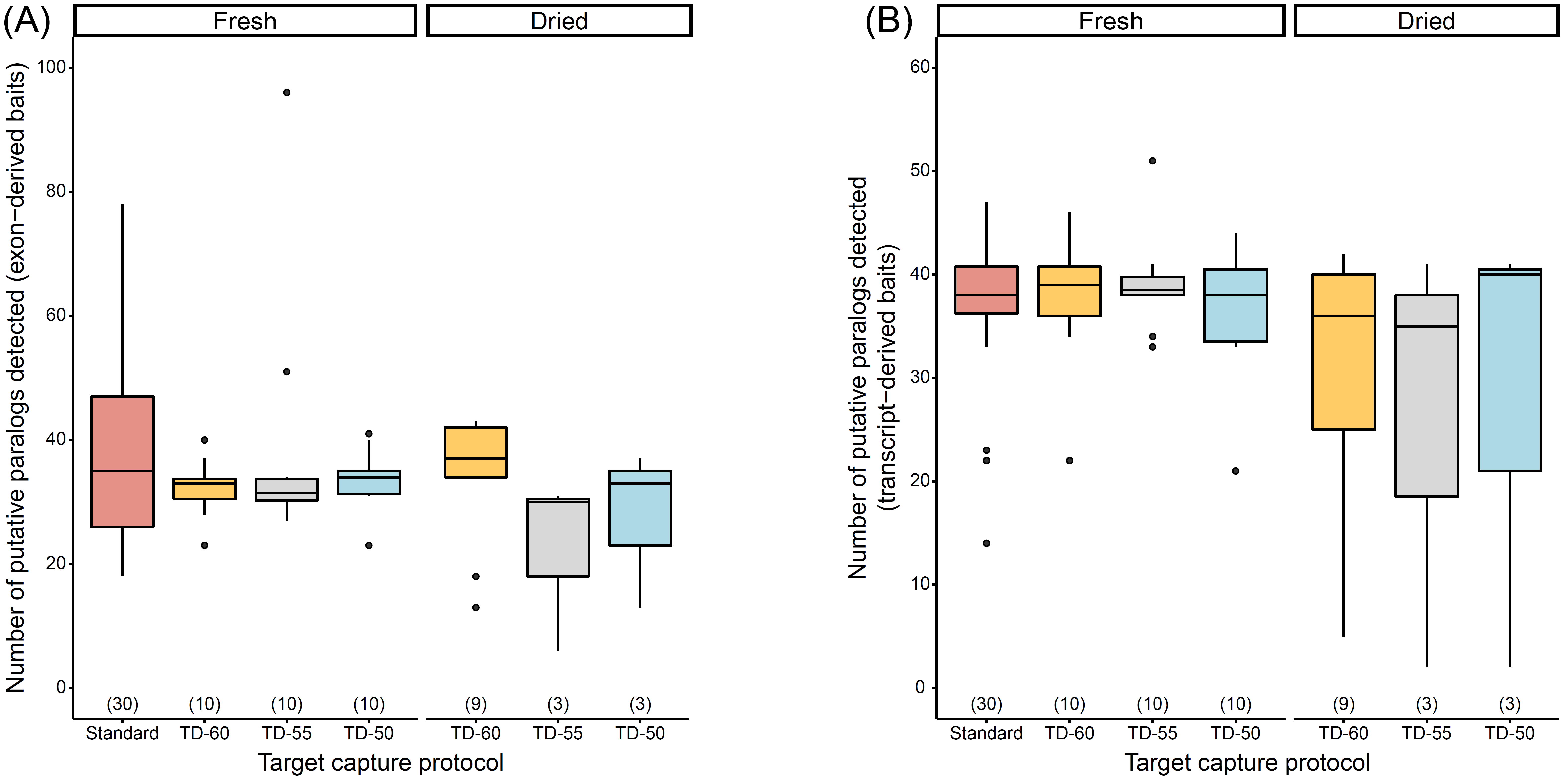

Figure S4. Effects of target capture protocols on the number of putative paralogs of loci captured (A) exon-derived and (B) transcript-derived baits, separated by sample preservation method. Numbers in parentheses above x-axis denote sample size. All pairwise comparisons are not significant (statistical analyses not performed on dried samples due to low sample sizes). See Tables S2–S4 for abbreviations.

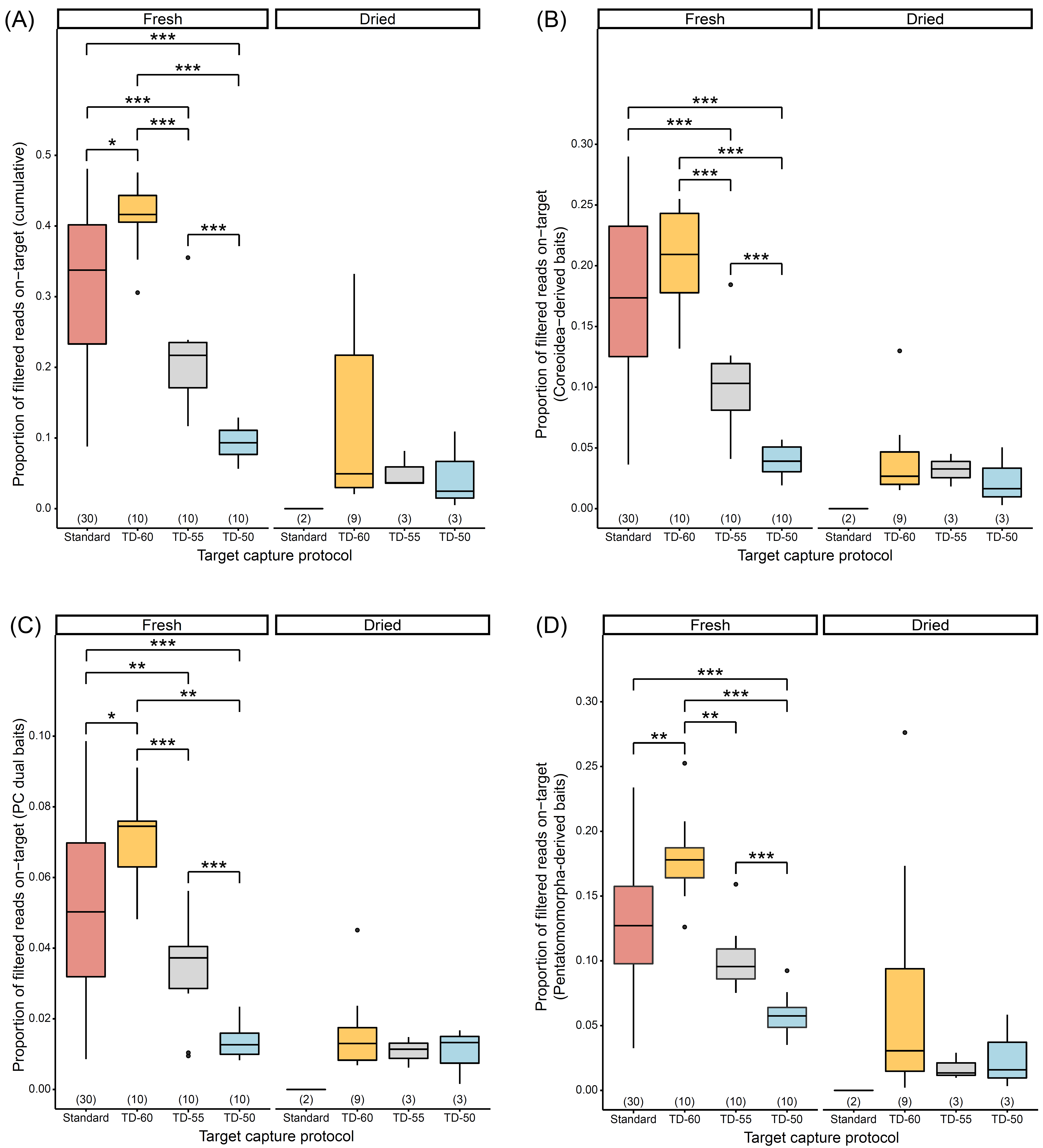

Figure S5. Effects of target capture protocols on the proportion of filtered reads on-target (A) across all targeted loci, (B) loci captured by Coreoidea-derived baits, (C) loci captured by PC dual baits, and (D) loci captured by Pentatomomorpha-derived baits, separated by preservation method. Numbers in parentheses above x-axis denote sample size. Single, double, and triple asterisks denote statistically significant pairwise comparisons, with p < 0.050, p < 0.010, and p < 0.001, respectively (statistical analyses not performed on dried samples due to low sample sizes). See Tables S2–S4 for abbreviations.

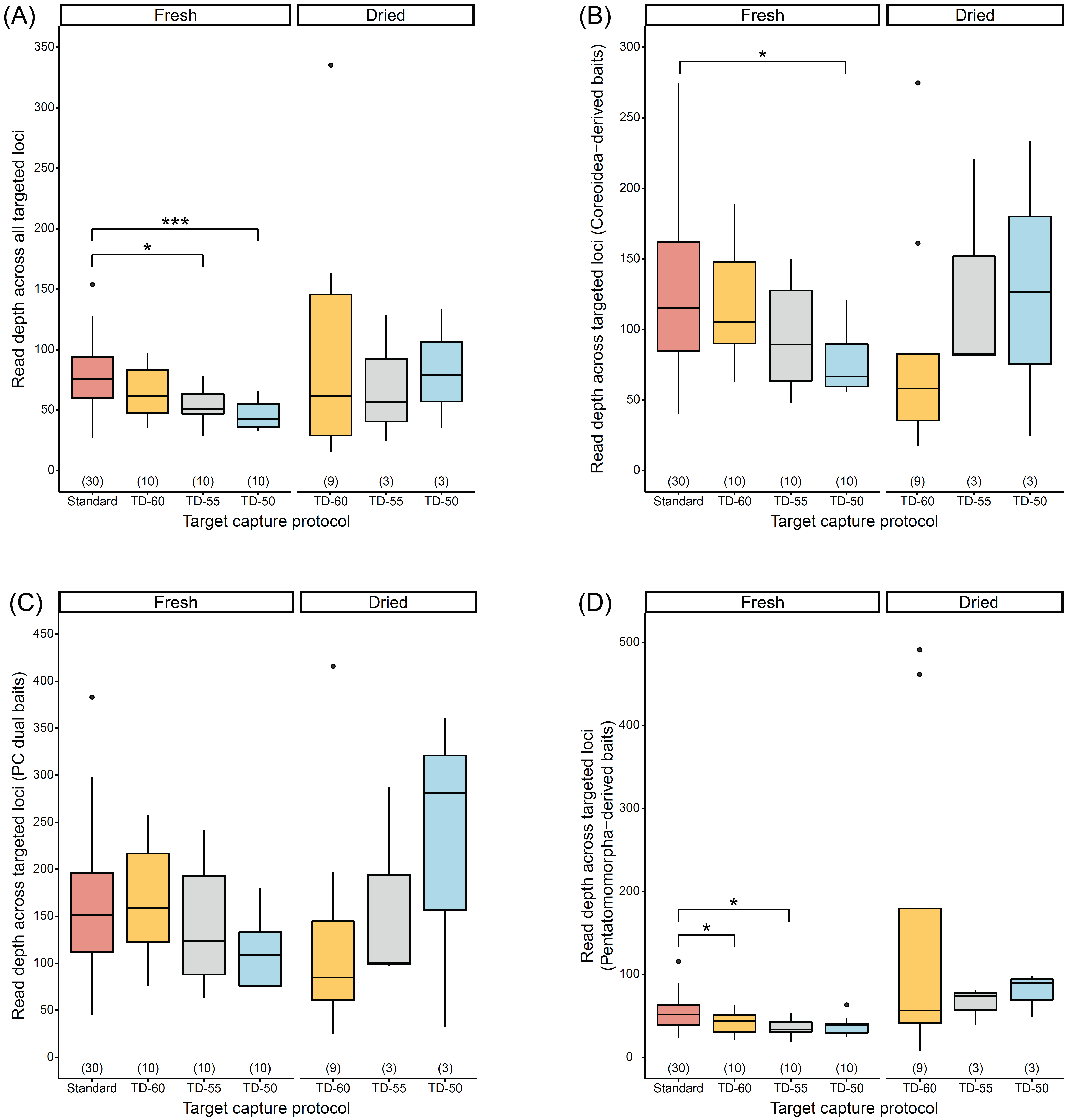

Figure S6. Effects of target capture protocols on read depth (A) across all targeted loci, (B) loci captured by Coreoidea-derived baits, (C) loci captured by PC dual baits, and (D) loci captured by Pentatomomorpha-derived baits, separated by preservation method. Numbers in parentheses above x-axis denote sample size. Single and triple asterisks denote statistically significant pairwise comparisons, with p < 0.050 and p < 0.001, respectively (statistical analyses not performed on dried samples due to low sample sizes). See Tables S2–S4 for abbreviations.

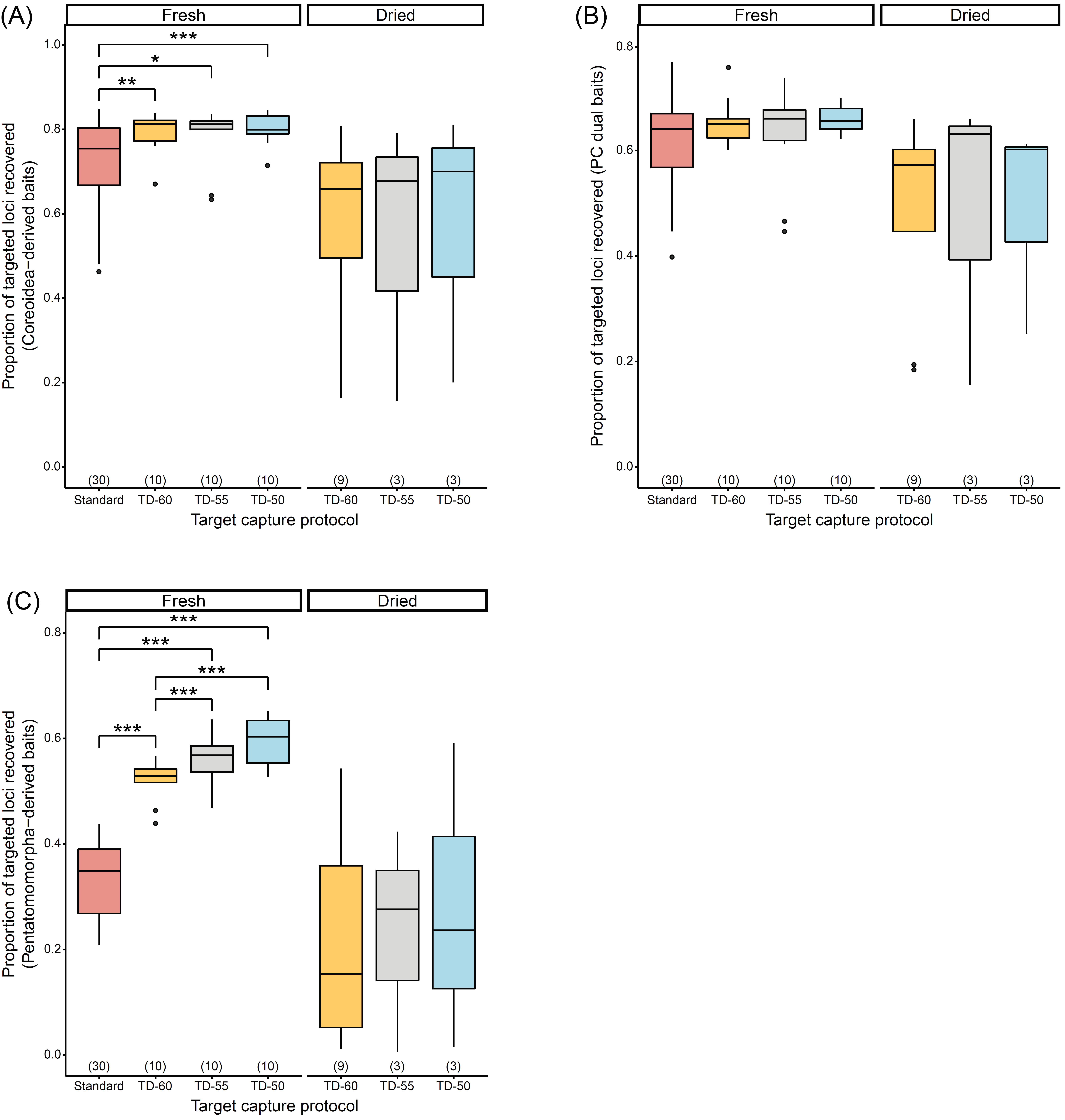

Figure S7. Effects of target capture protocols on the proportion of recovered loci targeted by (A) Coreoidea-derived baits, (C) PC dual baits, and (D) Pentatomomorpha-derived baits, separated by preservation method. Numbers in parentheses above x-axis denote sample size. Single, double, and triple asterisks denote statistically significant pairwise comparisons, with p < 0.050, p < 0.010, and p < 0.001, respectively (statistical analyses not performed on dried samples due to low sample sizes). See Tables S2–S4 for abbreviations.

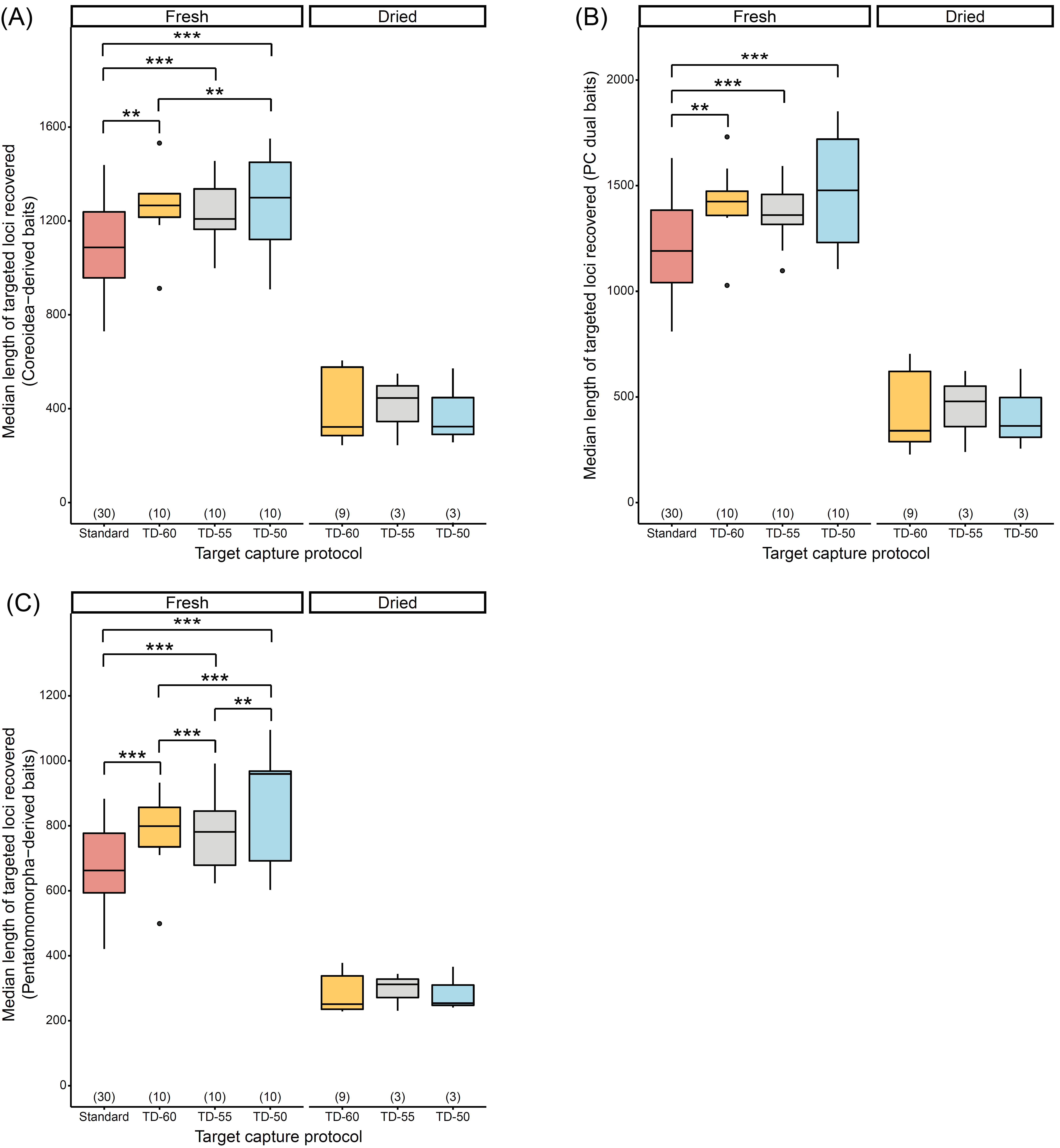

Figure S8. Effects of target capture protocols on the median length of recovered loci targeted by (A) Coreoidea-derived baits, (C) PC dual baits, and (D) Pentatomomorpha-derived baits, separated by preservation method. Numbers in parentheses above x-axis denote sample size. Double and triple asterisks denote statistically significant pairwise comparisons, with p < 0.010 and p < 0.001, respectively (statistical analyses not performed on dried samples due to low sample sizes). See Tables S2–S4 for abbreviations.

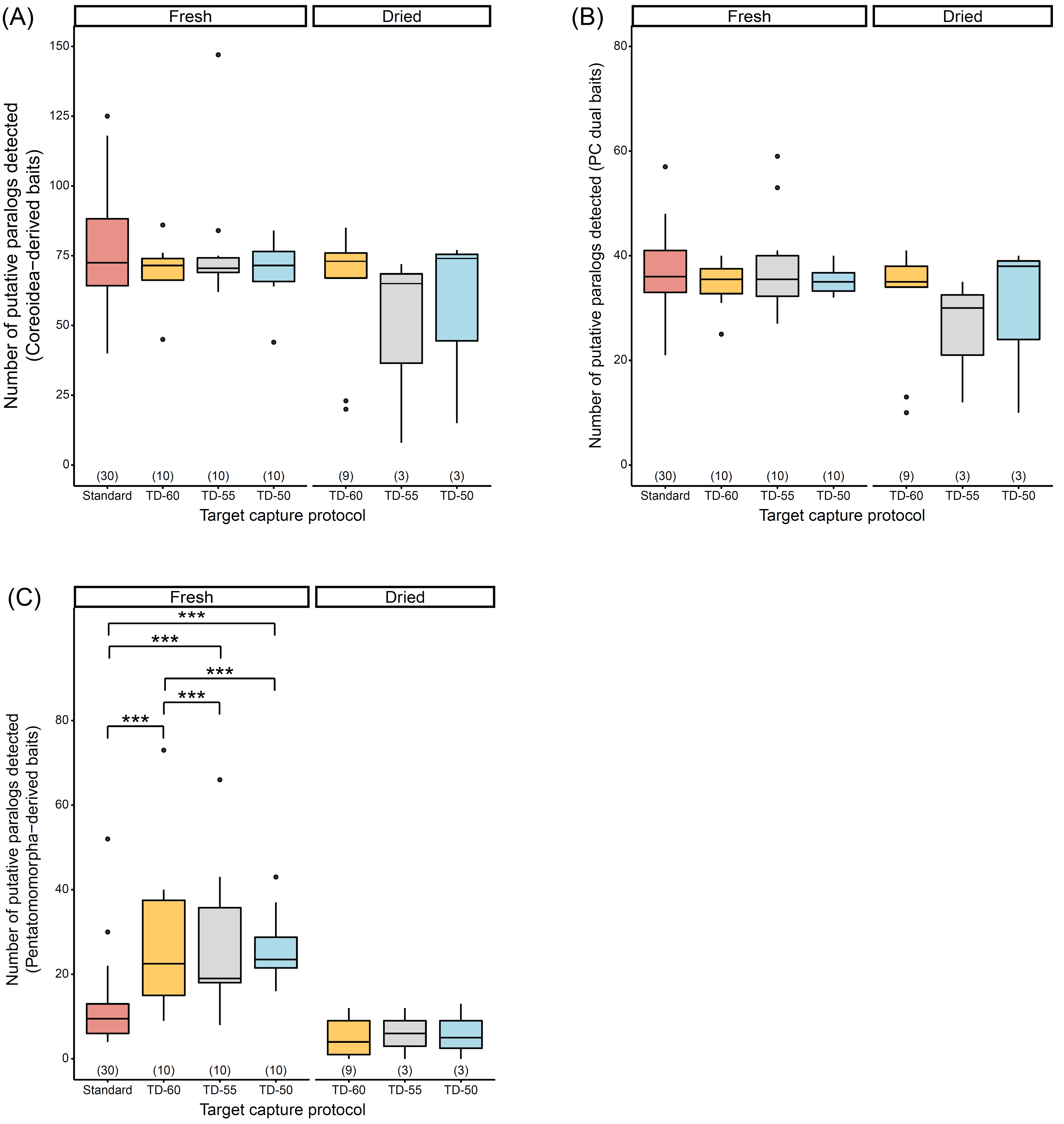

Figure S9. Effects of target capture protocols on the number of putative paralogs of loci captured by (A) Coreoidea-derived baits, (C) PC dual baits, and (D) Pentatomomorpha-derived baits, separated by preservation method. Numbers in parentheses above x-axis denote sample size. Triple asterisks denote statistically significant pairwise comparisons, with p < 0.001 (statistical analyses not performed on dried samples due to low sample sizes). See Tables S2–S4 for abbreviations.

Figure S10. Effects of tiling strategy (Coreoidea-derived ~2x tiling density; Pentatomomorpha-derived ~1.33x tiling density) on average read depth per locus (captured loci exhibit 0.05–0.10 average minimum bait-target divergences for each bait design strategy). Numbers in parentheses above x-axis denote sample size. Triple asterisks denote statistically significant pairwise comparisons, with p < 0.001. See Tables S2–S4 for abbreviations.

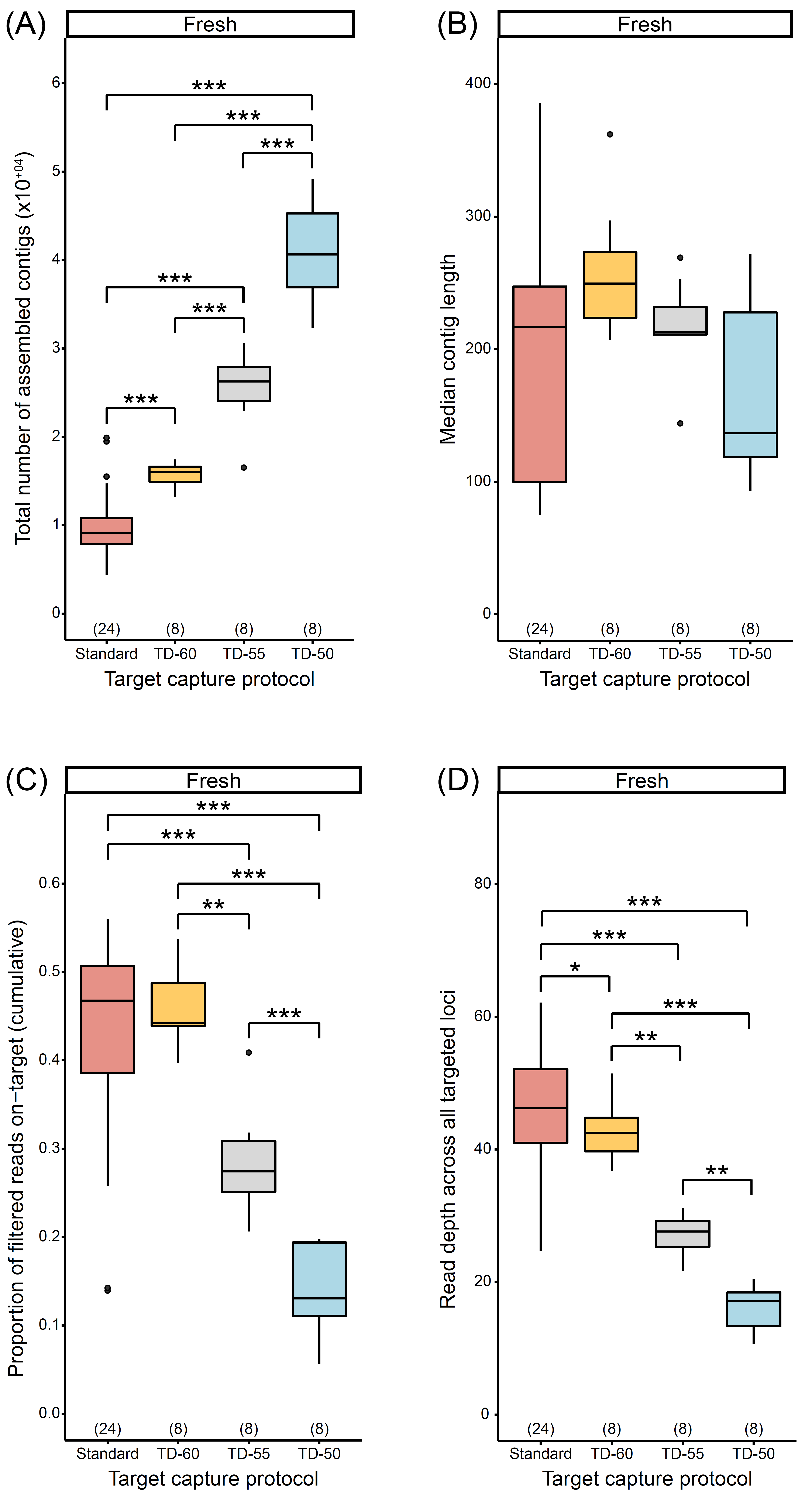

Figure S11. When controlling for sequencing depth, effects of target capture protocols on (A) the number of assembled contigs, (B) median contig length, (C) proportion of filtered reads on-target across all targeted loci, and (D) read depth across all targeted loci for samples preserved fresh. Numbers in parentheses above x-axis denote sample size. Single, double, and triple asterisks denote statistically significant pairwise comparisons, with p < 0.050, p < 0.010, and p < 0.001, respectively. See Tables S2–S4 for abbreviations.

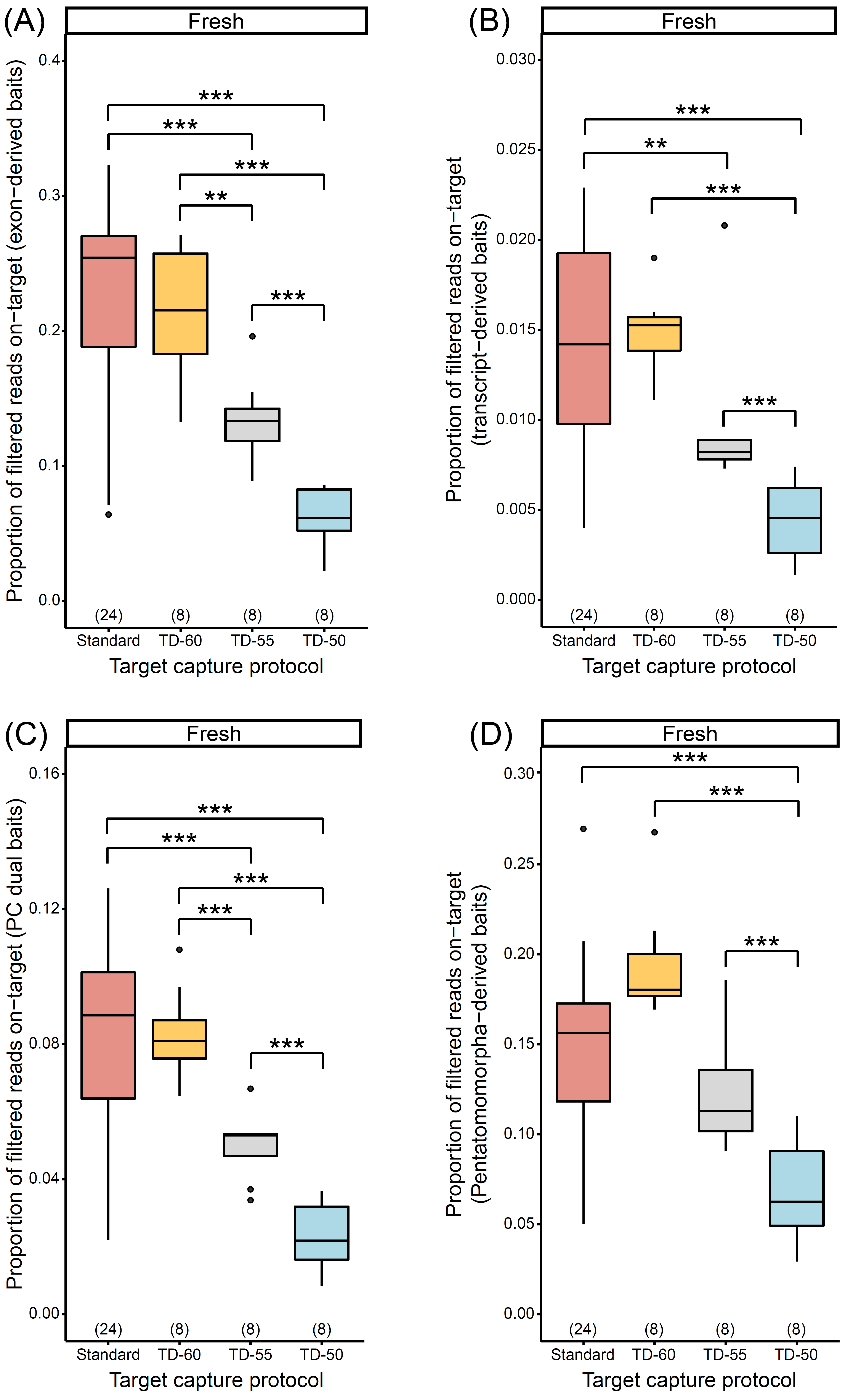

Figure S12. When controlling for sequencing depth, effects of target capture protocols on the proportion of filtered reads on-target for loci targeted by (A) exon-derived baits, (B) transcript-derived baits, (C) PC dual baits, and (D) Pentatomomorpha-derived baits for samples preserved fresh. Numbers in parentheses above x-axis denote sample size. Double and triple asterisks denote statistically significant pairwise comparisons, with p < 0.010 and p < 0.001, respectively. See Tables S2–S4 for abbreviations.

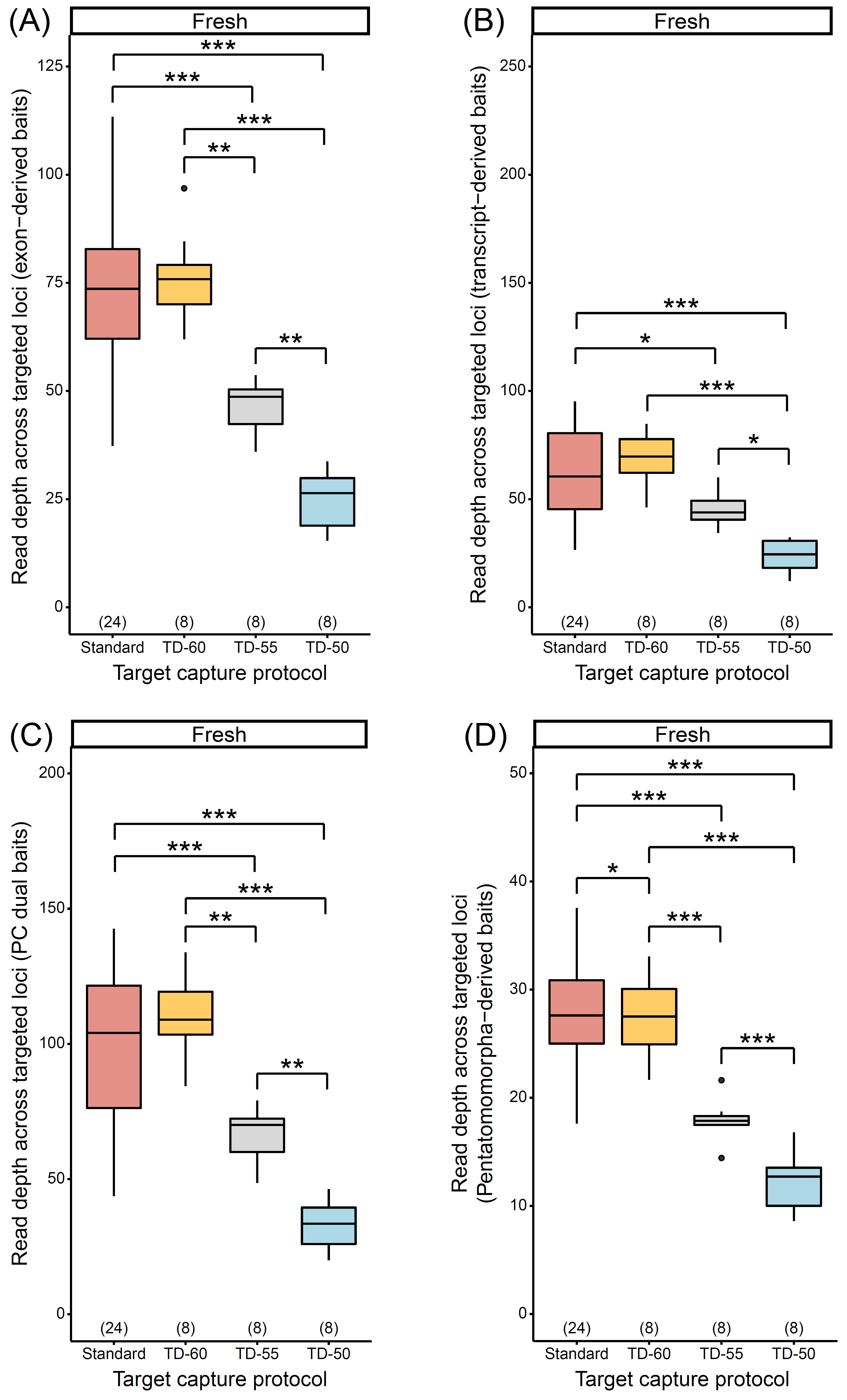

Figure S13. When controlling for sequencing depth, effects of target capture protocols on the read depth for loci targeted by (A) exon-derived baits, (B) transcript-derived baits, (C) PC dual baits, and (D) Pentatomomorpha-derived baits for samples preserved fresh. Numbers in parentheses above x-axis denote sample size. Single, double, and triple asterisks denote statistically significant pairwise comparisons, with p < 0.050, p < 0.010, and p < 0.001, respectively. See Tables S2–S4 for abbreviations.

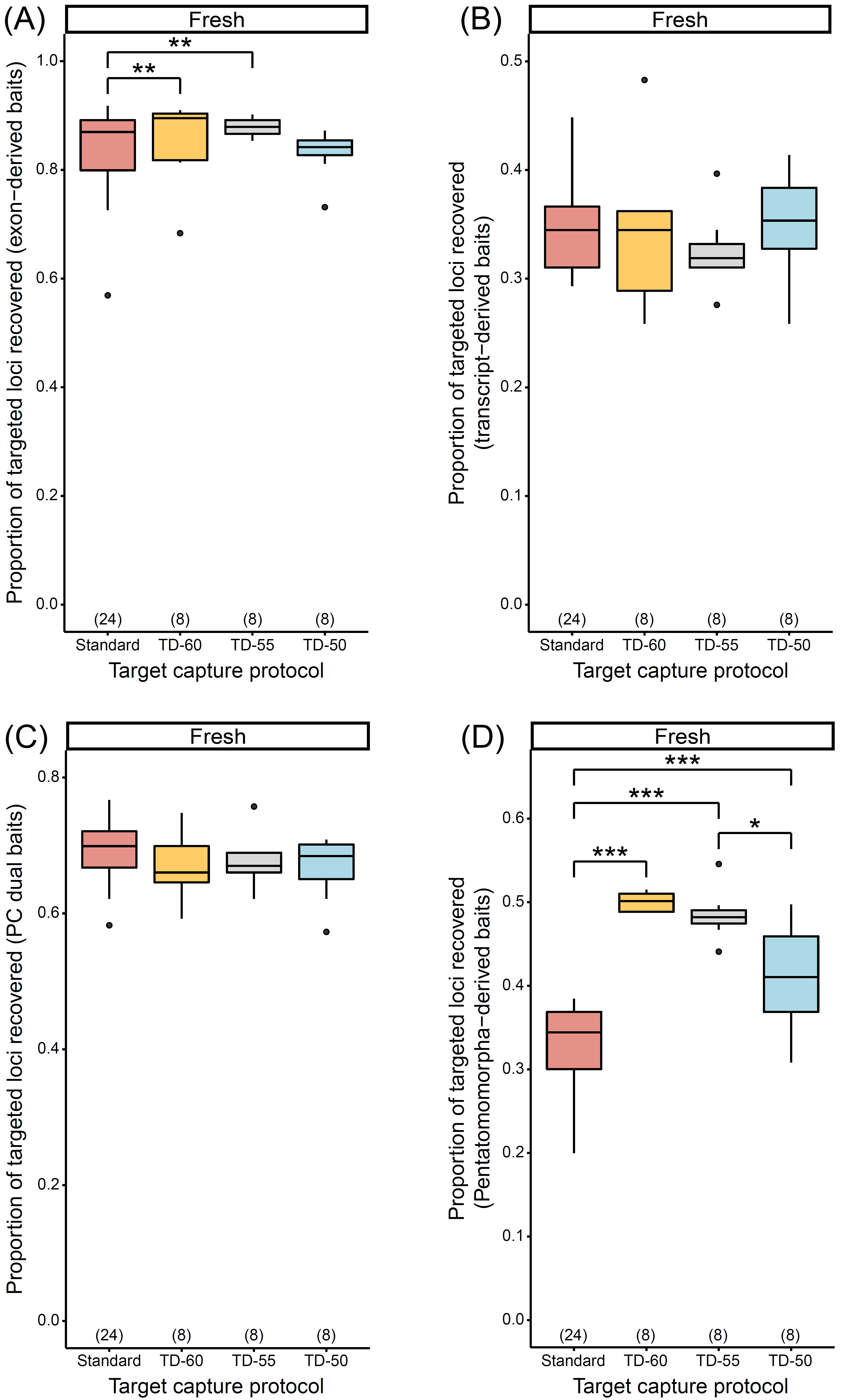

Figure S14. When controlling for sequencing depth, effects of target capture protocols on the proportion of loci captured by (A) exon-derived baits, (B) transcript-derived baits, (C) PC dual baits, and (D) Pentatomomorpha-derived baits for samples preserved fresh. Numbers in parentheses above x-axis denote sample size. Single, double, and triple asterisks denote statistically significant pairwise comparisons, with p < 0.050, p < 0.010, and p < 0.001, respectively. See Tables S2–S4 for abbreviations.

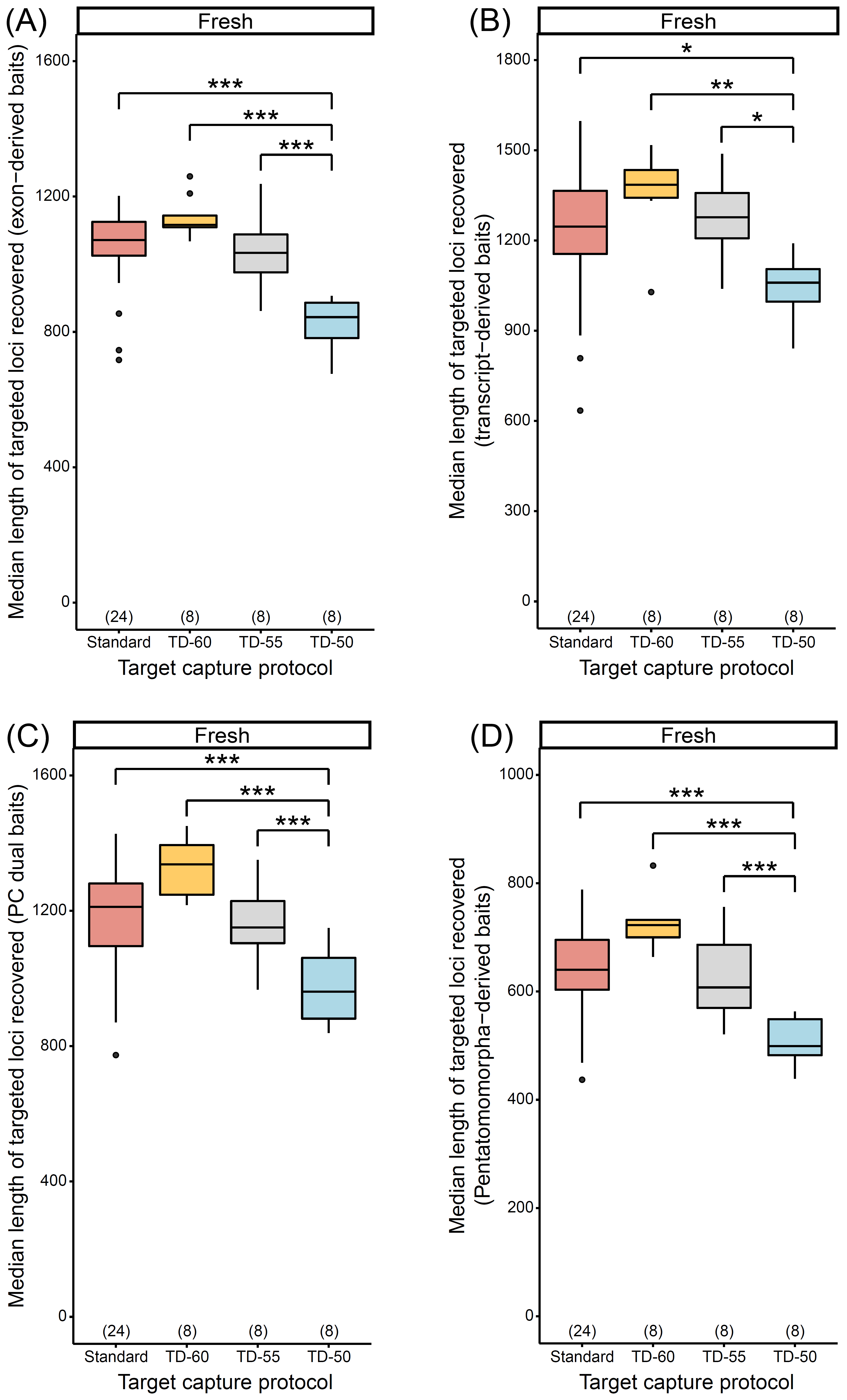

Figure S15. When controlling for sequencing depth, effects of target capture protocols on the median lengths of loci captured by (A) exon-derived baits, (B) transcript-derived baits, (C) PC dual baits, and (D) Pentatomomorpha-derived baits for samples preserved fresh. Numbers in parentheses above x-axis denote sample size. Single, double, and triple asterisks denote statistically significant pairwise comparisons, with p < 0.050, p < 0.010, and p < 0.001, respectively. See Tables S2–S4 for abbreviations.

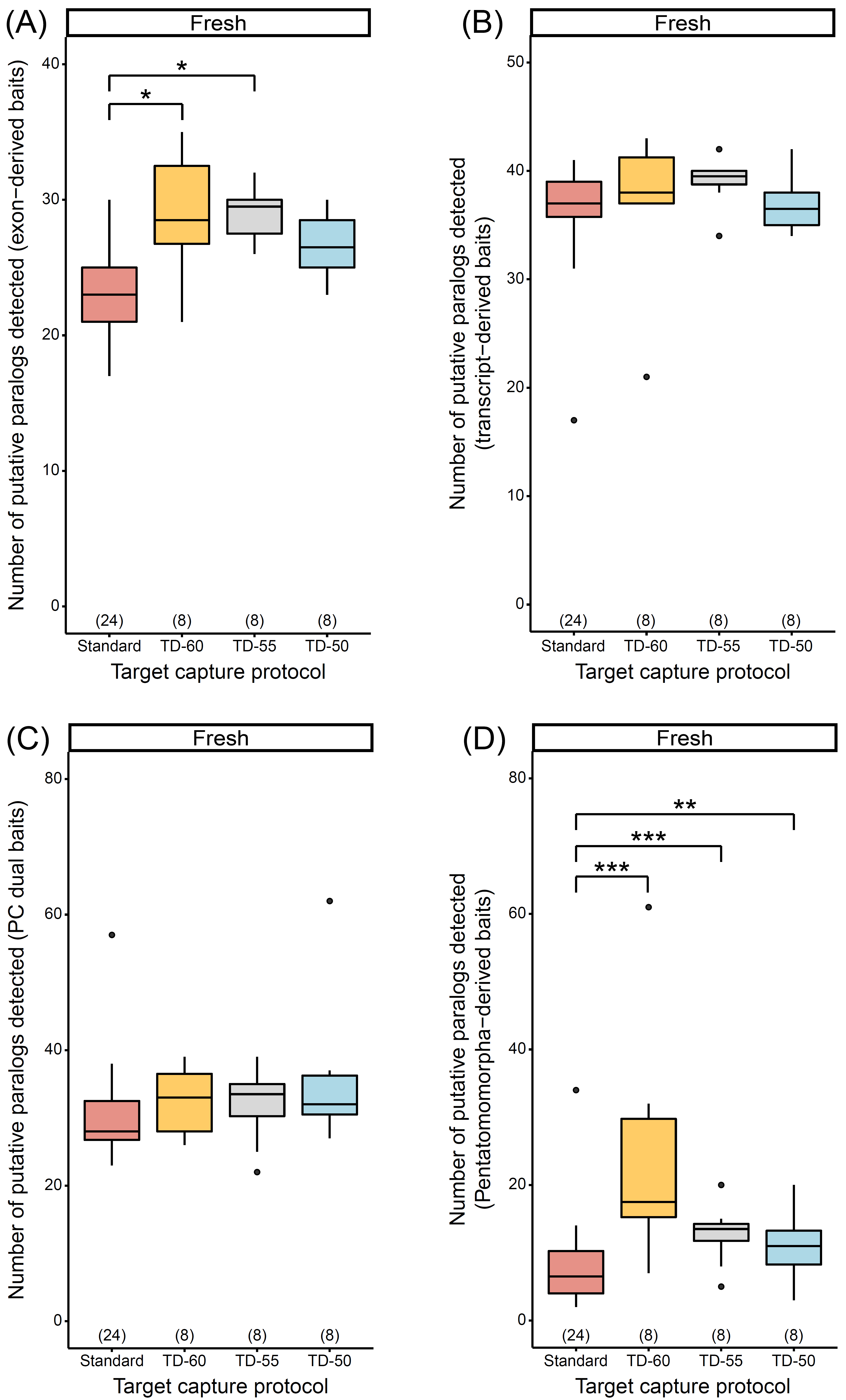

Figure S16. When controlling for sequencing depth, effects of target capture protocols on the number of putative paralogs of loci captured by (A) exon-derived baits, (B) transcript-derived baits, (C) PC dual baits, and (D) Pentatomomorpha-derived baits for samples preserved fresh. Numbers in parentheses above x-axis denote sample size. Single, double, and triple asterisks denote statistically significant pairwise comparisons, with p < 0.050, p < 0.010, and p < 0.001, respectively. See Tables S2–S4 for abbreviations.

Dallas, W. S. (1855). *List of the Specimens of Hemipterous Insects in the Collection of the British Museum. Part 2*. Trustees of the British Museum, London.

De Geer, C. (1773). *Mémoires pour server à l’Histoire des Insectes*. *Tome 3*. Stockholm: P. Hosselberg.

Emberts, Z., St. Mary, C. M., Howard, C. C., Forthman, M., Bateman, P. W., Somjee, U., Hwang, W. S., Li, D., Kimball, R. T., & Miller, C. W. (2020). The evolution of autotomy in leaf-footed bugs. *Evolution*, *74*(5), 897–910.

Fabricius, J. C. (1781). *Species insectorum exhibentes eorum differentias specificas, synonyma auctorum, loca natali, metamorphosin adjectis observationibus, descriptionibus*. Hamburgi et Kilonii, Bohnii.

Faircloth, B. C. (2017). Identifying conserved genomic elements and designing universal bait sets to enrich them. *Methods in Ecology and Evolution*, *8*(9), 1103–1112.

Forthman, M., Miller, C. W., & Kimball, R. T. (2019). Phylogenomic analysis suggests Coreidae and Alydidae (Hemiptera: Heteroptera) are not monophyletic. *Zoologica Scripta*, *48*(4), 520–534.

Forthman, M., Miller, C. W., & Kimball, R. T. (2020). Phylogenomics of the leaf-footed bug subfamily Coreinae (Hemiptera: Coreidae). *Insect Systematics and Diversity*, *4*, 2.

Glenn, T. C., Nilsen, R. A., Kieran, T. J., Sanders, J. G., Bayona-Vásquez, N. J., Finger, J. W., Pierson, T. W., Bentley, K. E., Hoffberg, S. L., Louha, S., Garcia-De León, F. J., del Rio Portilla, M. A., Reed, K. D., Anderson, J. L., Meece, J. K., Aggery, S. E., Rekaya, R., Alabady, M., Bélanger, M., Winker, K., & Faircloth, B. C. (2019). Adapterama I: universal stubs and primers for 384 unique dual-indexed or 147,456 combinatorially-indexed Illumina libraries (iTru & iNext). *PeerJ*, *7*, 049114.

Gomes, D. G. (2022). Should I use fixed effects or random effects when I have fewer than five levels of a grouping factor in a mixed-effects model?. *PeerJ*, *10*, e12794.

Grabherr, M. G., Haas, B. J., Yassour, M., Levin, J. Z., Thompson, D. A., Amit, I., Adiconis, X., Fan, L., Raychowdhury, R., Zeng, Q., Chen, Z., Mauceli, E., Hacohen, N., Gnirke, A., Rhind, N., di Palma, F., Birren, B. W., Nusbaum, C., Lindblad-Toh, K., Friedman, N., & Regev, A. (2011). Full-length transcriptome assembly from RNA-Seq data without a reference genome. *Nature Biotechnology*, *29*(7), 644–652.

Harrison, X. A. (2015). A comparison of observation-level random effect and Beta-Binomial models for modelling overdispersion in Binomial data in ecology & evolution. *PeerJ*, *3*, e1114.

Herrich-Schäffer, G. A. W. (1847). *Die wanzenartigen Insecten*. *8*. Nurnberg: C. H. Zeh.

Hothorn, T., Bretz, F., & Westfall, P. (2008). Simultaneous inference in general parametric models. *Biometrical Journal*, *50*(3), 246–363.

Johnson, K. P., Dietrich, C. H., Friedrich, F., Beutel, R. G., Wipfler, B., Peters, R. S., Allen, J. M., Petersen, M., Donath, A., Walden, K. K. O., Kozlov, A. M., Podsiadlowski, L., Mayer, C., Meusemann, K., Vasilikopoulos, A., Waterhouse, R. M., Cameron, S. L., Weirauch, C., Swanson, D. R., Percy, D. M., Hardy, N. B., Terry, I., Liu, S., Zhou, X., Misof, B., Robertson, H. M., & Yoshizawa, K. (2018). Phylogenomics and the evolution of hemipteroid insects. *Proceedings of the National Academy of Sciences*, *115*(50), 12775–12780.

Knyshov, A., Gordon, E. R. L., & Weirauch, C. (2019). Cost-efficient high throughput capture of museum arthropod specimen DNA using PCR-generated baits. *Methods in Ecology and Evolution,* *10*(6), 841–852.

Marçais, G., Yorke, J. A., & Zimin, A. (2015). QuorUM: an error corrector for Illumina reads. *PLoS ONE*, *10*(6), e0130821.

Portik, D. M., Smith, L. L., & Bi, K. (2016). An evaluation of transcriptome-based exon capture for frog phylogenomics across multiple scales of divergence (Class: Amphibia, Order: Anura). *Molecular Ecology Resources*, *16*(5), 1069–1083.

Quinlan, A. R., & Hall, I. M. (2010). BEDTools: a flexible suite of utilities for comparing genomic features. *Bioinformatics*, *26*(6), 841–842.

R Core Team. (2022). *R: a language and environment for statistical computing. R Foundation for Statistical Computing*. R Foundation for Statistical Computing: Vienna, Austria. https://www.R-project.org/.

Say, T. (1825). Descriptions of new hemipterous insects collected in the expedition to the Rocky Mountains, performed by order of Mr. Calhoun, Secretary of War, under command of Major Long. *Journal of the Academy of Natural Sciences of Philadelphia*, *4*, 7–345.

Schmieder, R., & Edwards, R. (2011). Quality control and preprocessing of metagenomic datasets. *Bioinformatics*, *27*(6), 863–864.

Stål, C. (1855). Hemiptera från kafferlandet. *Forhandlingar Svenska Vetenskaps-Akademien Ofversigt*, *12*, 27–46.

Williams, D. A. (1982). Extra‐binomial variation in logistic linear models. *Journal of the Royal Statistical Society: Series C (Applied Statistics)*, *31*(2), 144–148.
